# Synthesis and Biological Evaluation of Nipecotic Acid Derivatives with Terminally Double-Substituted Allenic Spacers as mGAT4 Inhibitors

**DOI:** 10.1101/2025.02.15.638425

**Authors:** Maren Jung, Georg Höfner, Kinga Sałat, Anna Furgała-Wojas, Klaus T. Wanner

## Abstract

A series of nipecotic acid derivatives, featuring a four- and five-carbon atom allenic spacer connecting the nitrogen of the polar nipecotic acid head with up to two aromatic residues, has been synthesized. For the synthesis of the respective nipecotic acid derivatives displaying *N*-substituents consisting of carbon chains with a terminal allenic function with two, mostly aryl residues a Cu^I^-catalyzed cross-coupling of diaryldiazomethanes or *N*-tosylhydrazones with nipecotic acid derived terminal alkynes served as a key step. Upon characterization of these compounds regarding their inhibitory potency at mGAT1-4 and binding affinity for mGAT1, a highly potent mGAT4 inhibitor has been identified, which is defined as nipecotic acid derivative with a four-carbon atom allenic spacer terminally carrying two 4-chlorophenyl residues. The (*S*)-enantiomer of this compound, (*S*)-1-[4,4-bis(4-chlorophenyl)buta-2,3-dien-1-yl]piperidine-3-carboxylic acid [(*S*)-**8d**, DDPM-3960], displays potencies in the higher nanomolar range at mGAT4 and its human equivalent hGAT-3 with pIC_50_ values of 6.59 ± 0.01 and 6.49 ± 0.10, respectively, which are significantly higher than that of the well-known mGAT4 inhibitor (*S*)-SNAP-5114. *In vivo* evaluation of this compound revealed its significant anticonvulsant activity in several mouse models of chemically- and electrically-induced seizures. In addition to this, anxiolytic-like properties of DDPM-3960 were shown. These beneficial biological effects observed in mice were not accompanied by any serious motor deficits, making DDPM-3960 an interesting lead structure for further development.

## Introduction

γ-Aminobutyric acid (GABA) (**1**, Figure 1), the major inhibitory neurotransmitter in the mammalian central nervous system (CNS), plays a central role in normal brain function. The uptake of GABA into neurons and glial cells mediated by GABA transporters (GATs) is an important part of the GABAergic system. Targeting the individual stages of GABA signaling to overcome a dysregulation of GABA neurotransmission has become a valuable strategy for the potential therapy of related diseases such as epilepsy, Parkinson’s disease, Huntington’s disease, schizophrenia, and Alzheimer’s disease.^1^ Through inhibition of the enzymatic breakdown of GABA, inhibition of the GABA transport proteins, or allosteric modulation of the GABA_A_ receptors, an increased GABAergic activity might be achieved.^2, 3, 4^ The antiepileptic drug vigabatrin, for instance, improves the GABA level by acting as a suicide inhibitor of GABA transaminase, an enzyme, that is responsible for the metabolic degradation of GABA. The enhancement of GABA neurotransmission through allosteric modulation of GABA_A_ receptors can be achieved by the use of benzodiazepines, which are amongst the most commonly used GABAergic drugs today. GABA transporters (GATs) mediate the transport of synaptically released GABA from the extracellular to the intracellular side of glial and neuronal cells. Inhibition of these transporters leads to enhanced GABA signaling as a consequence of increased extracellular GABA levels. This has been shown to be effective in the treatment of seizures in epileptic disorders and is further considered a promising approach for the treatment of diseases such as depression, anxiety, pain, Alzheimeŕs disease and sleep disorders.^5^ There are four different GABA transporter subtypes belonging to the solute carrier 6 family (SLC6), which use the co-transport of sodium ions as driving force for the translocation of their substrates against chemical gradients.^6^ The subtypes cloned from mouse brain are termed as mGAT1, mGAT2, mGAT3, and mGAT4,^7^ whereas the human-derived GABA transporters are named as hGAT-1, hBGT-1, hGAT-2, and hGAT-3, respectively. This nomenclature is also adopted by the Human Genome Organization (HUGO).^8^ mGAT1 and mGAT4 are almost exclusively expressed around the synaptic cleft, whereat mGAT1 is primarily located in presynaptic neuronal membranes, and mGAT4 is mainly located in astrocytes. mGAT1 is considered to be the most important transporter subtype for neuronal GABA uptake, which makes it an interesting drug target. Nipecotic acid (*rac*-**2**, Figure 1), a cyclic GABA analogue, has subsequently been used as lead structure in the efforts to synthesize potent GAT inhibitors. Among them, tiagabine (**3**, Figure 1) is the only currently approved antiepileptic drug that addresses mGAT1 with high affinity and subtype selectivity. In this context, it deserves mentioning, that a high-resolution structure of hGAT1 became available only recently, which can be expected to strongly support the development of further and even more potent and selective inhibitors of this target in the future. This structure determined by cryo-electron microscopy, disclosed by the Gati group in 2022, shows the GABA transporter hGAT-1 in complex with tiagabine in the inward-open conformation.^9^ In addition, in the same year also the cryo-EM structure of rGAT1 with bound GABA has been published.^10^

**Figure 1.**
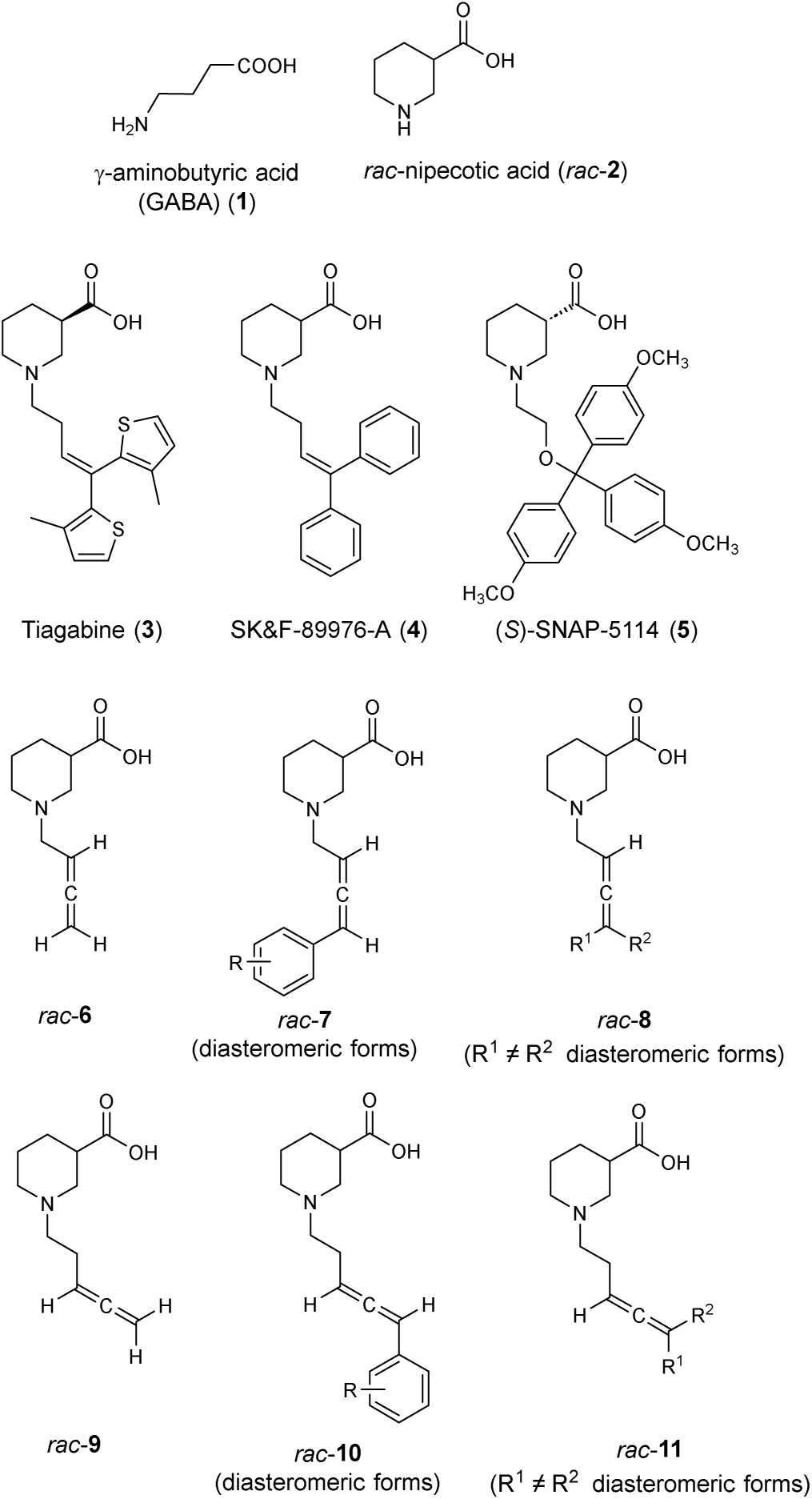
Structures of GABA (**1**), selected GAT inhibitors (**2-5**) and nipecotic acid derivatives with allenic spacer (rac-**6**-rac-**11**).

As tiagabine (**3**) has the disadvantage of occasionally observed adverse effects, which seem to be inherently coupled with the function of other GAT subtypes, in particular mGAT4, have gained increasing research interest.^11, 12, 13^ Although plenty of highly potent and subtype selective inhibitors are available for mGAT1, the lack of similar potent inhibitors for mGAT4 still retards its pharmacological elucidation. The first prototypic mGAT4 inhibitor with moderate potency and selectivity is represented by (*S*)-SNAP-5114 (**5**, Figure 1).^14^ However, the application of (*S*)-SNAP-5114 (**5**) *in vivo* is limited due to its low brain uptake and modest chemical stability.^15, 16^ In addition, its selectivity is not sufficiently pronounced to exclude effects on other GABA transporter subtypes. Still, it could be demonstrated, that (*S*)-SNAP-5114 (**5**) is able to exert anticonvulsant effects in the Frings audiogenic seizure-susceptible mouse model.^17^ The potential of mGAT4 inhibitors has become further evident from a recent report on a pharmacological study of a mGAT4 knockout mouse model. Based on their results the authors conclude that mGAT4 could be a target for the treatment of pain and several psychiatric disorders such as depression, anxiety, schizophrenia and autism:^18^ Hence, the development of more potent and selective mGAT4 inhibitors appears to be of great interest to further deepen the insights in the physiological effects and therapeutic potential of inhibitors of this transporter.

Besides, there is still a need for mGAT1 inhibitors with more favorable pharmacokinetic properties than tiagabine (**3**), which is used as an add-on therapeutic agent for epilepsy,^19, 20^ and that might also possess less adverse effects, such as dizziness, asthenia, nervousness, tremor and diarrhea than **3**.^21^

Owing to the unique structural features, related to the presence of two in perpendicular planes located π bonds, allenes are important and useful building blocks in organic synthesis.^22^ Many natural products with interesting biological activity, containing an allene moiety, have been identified.^23, 24^ The interest in this scaffold is constantly growing, due to its intrinsic three-carbon axial chirality, less-hindered linear structure and high substituent-loading ability. Recently, a compound class characterized by a four-carbon atom allenic spacer connecting the nitrogen of the nipecotic acid subunit with an aromatic residue (*rac*-**7**, Figure 1) has been synthesized in our group and investigated regarding its biological activities at mGAT1-4. As a result, new mGAT1 inhibitors with good inhibitory potencies and subtype selectivities have been identified among these nipecotic acid derivatives with a terminally mono-substituted allenic spacer.^25^ In continuation of this study, we now aimed at the synthesis and biological characterization of related compounds. A first set should comprise compounds, in which the allenic spacer in *rac*-**7** has been extended by one carbon to an allenic five-carbon subunit with one terminal aryl substituent as represented by formula *rac*-**10** (Figure 1). This set should also include the terminally unsubstituted nipecotic acid derivative *rac*-**9**. In case of compounds *rac*-**7** the nipecotic acid derivative *rac*-**6** with an unsubstituted *N*-buta-2,3-dien-1-yl moiety had served as a reference compound in the biological studies. Compound *rac*-**9** exhibiting a penta-3,4-dien-1-yl residue was thought to analogously fulfill the same purpose for compounds *rac*-**10**.

Furthermore, we aimed at nipecotic acid derivatives with an allenic four- and five-carbon atom spacer with two terminal substituents, i.e. *rac*-**8** and *rac*-**11** (Figure 1). Regarding the two latter classes of compounds, it was of interest to introduce two lipophilic aryl residues, like those present in SK&F-89976 (**4**, Figure 1), which are known to be favorable for enhanced potency at and selectivity for mGAT1.

When the synthesis of *rac*-**9** has been attempted in close analogy to that of *rac*-**6**, already reported before,^25, 26^ this compound, *rac*-**9**, turned out to undergo side reactions, when conditions required for the final carboxylic acid ester hydrolysis and subsequent workup were employed (see SI). As this instability had not been observed for *rac*-**6**, it appears likely to be due to the extended length of the allenic *N*-substituent in *rac*-**9**. Whatsoever, the chemical stability of compound *rac*-**9** was too low for *rac*-**9** being used in biological studies.

Furthermore, the synthesis of nipecotic acid derivatives *rac*-**10** with an allenic five-carbon atom spacer and a terminal phenyl (R = H) and biphenyl group (R = *ortho*- phenyl) has been attempted. To this end, a method published by Wang et al. has been used, that allows the direct transformation of terminal alkynes into the respective allene derivatives upon reaction with appropriate *N*-tosylhydrazones under Cu^I^ catalysis.^27, 28^

Though the preparation of the carboxylic acid ethyl esters of the target compounds *rac*- **10** (R = H, R = *ortho*-phenyl) could be accomplished this way, attempts to transform these carboxylic acid esters into the free carboxylic acid derivatives *rac*-**10** (R = H, R = *ortho*-phenyl) by acidic or basic hydrolysis despite variations of the reaction conditions failed, the reactions yielding only decomposition products (see SI).

Taking the results observed for parent compound *rac*-**9** and *rac*-**10** (R = H, R = *ortho*- phenyl, Figure 1) together, nipecotic acid derivatives with a five-carbon atom allenic spacer appear to be less stable than their analogues with a four-carbon atom allenic unit.

Due to these disappointing results, the synthesis of nipecotic acid derivatives with an allenic five-carbon residue and a single terminal substituent as represented by formula *rac*-**10** was not further pursued, instead the synthesis of compounds *rac*-**8** and *rac*-**11** with two terminal residues attached to the buta-2,3-dien-1-yl and the penta-3,4-dien-1- yl spacer, respectively, were taken into focus.

## Results and Discussion

### Chemistry

For the synthesis of the desired nipecotic acid derivatives with terminally double substituted four- and five-carbon allenic spacers we intended to follow a synthetic approach related to the ATA reaction (allenylation of terminal alkynes), that we had used before with great success for the preparation of the terminally un- and monosubstituted allenic derivatives *rac*-**6** and *rac*-**7** (Figure 1).^25, 26^

To this end, envisaging a stepwise process, at first propargylic amines derived from *rac*-**12** and *rac*-**15** were required, which we thought to synthesize via the ketone-alkyne-amine coupling reactions (KA^2^ coupling), in which ketones instead of aldehydes together with the respective amine are reacted with the terminal alkyne. According to literature harsher reaction conditions, i.e. higher temperatures are required,^[29]^ for the coupling of terminal alkynes with ketones as compared to aldehydes. Unfortunately, in our case all attempts to generate ketone derived propargylic amine intermediates for the allenylation of terminal alkynes such as *rac*-**12** failed. Therefore, an alternative synthesis route had to be found for the preparation of the target compounds. The copper catalyzed coupling of terminal alkynes with diaryldiazomethanes via a Cu^I^ carbene migratory insertion mechanism appeared as a promising method, since it should rapidly lead to substituted allenes from simple and readily available starting materials.^30^

Hence, this method was used in the attempts to synthesize the desired nipecotic acid derivatives with a terminally double-substituted allenic four- and five-carbon atom spacer (Scheme 1). The required diaryldiazomethanes **13a**-**f** were synthesized in a general procedure from the corresponding ketones, which were transformed into their hydrazones and subsequently oxidized by MnO_2_ (see Experimental Section). Cu^I^-catalyzed coupling of diaryldiazomethanes **13a**-**e** with terminal alkynes *rac*-**12** and *rac*- **15**, respectively, in the presence of *i*-Pr_2_NH provided finally the desired 1,3,3- trisubstituted allenes *rac*-**14a**-**e** and *rac*-**16a**-**e** in yields of up to 97%. When diazomethane **13f** exhibiting two different aryl residues was employed with *rac*-**12** and *rac*-**15** in this reaction, accordingly, *rac*-(*3R*,*R_a_*)-**14f**/*rac*-(*3R*,*S_a_*)-**14f** and *rac*-(*3R*,*R_a_*)- **16f**/*rac*-(*3R*,*S_a_*)-**16f** as ∼ 1:1 mixture of racemic diastereomers were obtained [yield: *rac*-(*3R*,*R_a_*)-**14f**/*rac*-(*3R*,*S_a_*)-**14f**, 94%; *rac*-(*3R*,*R_a_*)-**16f**/*rac*-(*3R*,*S_a_*)-**16f**, 88%]. Subsequent hydrolysis under basic conditions delivered the corresponding free amino acids rac-**8a**-**d**, *rac*-(*3R*,*R_a_*)-**8f**/*rac*-(*3R*,*S_a_*)-**8f**, *rac*-**11a-d** and *rac*-(*3R*,*R_a_*)-**11f**/*rac*- (*3R*,*S_a_*)-**11f** in yields up to 52% (Scheme 1). The nipecotic acid esters *rac*-**14e** and *rac*- **16e** were found to undergo rapid decomposition, wherefore no attempts for the synthesis of the corresponding free nipecotic acid derivatives rac-**8e** and *rac*-**11e** were undertaken. Furthermore, the enantiopure compounds (*R*)-**8d** and (*S*)-**8d** have been synthesized in an analogous manner via (*R*)-**14d** and (*S*)-**14d** applying terminal alkynes (*R*)-**12** and (*S*)-**12**, respectively, as starting material [yields: (*R*)-**8d**, 51%; (*S*)- **8d**, 68%].

Nipecotic acid derivatives *rac*-**11a-d** and *rac*-(*3R*,*R_a_*)-**11f**/*rac*-(*3R*,*S_a_*)-**11f** with an allenic five-carbon atom spacer, were found to be prone to side reactions during RP-MPLC purification and subsequent freeze drying. Without recognizable differences in reaction performance the decomposition tendency is varying within this series of nipecotic acid derivatives, whereby these compounds distinguish from their more stable analogues [*rac*-**8a**-**d**, *rac*-(*3R*,*R_a_*)-**8f**/*rac*-(*3R*,*S_a_*)-**8f**] only by the length of the allenic spacer. The substituents on the two phenyl residues were observed to have an influence on the decomposition tendency.

**Scheme 1.**
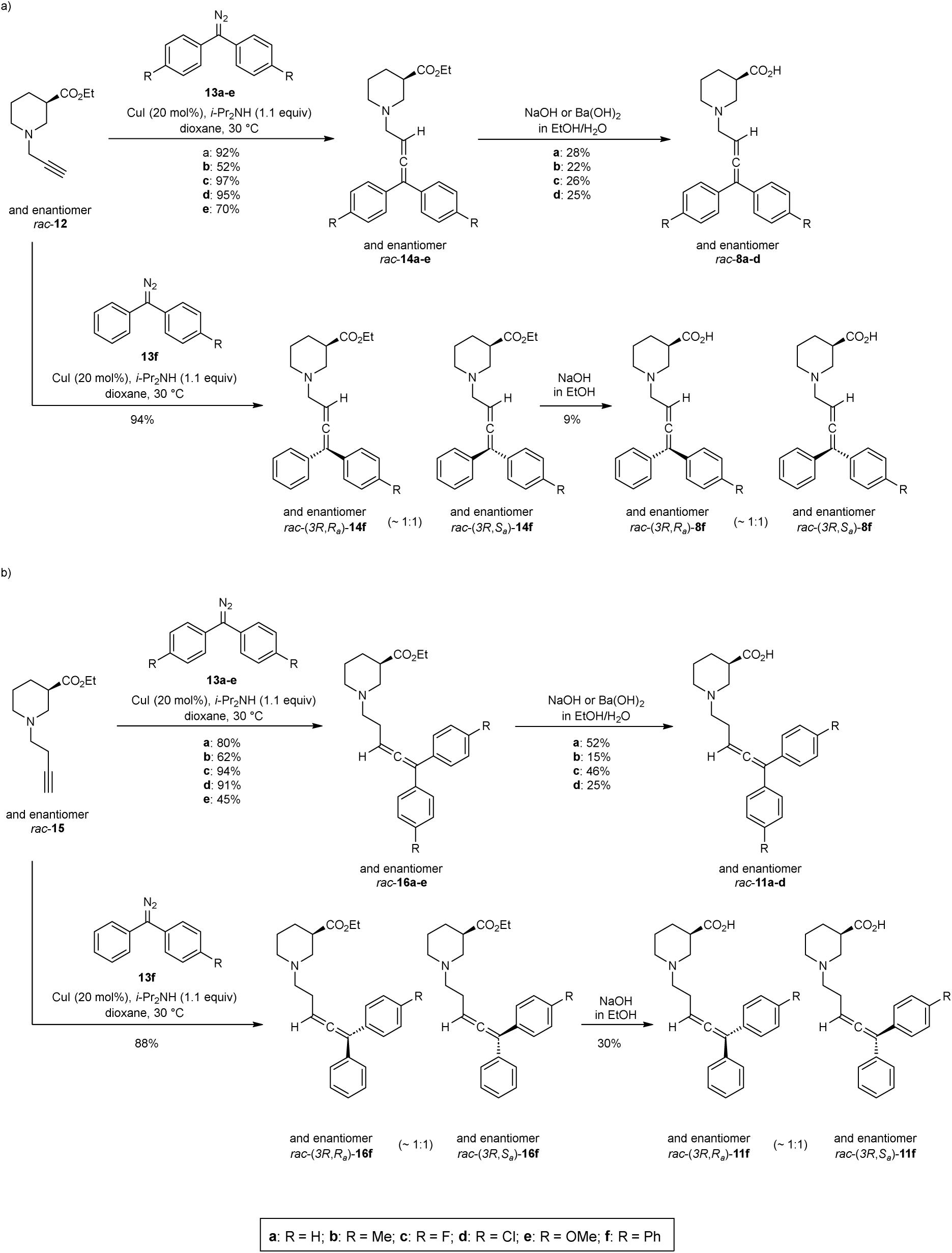
Synthesis of nipecotic acid derivatives with terminally double-substituted allenic spacer via Cu^I^- catalyzed cross-coupling of diaryldiazomethanes **13a-f** and terminal alkynes (rac-**12)** and (rac-**15**).

For compound *rac*-**11d** for example, complete decomposition was observed following the common work-up procedure. By means of ^1^H NMR of the isolated side product **17** (see supporting information), the most prominent side reaction appears to be a reversible cyclization reaction (Scheme 2). As side product **17** exists as a mixture of racemic diastereomers, NMR signals could not be clearly assigned, wherefore the proposed structure is not completely confirmed. In literature, related allene involved 6- endo-trig cyclization reactions have already been described.^31^ In contrast to *rac*-**11d**, exhibiting a *p*-chloro substituent at both phenyl groups, that had completely decomposed to putatively **17** after MPLC purification in a MeOH/H_2_O solvent mixture and subsequent lyophilization, no decomposition has been observed for nipecotic acid derivative *rac*-**11c**, bearing *p*-fluoro substituents. Fortunately, when repeating the synthesis of *rac*-**11d** under identical reaction conditions, except that the final MPLC purification step was omitted, compound *rac*-**11d** could be isolated in pure form after aqueous workup. Hence, the MPLC purification step is likely to be responsible for the decomposition, that had been observed before.

With compound *rac*-**11d** in hand, further experiments have been performed to investigate its tendency to decompose during the required incubation time in the applied incubation buffers for uptake (Tris-NaCl buffer) and binding assays (Krebs buffer). For this purpose, a 100 µM solution of *rac*-**11d** in buffer solution, containing 1% DMSO was stirred for 40 min, respectively 35 min, at 37 °C and was quenched by ether extraction. ^1^H NMR analysis of the extracted residue revealed that no decomposition had occurred and therefore results of the following biological studies can be assumed to result from the allenic compound *rac*-**11d**. As nipecotic acid derivative *rac*-**11d** displayed the highest tendency for decomposition in this study, it is reasonable to assume, that also the nipecotic acid derivatives *rac*-**11a**-**c** and *rac*- (*3R*,*R_a_*)-**11f**/*rac*-(*3R*,*S_a_*)-**11f**, as the apparently more stable analogues, did not undergo decomposition during the biological testing.

**Scheme 2.**
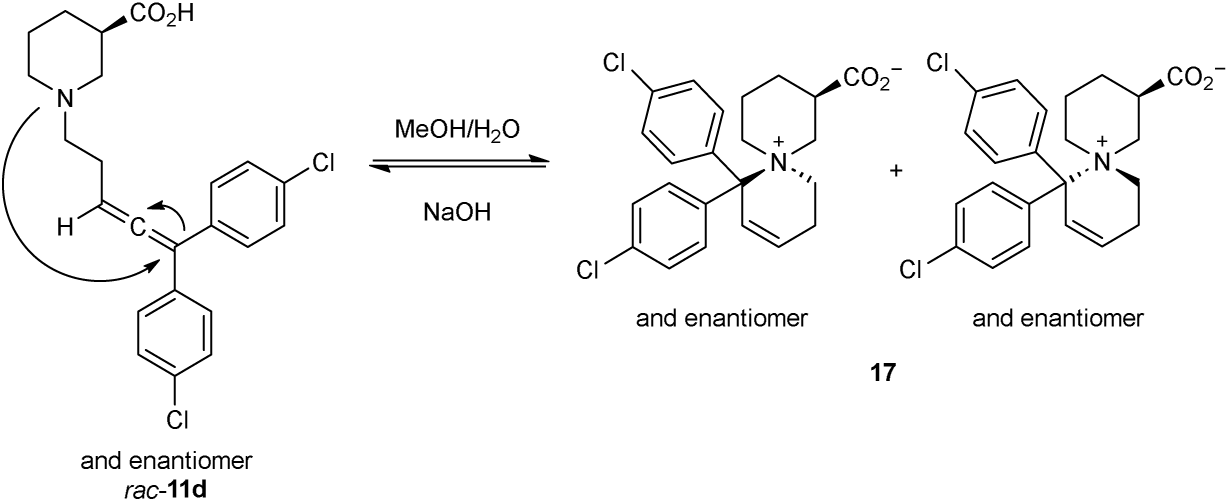
Proposed reversible 6-endo cyclization of nipecotic acid derivative rac-**11d** to side product **17**.

When applying ketones other than diaryl-ketones, the coupling procedure via *N*- tosylhydrazones reported by Wang et al. for this type of substrates, i.e. of aryl-alkyl ketones and aldehydes was used (Scheme 3).^27, 28^ Reaction of the *N*-tosylhydrazone of 6-methoxy-1-tetralone **18** with terminal alkyne *rac*-**12** under copper catalysis delivered the corresponding mixture of nipecotic acid ester derivatives *rac*-(*3R*,*R_a_*)-**14g** and *rac*-(*3R*,*S_a_*)-**14g** (ratio ∼ 1: 1) in 60% yield. Basic hydrolysis provided the desired target compounds, the free carboxylic acids derivatives (*3R*,*R_a_*)-**8g** and *rac*-(*3R*,*S_a_*)-**8g** (ratio ∼ 1: 1) in 61 % yield.

**Scheme 3.**
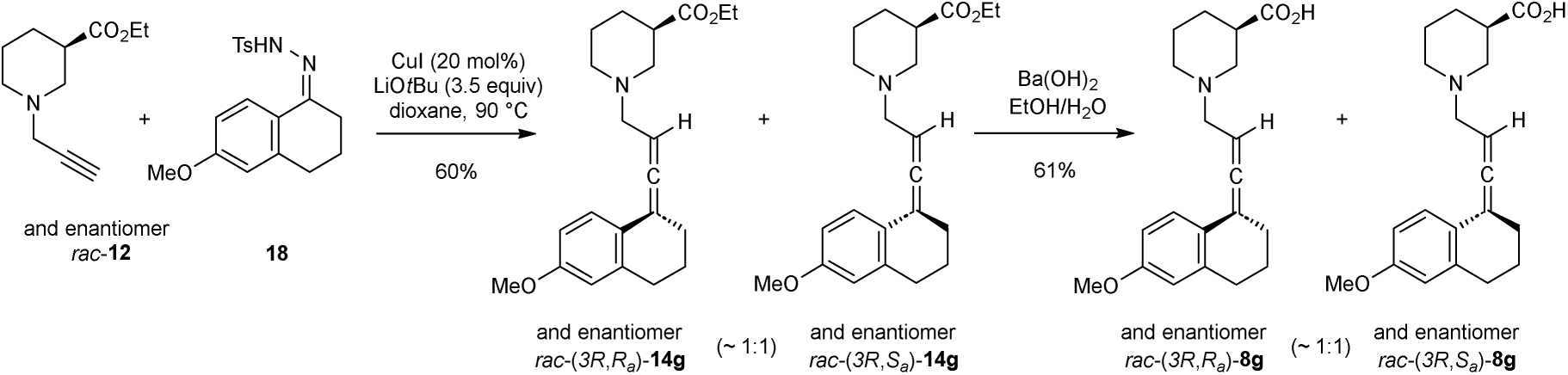
Synthesis of nipecotic acid derivatives [rac-(3R,R_a_)-**8g**] and [rac-(3R,S_a_)-**8g**] with terminally double-substituted four-carbon atom allenic spacer as ∼ 1:1 mixture of racemic diastereomers by Cu^I^- catalyzed reaction of rac-**12** with N-tosylhydrazone **18** and subsequent ester hydrolysis.

### Biological Evaluations

#### *In vitro* part

For synthesized nipecotic acid derivatives *rac*-**8a-d**, *rac*-(*3R*,*R_a_*)-**8f**-**g**/*rac*-(*3R*,*S_a_*)-**8f**-**g**, *rac*-**11a**-**d** and *rac*-(*3R*,*R_a_*)-**11f**/*rac*-(*3R*,*S_a_*)-**11f** binding affinities for mGAT1 (p*K*_i_ values) were determined in MS Binding Assays with NO711 as native MS marker.^32^ In addition, their functional activity was characterized as pIC_50_ values in [^3^H]GABA-Uptake-Assays with HEK293 cells stably expressing the individual mouse GABA transporters mGAT1-mGAT4.^33^ Compounds *rac*-(*3R*,*R_a_*)-**8f**-**g**/*rac*-(*3R*,*S_a_*)-**8f**-**g** and *rac*-(*3R*,*R_a_*)-**11f**/*rac*-(*3R*,*S_a_*)-**11f** have been tested as ∼ 1:1 mixtures of racemic diastereomers as which they had been formed. The biological results for each of the stereoisomers, i.e. enantiomers and diastereomers, will most likely differ from the results of the mixture. Hence, the derived structure-activity relationships for these compounds should be regarded as estimates. The data of the biological tests are listed in Table 1 supplemented with those of parent compound *rac*-**6** bearing an allenic four-carbon atom residue attached to the nitrogen of the nipecotic acid serving as spacer in the more elaborate compounds, that had been determined in a former study.^25^

**Table 1:**
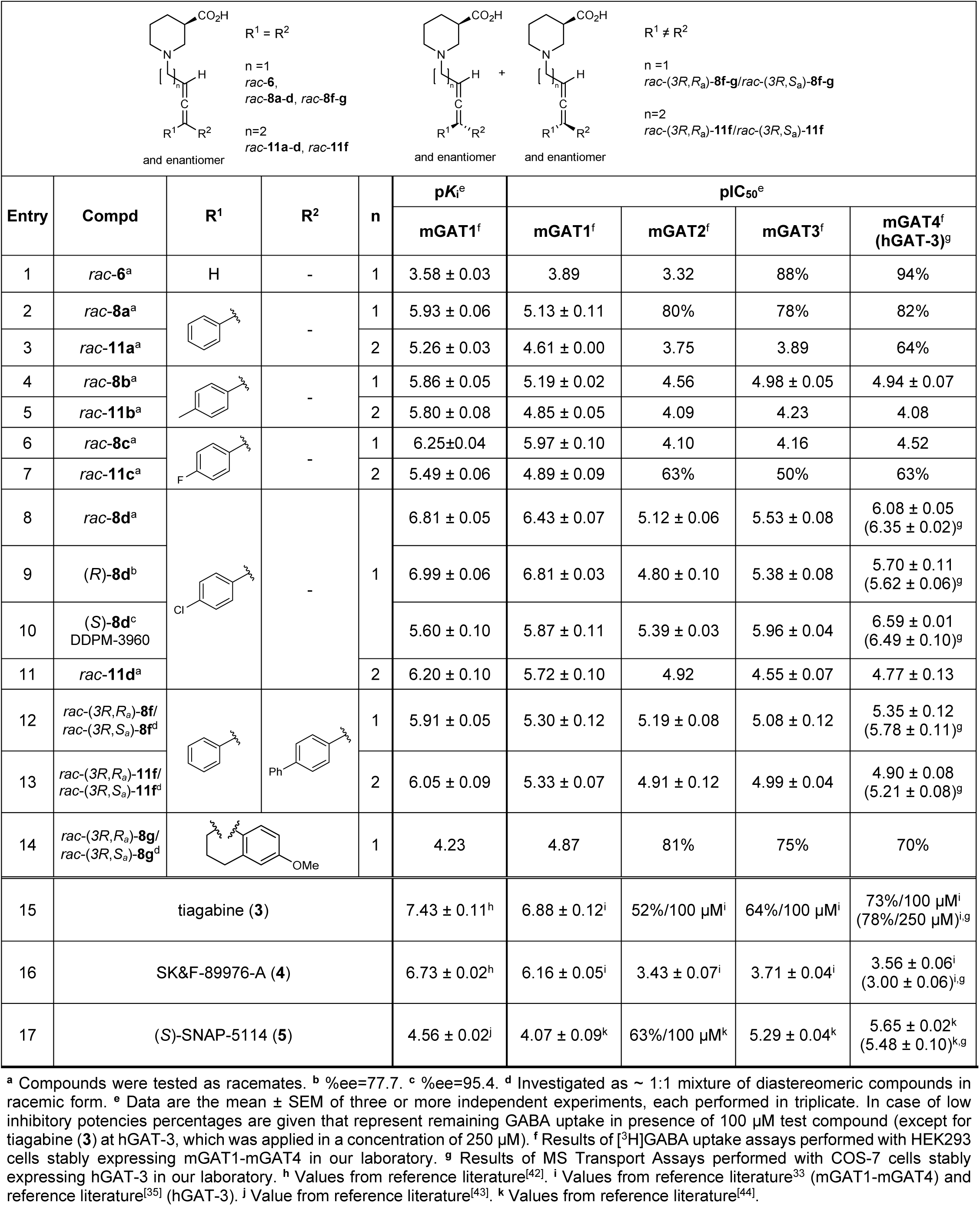
Nipecotic acid derived GAT inhibitors containing terminally double-substituted allenic spacers and their inhibitory potencies at mGAT1-mGAT4 (hGAT-3) and binding affinities for mGAT1.

| $R^1 = R^2$<br>$n=1$<br>*rac-6*,<br>*rac-8a-d*, *rac-8f-g*<br>$n=2$<br>*rac-11a-d*, *rac-11f*<br>and enantiomer<br>$R^1 \neq R^2$<br>$n=1$<br>*rac-(3R,R_a)-8f-g/rac-(3R,S_a)-8f-g*<br>$n=2$<br>*rac-(3R,R_a)-11f/rac-(3R,S_a)-11f*<br>and enantiomer |  |  |  |  |  |  |  |  |  |
| --- | --- | --- | --- | --- | --- | --- | --- | --- | --- |
| Entry | Compd | R<sup>1</sup> | R<sup>2</sup> | n | pK<sub>i</sub><sup>e</sup> | pIC<sub>50</sub><sup>e</sup> |  |  |  |
|  |  |  |  |  | mGAT1<sup>f</sup> | mGAT1<sup>f</sup> | mGAT2<sup>f</sup> | mGAT3<sup>f</sup> | mGAT4<sup>f</sup><br>(hGAT-3)<sup>g</sup> |
| 1 | *rac-6*<sup>a</sup> | H | - | 1 | 3.58 ± 0.03 | 3.89 | 3.32 | 88% | 94% |
| 2 | *rac-8a*<sup>a</sup> |  | - | 1 | 5.93 ± 0.06 | 5.13 ± 0.11 | 80% | 78% | 82% |
| 3 | *rac-11a*<sup>a</sup> |  |  | 2 | 5.26 ± 0.03 | 4.61 ± 0.00 | 3.75 | 3.89 | 64% |
| 4 | *rac-8b*<sup>a</sup> |  | - | 1 | 5.86 ± 0.05 | 5.19 ± 0.02 | 4.56 | 4.98 ± 0.05 | 4.94 ± 0.07 |
| 5 | *rac-11b*<sup>a</sup> |  |  | 2 | 5.80 ± 0.08 | 4.85 ± 0.05 | 4.09 | 4.23 | 4.08 |
| 6 | *rac-8c*<sup>a</sup> |  | - | 1 | 6.25±0.04 | 5.97 ± 0.10 | 4.10 | 4.16 | 4.52 |
| 7 | *rac-11c*<sup>a</sup> |  |  | 2 | 5.49 ± 0.06 | 4.89 ± 0.09 | 63% | 50% | 63% |
| 8 | *rac-8d*<sup>a</sup> |  | - | 1 | 6.81 ± 0.05 | 6.43 ± 0.07 | 5.12 ± 0.06 | 5.53 ± 0.08 | 6.08 ± 0.05<br>(6.35 ± 0.02)<sup>g</sup> |
| 9 | (*R*)-*8d*<sup>b</sup> |  |  |  | 6.99 ± 0.06 | 6.81 ± 0.03 | 4.80 ± 0.10 | 5.38 ± 0.08 | 5.70 ± 0.11<br>(5.62 ± 0.06)<sup>g</sup> |
| 10 | (*S*)-*8d*<sup>c</sup><br>DDPM-3960 |  |  |  | 5.60 ± 0.10 | 5.87 ± 0.11 | 5.39 ± 0.03 | 5.96 ± 0.04 | 6.59 ± 0.01<br>(6.49 ± 0.10)<sup>g</sup> |
| 11 | *rac-11d*<sup>a</sup> |  |  | 2 | 6.20 ± 0.10 | 5.72 ± 0.10 | 4.92 | 4.55 ± 0.07 | 4.77 ± 0.13 |
| 12 | *rac-(3R,R<sub>a</sub>)-8f/*<br>*rac-(3R,S<sub>a</sub>)-8f*<sup>d</sup> |  |  | 1 | 5.91 ± 0.05 | 5.30 ± 0.12 | 5.19 ± 0.08 | 5.08 ± 0.12 | 5.35 ± 0.12<br>(5.78 ± 0.11)<sup>g</sup> |
| 13 | *rac-(3R,R<sub>a</sub>)-11f/*<br>*rac-(3R,S<sub>a</sub>)-11f*<sup>d</sup> |  |  | 2 | 6.05 ± 0.09 | 5.33 ± 0.07 | 4.91 ± 0.12 | 4.99 ± 0.04 | 4.90 ± 0.08<br>(5.21 ± 0.08)<sup>g</sup> |
| 14 | *rac-(3R,R<sub>a</sub>)-8g/*<br>*rac-(3R,S<sub>a</sub>)-8g*<sup>d</sup> |  |  | 1 | 4.23 | 4.87 | 81% | 75% | 70% |
| 15 | tiagabine (**3**) |  |  |  | 7.43 ± 0.11<sup>h</sup> | 6.88 ± 0.12<sup>i</sup> | 52%/100 μM<sup>i</sup> | 64%/100 μM<sup>i</sup> | 73%/100 μM<sup>i</sup><br>(78%/250 μM)<sup>i,g</sup> |
| 16 | SK&F-89976-A (**4**) |  |  |  | 6.73 ± 0.02<sup>h</sup> | 6.16 ± 0.05<sup>i</sup> | 3.43 ± 0.07<sup>i</sup> | 3.71 ± 0.04<sup>i</sup> | 3.56 ± 0.06<sup>i</sup><br>(3.00 ± 0.06)<sup>i,g</sup> |
| 17 | (S)-SNAP-5114 (**5**) |  |  |  | 4.56 ± 0.02<sup>j</sup> | 4.07 ± 0.09<sup>k</sup> | 63%/100 μM<sup>k</sup> | 5.29 ± 0.04<sup>k</sup> | 5.65 ± 0.02<sup>k</sup><br>(5.48 ± 0.10)<sup>k,g</sup> |

<sup>a</sup> Compounds were tested as racemates. <sup>b</sup> %ee=77.7. <sup>c</sup> %ee=95.4. <sup>d</sup> Investigated as ~ 1:1 mixture of diastereomeric compounds in racemic form. <sup>e</sup> Data are the mean ± SEM of three or more independent experiments, each performed in triplicate. In case of low inhibitory potencies percentages are given that represent remaining GABA uptake in presence of 100 μM test compound (except for tiagabine (**3**) at hGAT-3, which was applied in a concentration of 250 μM). <sup>f</sup> Results of [<sup>3</sup>H]GABA uptake assays performed with HEK293 cells stably expressing mGAT1-mGAT4 in our laboratory. <sup>g</sup> Results of MS Transport Assays performed with COS-7 cells stably expressing hGAT-3 in our laboratory. <sup>h</sup> Values from reference literature<sup>[42]</sup>. <sup>i</sup> Values from reference literature<sup>33</sup> (mGAT1-mGAT4) and reference literature<sup>[35]</sup> (hGAT-3). <sup>j</sup> Value from reference literature<sup>[43]</sup>. <sup>k</sup> Values from reference literature<sup>[44]</sup>.

As compared to *rac*-nipecotic acid (*rac*-**2**: pIC_50_ = 4.88 ± 0.07) the inhibitory potency of parent compound *rac*-**6** is one log unit lower at mGAT1 (*rac*-**6**: pIC_50_ = 3.89, Table 1, entry 1). Furthermore, compound *rac*-**6** exhibits some but low subtype selectivity for mGAT1. Introducing phenyl groups at the terminal position of the allenic four-carbon atom residue resulted in nipecotic acid derived *rac*-**8a** as a close analogue to the known mGAT1 inhibitor SK&F-89976-A (**4**) from which it differs only by an additional double bond transforming the mono-into a di-unsaturated allenic spacer. Due to this substitution, the inhibitory potency at mGAT1 increased to pIC_50_ = 5.13 ± 0.11 (*rac*-**8a**, Table 1, entry 2), which is now in the range of that of *rac*-nipecotic acid. The mGAT1 inhibitory potency of compound *rac*-**8a**, equipped with an allenic spacer, is, however, about one log unit lower than that of SK&F-89976-A, possessing an alkenyl spacer (**4**: pIC_50_ = 6.16 ± 0.05, Table 1, entry 16), what might be explained by the different flexibility and geometry of both spacer types. Since the nature of the spacer seems to influence the inhibitory potency, in addition to *rac*-**8a** with a four-carbon atom allenic spacer also compound *rac*-**11a** with an extended five-carbon atom allenic spacer has been studied. This spacer extension, however, leads to a decrease in mGAT1 inhibition by 0.52 log units (*rac*-**11a**: pIC_50_ = 4.61 ± 0.00, Table 1, entry 3). The introduction of methyl groups in *para*-position of both phenyl residues in *rac*-**8a** and *rac*-**11a** led to a slight increase of about 0.24 log units for the derivative with a five-carbon atom allenic spacer (*rac*-**11b**: pIC_50_ = 4.85 ± 0.05, Table 1, entry 5), whereas the mGAT1 inhibition potency of the derivative with a four-carbon atom allenic spacer remained unchanged compared to *rac*-**8a** (*rac*-**8b**: pIC_50_ = 5.19 ± 0.02, Table 1, entry 4). An increase in inhibitory potency at mGAT1 was observed when in *rac*-**8b**, with the four-carbon atom allenic spacer, the methyl groups were replaced by fluorine substituents resulting in compound *rac*-**8c** with a pIC_50_ of 5.97 ± 0.10 (Table 1, entry 6). In contrast, the inhibitory potency of related compound *rac*-**11c** with an extended five-carbon atom allenic spacer did not increase with fluorine instead of the methyl substituents, it remained more or less unchanged (*rac*-**8c**: pIC_50_ = 4.89 ± 0.09, Table 1, entry 7). The so far described nipecotic acid derivatives with a terminally double-substituted allenic four- and five-carbon atom spacer *rac*-**8a**-**c** and *rac*-**11a**-**c** displayed a subtype preference for mGAT1. This is least distinct for compound *rac*-**8b** for which the inhibitory potency at mGAT3 and mGAT4 is only slightly lower than at mGAT1 and further slowly declines for mGAT2. A reasonable inhibitory potency was also found for *rac*-**8c** carrying fluorine substituents at the terminal phenyl groups the pIC_50_ at mGAT4 amounting to 4.52. In all other cases pIC_50_ values of the aforementioned compounds at mGAT2 - mGAT4 were around 4 or distinctly below (percentage values of remaining GABA uptake of 50% and above correspond to pIC_50_ values of 4 and below).

An additional gain in mGAT1 inhibition of about 0.5 log units was observed when *p*- chlorine instead of *p*-fluorine is present in the terminal phenyl rings of *rac*-**8c**, resulting in *rac*-**8d** with pIC_50_ = 6.43 ± 0.07 (Table 1, entry 8). Surprisingly, for this compound not only the inhibitory potency at mGAT1 increased but also at mGAT4 which increase was even distinctly higher (∼ 1.5 log units for mGAT4 as compared to 0.5 log units for mGAT1 for *rac*-**8d** as compared to *rac*-**8c**), leading to a notable pIC_50_ value of 6.08 ± 0.05 (Table 1, entry 8). The pIC_50_ values of mGAT4 benchmark inhibitor (*S*)-SNAP-5114 (**5**) displayed in Table 1 are based on repetitive testing over several years of (*S*)-SNAP-5114 as a reference mGAT4 inhibitor in our laboratory, leading to continuously updated mean values (pIC_50_ = 5.65 ± 0.02 at mGAT4, Table 1, entry 17). Accordingly, when *rac*-**8d** was characterized in [^3^H]GABA-uptake-assays on HEK293 cells stably expressing mGAT4 in parallel also (*S*)-SNAP-5114 (**5**) as a reference was studied, what allows a direct comparison of the inhibitory potency of *rac*-**8d** with that of (*S*)-SNAP-5114 (**5**). The obtained pIC_50_ value of (*S*)-SNAP-5114 (**5**) in this experiment was found to be 6.09 ± 0.09, and thus relatively high. Hence, considering the SEM, the pIC_50_ value of *rac*-**8d** at this transporter subtype (*rac*-**8d**: pIC_50_ = 6.08 ± 0.05 at mGAT4, Table 1, entry 8) is identical with that of (*S*)-SNAP-5114 (**5**). However, compound *rac*- **8d** possesses no distinct subtype selectivity for mGAT4, its inhibitory potency at mGAT1 being even higher than that at mGAT4.

For mGAT1 inhibitors delineated from nipecotic acid, it is well known that the biological activity resides mainly in the *R* enantiomer, whereas for mGAT4 inhibitors the *S* enantiomer is more active. Hence, the enantiopure *R* and *S* isomers of *rac*-**8d** had been synthesized and characterized anticipating a further increase of mGAT1 and mGAT4 inhibitory potency and subtype selectivity for the individual enantiomers of *rac*- **8d**. Indeed, as compared to the racemate, the *R* isomer *(R)*-**8d** possesses an increased mGAT1 activity [*(R)*-**8d**: pIC_50_ = 6.81 ± 0.03, Table 1, entry 9], which is in the range of the known inhibitor tiagabine (**3**: pIC_50_ = 6.88 ± 0.12, Table 1, entry 15), whereas mGAT4 inhibition is reduced [*(R)*-**8d**: pIC_50_ = 5.70 ± 0.11, Table 1, entry 9]. The opposite is true for isomer *(S)*-**8d**, for which the inhibitory potency at mGAT1 is lower [*(S)*-**8d**: pIC_50_ = 5.87 ± 0.11, Table 1, entry 10] than that of the racemate *rac*-**8d**. In contrast, the inhibitory potency of *(S)*-**8d** at mGAT4 is distinctly higher than that of the racemic form, the pIC_50_ value reaching 6.59 ± 0.01 (Table 1, entry 10). This is now significantly higher than that of (*S*)-SNAP-5114 (**5**, pIC_50_ = 6.09 ± 0.09) and, to the best of our knowledge, the highest described for mGAT4 inhibitors so far. The lipophilic domain of *(S)*-**8d** consists of two aryl moieties. While such a structural motif is well known as lipophilic domain of mGAT1 inhibitors, it is to our knowledge unprecedented for mGAT4 inhibitors with high potencies. Regarding the subtype selectivity of *(S)*-**8d** for mGAT4 over mGAT1 and mGAT3 the difference of the pIC_50_ values is about 0.6-0.7 log units, whereas for mGAT2 it reaches 1.2 log units. This indicates that in *in vivo* studies a mixed pharmacological profile particularly influenced by the former two GABA transporter subtypes cannot be excluded. The poor subtype selectivity for mGAT4 over mGAT3 is, however, a problem frequently observed for mGAT4 inhibitors.^16, 33, 34, 35^ Isomer *(R)*-**8d** is at least one order of magnitude more potent towards mGAT1 than towards mGAT2, mGAT3 and mGAT4, respectively, indicating a reasonable subtype selectivity over the other subtypes. In order to confirm potencies and to establish pIC_50_ values also at the human equivalent of mGAT4, compounds *rac*-**8d**, *(R)*-**8d**, and *(S)*- **8d** were additionally characterized at hGAT-3 in competitive MS based GABA transport experiments (MS Transport Assays)^35, 36^ utilizing COS cells stably expressing hGAT-3. The results obtained in these experiments at hGAT-3 reassured the high inhibitory potencies of *rac*-**8d** and *(S)*-**8d** initially found at mGAT4 [pIC_50_ values at hGAT-3: *rac*- **8d** 6.35 ± 0.02, Table 1, entry 8; *(S)*-**8d** 6.49 ± 0.10, Table 1, entry 10], whereby *(S)*- **8d** possessed this time only a slightly higher pIC_50_ value than *rac*-**8d**. Accordingly, also the inhibitory potency of *(R)*-**8d** at hGAT-3 [*(R)*-**8d**: pIC_50_ = 5.62 ± 0.06, Table 1, entry 9] is lower than that of the racemic compound *rac*-**8d** and even more than that of its enantiomer *(S)*-**8d**, similar to the results observed for *(R)*-**8d** at mGAT4. In nipecotic acid derivative *rac*-**11d** (Table 1, entry 11) the extension of the allenic spacer to five carbons resulted in a loss of activity at all four transporter subtypes compared to compound *rac*-**8d**, equipped with the four-carbon atom allenic spacer.

To further explore the influence of structural characteristics of these compounds on their potency as GABA uptake inhibitors, two different diaryl residues, a phenyl and a 4’-biphenyl moiety, had been introduced as lipophilic domains at the termini of the allenic spacer of the nipecotic acid derivatives. The respective compounds had been obtained as ∼ 1:1 mixture of racemic diastereomers, i.e. of *rac*-(*3R*,*R_a_*)-**8f**/*rac*-(*3R*,*S_a_*)- **8f** and *rac*-(*3R*,*R_a_*)-**11f**/*rac*-(*3R*,*S_a_*)-**11f** exhibiting a four-carbon and a five carbon atom allenic spacer, respectively, and were tested as these mixtures with regard to their biological activity as GABA uptake inhibitors. At the subtypes mGAT1 - mGAT3 both compound mixtures displayed pIC_50_ values of around 5, though for mGAT4 inhibition, the derivative mixture possessing the shorter allenyl spacer is about 0.45 log units more active than the derivative mixture with extended spacer [*rac*-(*3R*,*R_a_*)-**8f**/*rac*- (*3R*,*S_a_*)-**8f**: pIC_50_ = 5.35 ± 0.12, Table 1, entry 12; *rac*-(*3R*,*R_a_*)-**11f**/*rac*-(*3R*,*S_a_*)-**11f**: pIC_50_ = 4.90 ± 0.08, Table 1, entry 13]. Thereby, the mixture of racemic diastereomers *rac*-(*3R*,*R_a_*)-**8f**/*rac*-(*3R*,*S_a_*)-**8f** with a four-carbon atom allenic spacer has almost equal potencies at all four GABA transporter subtypes, whereas *rac*-(*3R*,*R_a_*)-**11f**/*rac*-(*3R*,*S_a_*)- **11f** with a five-carbon atom allenic spacer is slightly more potent towards mGAT1 than towards mGAT2-mGAT4. The moderate to good potencies at mGAT4 were in good accord with those at hGAT-3, at which even higher pIC_50_ values for both compound mixtures have been obtained [*rac*-(*3R*,*R_a_*)-**8f**/*rac*-(*3R*,*S_a_*)-**8f**: pIC_50_ = 5.78 ± 0.11, Table 1, entry 12; *rac*-(*3R*,*R_a_*)-**11f**/*rac*-(*3R*,*S_a_*)-**11f**: pIC_50_ = 5.21 ± 0.08, Table 1, entry 13].

Finally, to mimic one of the anisole groups present in (*S*)-SNAP-5114 (**5**), the nipecotic acid derived ∼ 1:1 mixture of racemic diastereomers *rac*-(*3R*,*R_a_*)-**8g**/*rac*-(*3R*,*S_a_*)-**8g** with a four-carbon atom allenic spacer of which the terminal allene carbon is part of a methoxy substituted tetrahydronaphthalene ring system has been synthesized. Contrary to the expected results, with pIC_50_ values below 4.87 at all four GABA transporter subtypes (mGAT1-mGAT4), this compound mixture is amongst the least potent inhibitors in this study (Table 1, entry 14).

Overall, the length of the allenic spacer of the studied compounds has a distinct influence on their potencies as GABA uptake inhibitors. The nipecotic acid derivatives with the shorter four-carbon atom allenic spacer give in general rise to the higher inhibitory potencies, with one exception, the terminally unsymmetrically substituted compound mixtures *rac*-(*3R*,*R_a_*)-**8f**/*rac*-(*3R*,*S_a_*)-**8f** and *rac*-(*3R*,*R_a_*)-**11f**/*rac*-(*3R*,*S_a_*)- **11f**. Here, the inhibitory potency at mGAT1-3 is only marginally influenced by the spacer length.

In addition to the inhibitory potencies of the test compounds at mGAT1 (pIC_50_) also their binding affinities (p*K*_i_) for this transporter subtype have been determined. For most of the studied compounds the inhibitory potencies at mGAT1 were about a half to one log unit lower than the corresponding p*K*_i_ values. Such a difference between pIC_50_ and p*K*_i_ values is a common phenomenon constantly observed for mGAT1 inhibitors when characterized in this test system.^37, 38, 39, 40, 41^

Hence, the conclusions drawn for the above discussed pIC_50_ values at mGAT1 are equally well supported by the observed p*K*_i_ values.

#### *In vivo* part

Considering the results obtained in the *in vitro* part of this present research, *(S)*-**8d** also termed DDPM-3960 was further selected for *in vivo* studies which aimed to assess its impact on the central nervous system function in mice. For this purpose, intraperitoneally administered DDPM-3960 at doses 1.25-60 mg/kg was first tested in mouse models of seizures that were induced electrically (maximal electroshock test, 6-Hz test and electroconvulsive threshold test) and chemically (pentylenetetrazole (PTZ) test, pilocarpine test). We also assessed its effect on motor functions (rotarod test, grip strength test, locomotor activity test). Since the anticonvulsant active hGAT- 1 inhibitor, tiagabine shows anxiolytic properties both in humans and experimental animals and it has a potential antidepressant-like activity in mice,^45^ anxiolytic-like and antidepressant-like activities of DDPM-3960 were also assessed in the four-plate, elevated plus maze tests, and in the forced swim test. The effect of DDPM-3960 on pain threshold was additionally assessed in the acute, thermally-induced (hot plate test) or acute, chemically-induced (capsaicin test), and tonic, chemically-induced (formalin test) pain models as antiepileptic drugs are widely used as analgesics in some pain types.^46, 47^

##### Anticonvulsant activity

###### PTZ seizure test

In the PTZ test an overall effect of treatment on latency to seizure onset was observed (F[4,38]=24.02, p < 0.0001). DDPM-3960 at doses 10, 30 and 60 mg/kg significantly (p < 0.0001 vs. control group) delayed the onset of PTZ seizures. The dose 5 mg/kg was not effective (Fig. 2A).

**Fig. 2.**
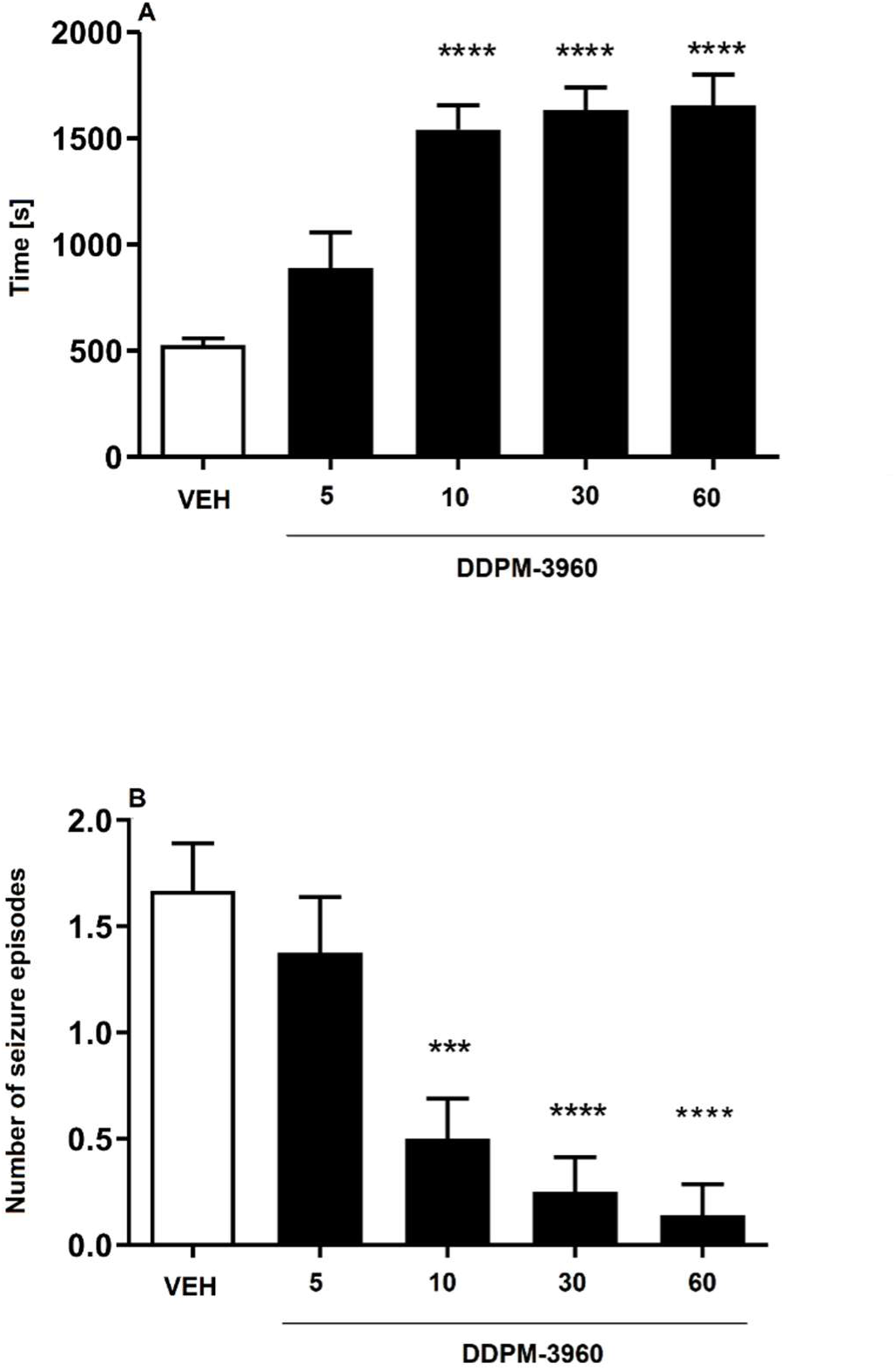
Influence of DDPM-3960 used at doses 5, 10, 30 and 60 mg/kg on latency to first clonus (A) and the number of seizure episodes (B) induced by PTZ. Results are shown for n = 7-12. Statistical analysis: one-way ANOVA followed by Dunnett’s *post hoc* test. Significance vs. vehicle-treated mice: *** p < 0.001, **** p < 0.0001.

DDPM-3960 also influenced the number of PTZ-induced seizure episodes (F[4,38]=11.17, p < 0.0001). *Post hoc* analysis revealed that DDPM-3960 (doses: 10, 30 and 60 mg/kg) significantly (p < 0.001) and dose-dependently reduced the number of seizure episodes caused by PTZ (Fig. 2B).

Mortality rate in PTZ-treated mice was also assessed. It has been revealed that 16% of control mice died after seizure episodes caused by PTZ administration. In DDPM- 3960 (5 mg/kg)-treated group 13% of mice died after seizure episode caused by PTZ administration, while in all other groups treated with DDPM-3960 no mortality was noted.

##### Maximal electroshock seizure test

In MES test the compound DDPM-3960 (10, 30 and 60 mg/kg) dose-dependently reduced the number of mice with tonic hind limb extension. The dose 60 mg/kg prevented seizures in all mice tested (Tab. 2).

**Table 2.** Anticonvulsant activity of compound DDPM-3960 in MES test.

| Compound | Dose [mg/kg] | X/Ya (% of mice protected) |
| --- | --- | --- |
| Vehicle | - | 0/6 (0) |
| **(S)-8d** | 10 | 1/6 (17) |
|  | 30 | 4/6 (67) |
|  | 60 | 6/6 (100) |

<sup>a</sup> Results are shown as the number of mice protected (X) per the number of mice tested (Y).

In this test, the ED_50_ defined as the dose that protected 50% of mice from seizures was established (18.7 mg/kg).

##### Electroconvulsive threshold test

The effect of DDPM-3960 on the electroconvulsive threshold was assessed for 2 doses of the test compound, i.e. 30 and 60 mg/kg. The CS_50_ (median current strength, i.e. current intensity that induced tonic hind limb extension in 50% of the mice challenged) for the vehicle-treated group was 8.6 mA. The CS_50_ value for DDPM-3960 used at 30 mg/kg was 27.1 mA. For the dose 60 mg/kg of DDPM-3960, CS_50_ could not be established as this dose protected all animals tested from seizures in the range of current intensities tested (max. current intensity offered by the rodent shocker was 34 mA).

##### 6-Hz test

In this mouse model of psychomotor seizures the compound DDPM-3960 was less effective than in the PTZ or MES tests. The dose 60 mg/kg reduced seizures in 50% of animals tested (Table 3).

**Table 3.** Anticonvulsant activity of DDPM-3960 measured in 6-Hz test.

| Compound | Dose (mg/kg) | X/Ya(% of mice protected) |
| --- | --- | --- |
| Vehicle | - | 0/6 (0) |
| **(S)-32d** | 30 | 2/6 (33) |
|  | 60 | 3/6 (50) |

<sup>a</sup> Results are shown as the number of mice protected (X) per the number of mice tested (Y).

##### Pilocarpine-induced seizures

The pilocarpine seizure test is used to model human temporal lobe epilepsy.^48^ This assay is also regarded as a rodent model of *status epilepticus*.^48^ In this test an overall effect of treatment on latency to *status epilepticus* was observed (F[2,20]=8.759, p < 0.01). *Post hoc* analysis revealed that both doses of DDPM-3960 (30 and 60 mg/kg) significantly (p < 0.01) prolonged latency to pilocarpine-induced *status epilepticus* but the dose 30 mg/kg was slightly more effective (Fig. 3A).

**Fig. 3.**
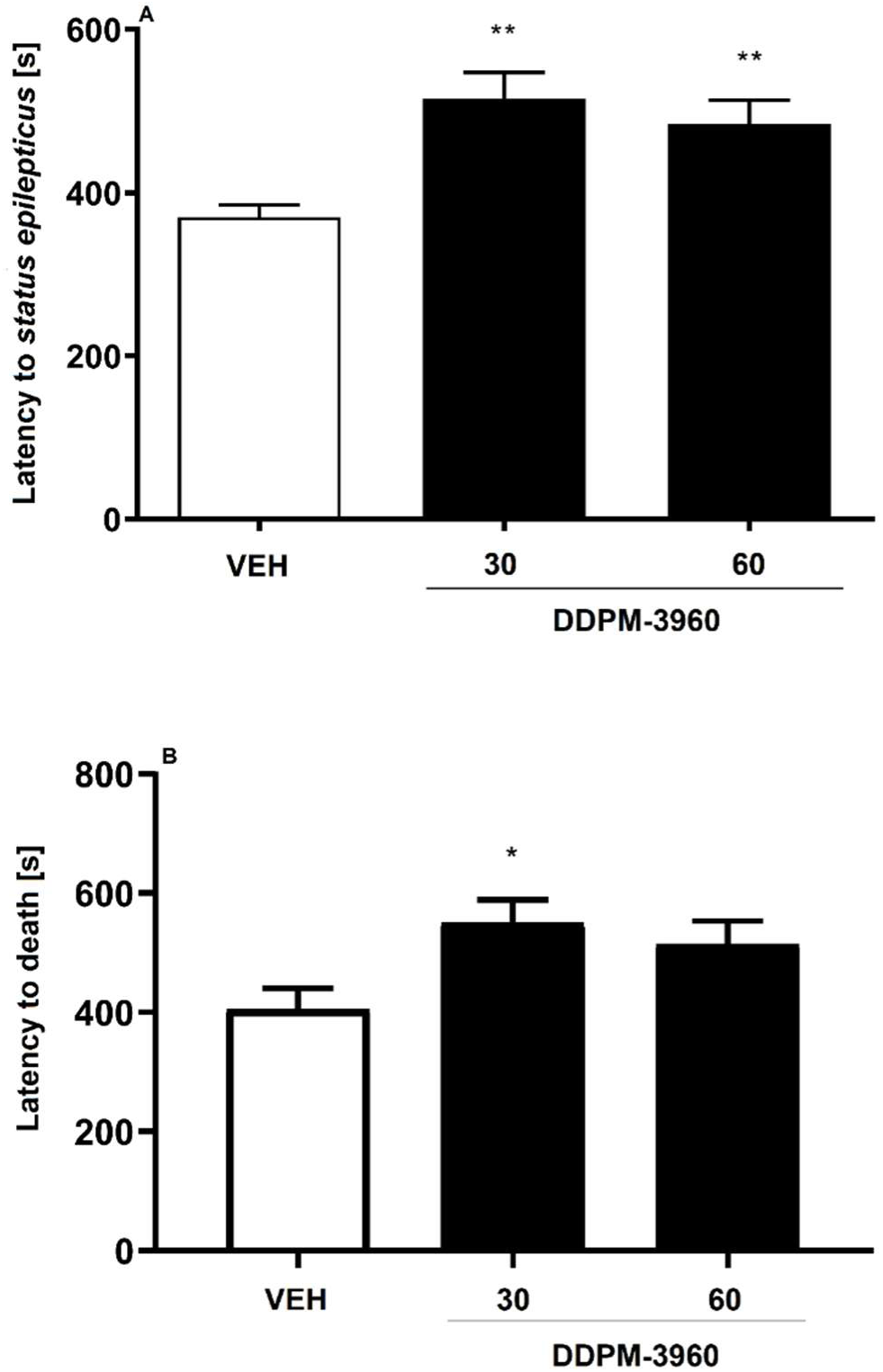
Influence of the compound DDPM-3960 used at the doses of 30 and 60 mg/kg on latency to pilocarpine-induced *status epilepticus* (A), and latency to death (B). Results are shown as mean latency (± SEM) for n = 7-8. Statistical analysis: one-way ANOVA followed by Dunnett’s *post hoc* comparison. Significance vs. vehicle-treated mice: * p < 0.05, ** p < 0.01.

One-way ANOVA also revealed a significant effect of treatment on the latency to death (F[2,20]=4.077, p < 0.05). Compared to the vehicle-treated group, only the dose 30 mg/kg of DDPM-3960 significantly (p < 0.05) prolonged latency to death in experimental animals (Fig. 3B).

In the *in vivo* research five distinct mouse models of seizures were implemented. Two of them, i.e. MES test and PTZ test are regarded as simple screening seizure models that allow to test large numbers of compounds for their potential anticonvulsant activity in relatively short time.^49^ These ‘gold standard’ screening assays are not only used to identify novel antiepileptic drugs (AEDs), but they also enable to predict the efficacy of AEDs against different types of seizures in humans.**^Fehler! Textmarke nicht definiert.^** The PTZ model of clonic seizures generally refers to nonconvulsive (absence or myoclonic) seizures in humans, whereas the MES test is predictive for generalized tonic-clonic seizures.**^Fehler! Textmarke nicht definiert.^** The efficacy of the compound DDPM-3960 in the PTZ model and its anticonvulsant activity demonstrated in the MES test suggest that this compound might be potentially effective in absence seizures and in grand mal epilepsy in humans.

Since in 6-Hz test (a model of psychomotor seizures) DDPM-3960 was less active, it seems that in human psychomotor seizures this compound would have limited protective efficacy.

In contrast to 6-Hz test, both doses of DDPM-3960 had a strong influence on the electroconvulsive threshold assessed in the electroconvulsive threshold test. This test is very sensitive to GAT1 inhibitors^45, 50^ and it indicates that GAT inhibition due to DDPM-3960 administration might underlie its *in vivo* effect.

The observed efficacy of the compound DDPM-3960 in PTZ test and MES test might also suggest a potential mechanisms of its action. Drugs acting as antagonists of voltage-gated sodium and to a lesser degree calcium channels (with the exception of ethosuximide) are effective in MES test, and they show no anticonvulsant protection in PTZ model, whereas numerous GABAergic AEDs, including GAT inhibitors^45, 51^ and the T-type calcium channel blocker ethosuximide are effective in PTZ model. Mixed AEDs are active in both tests and this effect is thought to be due to their multi-target profile of the anticonvulsant activity.**^Fehler! Textmarke nicht definiert.^**^, 52^ Considering this, such a mixed profile of DDPM-3960 should be potentially taken into consideration and the mechanism of DDPM-3960 action seems to be broader (hGAT-1, hGAT-3 and possibly other, non-GAT targets) than that of tiagabine as in the previous studies tiagabine, a selective hGAT-1 inhibitor, was not effective in the MES test**^Fehler! Textmarke nicht definiert.^**^, 45^ but it was effective in PTZ test.^45, 53^

Similarly to the PTZ model, pilocarpine-induced seizure test was the second animal model that used a chemoconvulsant drug to induce seizures. In contrast to PTZ model, pilocarpine evokes M1 muscarinic receptor-dependent spontaneous recurrent seizures,^48^ that mimic *status epilepticus* in humans and pilocarpine is regarded as a very useful tool to study drug-resistant epilepsy.^54, 48^ In this model of human temporal lobe epilepsy^48^ the compound DDPM-3960 was also effective. Based on the available evidence from different laboratories, it appears that the pilocarpine model may represent a suitable tool for investigating the mechanisms of temporal lobe epilepsy.^48^ Experiments in cultured hippocampal neurons have demonstrated that pilocarpine causes an imbalance between excitatory and inhibitory transmission resulting in the generation of status epilepticus and *in vivo* microdialysis studies have revealed that pilocarpine induces elevation in glutamate levels in the hippocampus resulting in the appearance of seizures which are maintained by NMDA receptor activation.^48^ Considering this, it seems plausible that hGAT-1, hGAT-3 inhibition due to DDPM-3960 administration might attenuate seizures caused by the hyperactivation of the excitatory amino acid system.

##### Motor coordination

###### Rotarod test

In the rotarod test no motor deficit were observed in DDPM-3960-treated mice, either for 10 or 30 mg/kg. Motor impairments, defined as inability to remain on the rotating rod for 1 min, were noted in mice that received DDPM-3960 at the dose of 60 mg/kg (p < 0.01 vs. vehicle-treated mice).

###### Grip strength test

The test compound DDPM-3960 used at doses 10, 30, 60 mg/kg did not affect animals’ muscular strength measured in the grip strength test (F[3,20]=1.522, p > 0.05).

To sum up, motor coordination deficits are one of the most frequent adverse effects of AEDs and this effect might be a serious limitation of antiepileptic pharmacotherapy. We demonstrated that the test compound DDPM-3960 only at the highest dose tested induced some motor deficits in the rotarod test, but it did not influence negatively muscular strength of experimental animals.

##### Effect on locomotor activity

The locomotor activity test was used to assess the effect of DDPM-3960 on animals’ general activity. Drug-induced decrease of locomotor activity (i.e. sedation) or hyperlocomotion might give false positive results in some *in vivo* tests. To exclude this obstacle, we measured locomotor activity of mice at selected time points at which further assays were conducted: 1 min (important for the assessment of the results obtained in the four-plate and hot plate tests), 5 min (important for the assessment of the results obtained in the elevated plus maze test, capsaicin test and the first phase of the formalin test), 6 min (important for the assessment of the results obtained in the forced swim test) and 30 min (important for the assessment of the results obtained in the second phase of the formalin test).

Locomotor activity was significantly affected by treatment (F[3,112]=128.1, p < 0.0001). Time also affected the results significantly (F[3,112]=159.5, p < 0.0001) and drug x time interaction was also significant (F[9,112]=49.67, p < 0001). *Post hoc* analysis revealed that in the 1^st^ min of the test DDPM-3960 did not affect animals’ locomotor activity. In the 5^th^ and in the 6^th^ mins of the locomotor activity test the doses 30 and 60 mg/kg significantly (p < 0.05 and p < 0.01 vs. control mice, for both time points, respectively) reduced locomotor activity of mice. In the 30^th^ min of the test, all doses of DDPM-3960 significantly (p < 0.0001) affected locomotor activity of mice. At this time point the dose 2.5 mg/kg increased, while doses 30 and 60 mg/kg decreased the number of light-beam crossings (Fig. 4).

**Fig. 4.**
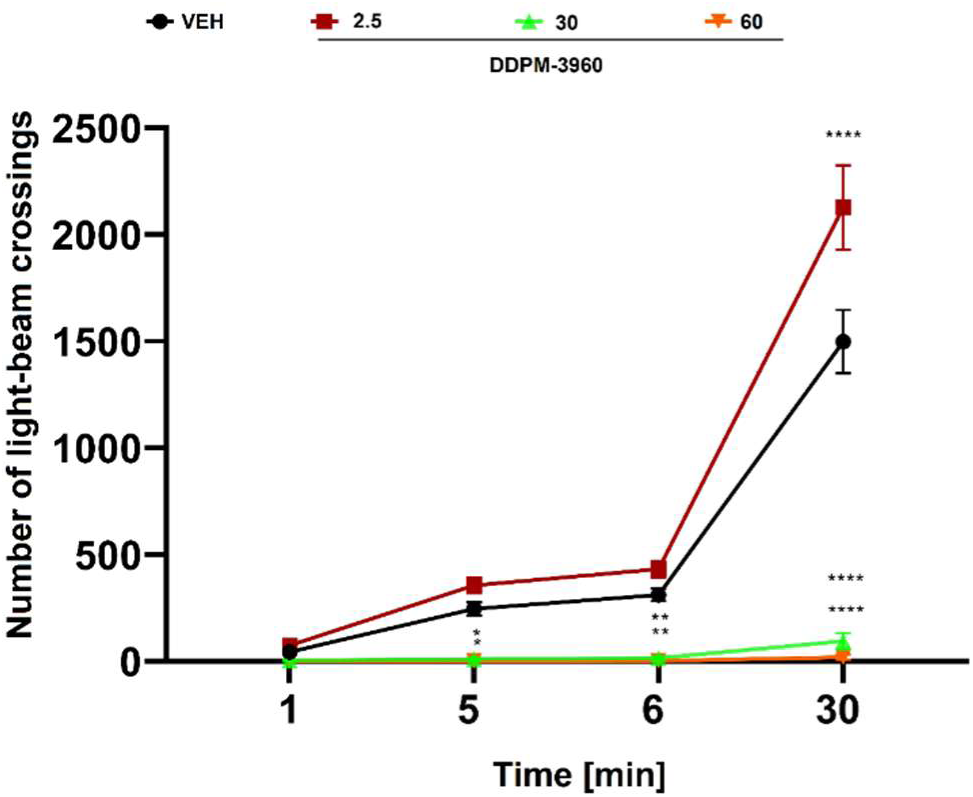
Influence of the test compound DDPM-3960 on the locomotor activity of mice measured during the 30-min observation period. Results are shown as mean number of light-beam crossings (±SEM) measured at selected time points: 1 min, 5 min, 6 min and 30 min for n = 8. Statistical analysis of the results was conducted using repeated-measures ANOVA followed by Dunnett’s *post hoc* comparison. Significance vs. vehicle-treated mice at the respective time point: * p < 0.05, ** p < 0.01, **** p < 0.0001.

##### Anxiolytic-like activity

###### Four-plate test

In the four-plate test an overall effect of treatment with DDPM-3960 was observed (F[5,50]=6.743, p < 0.0001). *Post hoc* analysis revealed that the dose of 2.5 mg/kg significantly (p < 0.0001) increased the number of punished crossings. This suggests that DDPM-3960 at this dose displayed anxiolytic-like properties (Fig. 5).

**Fig. 5.**
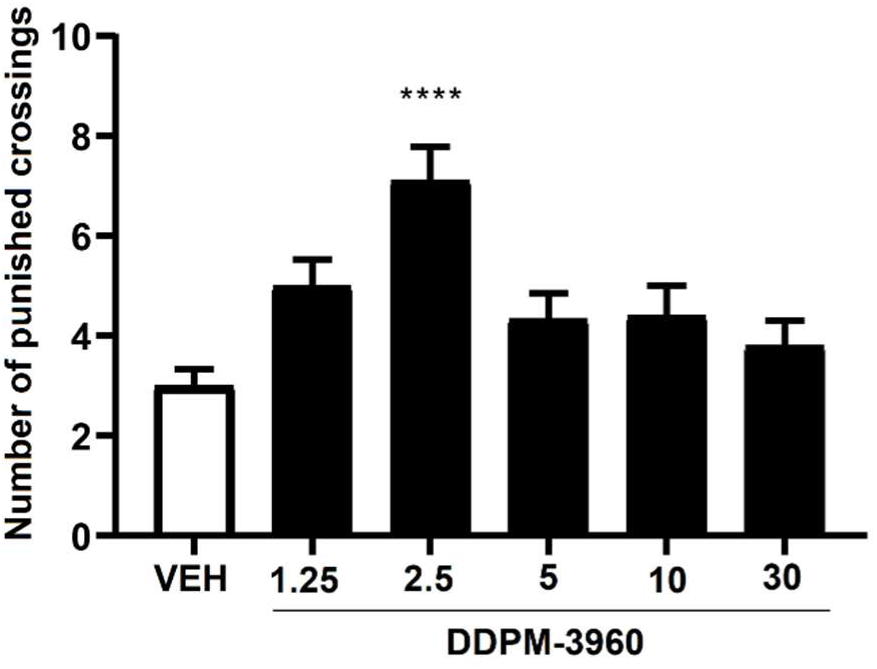
Anxiolytic-like activity of the compound DDPM-3960 in the mouse four-plate test. Results are shown as the mean number of punished crossings (±SEM) for n = 8-10. Statistical analysis: one-way ANOVA followed by Dunnett’s *post hoc* comparison. Significance vs. vehicle-treated group: **** p < 0.0001.

###### Elevated plus maze test

In the elevated plus maze test the dose 2.5 mg/kg of DDPM-3960 was assessed to further confirm whether this dose displayed anxiolytic-like properties. Compared to the vehicle-treated group, DDPM-3960 increased both time spent in open arms (significant at p < 0.0001, Fig. 6A) and % time spent in open arms (significant at p < 0.0001, Fig. 6B). Compared to control, it also significantly (p < 0.001) increased the number of open arm entries (Fig. 6C). The % of open arm entries was however similar in both control and DDPM-3960 groups (Fig. 6D).

**Fig. 6.**
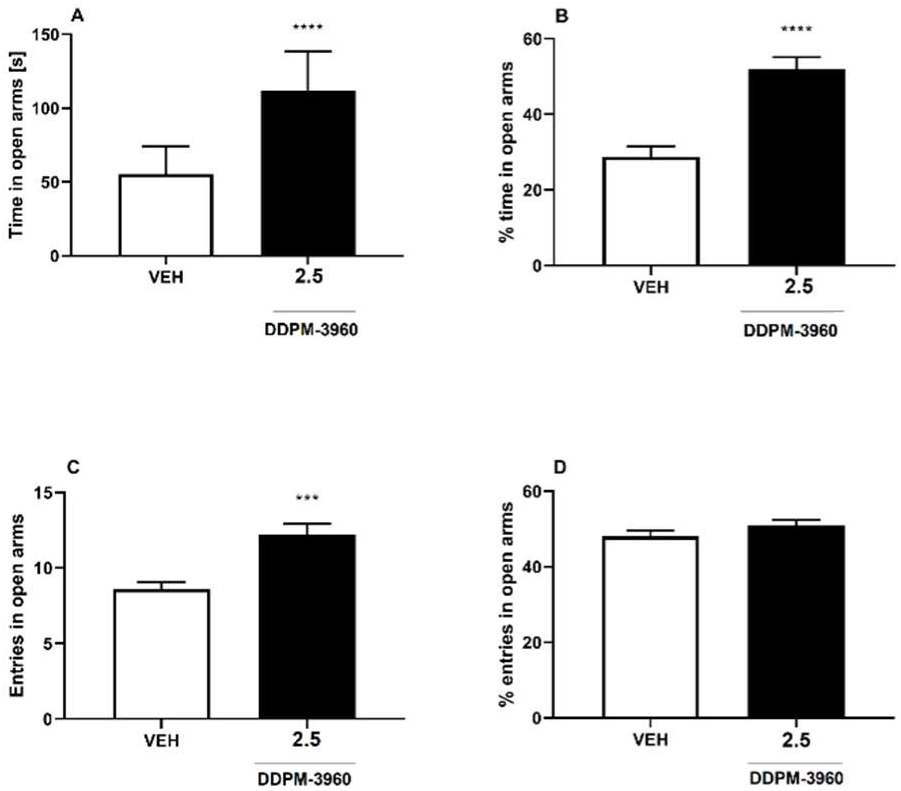
Anxiolytic-like activity of DDPM-3960 (2.5 mg/kg) in the elevated plus maze test. Results are shown as mean time spent in open arms (A), % time spent in open arms (B), entries in open arms (C) and % entries in open arms (D) for n = 10. Statistical analysis: Student’s t-test. Significance vs. vehicle-treated group: *** p < 0.001, **** p < 0.0001.

Considering the results from the locomotor activity test (no sedation or hyperlocomotion 1 min after the locomotor activity test started; see Fig. 4) and those obtained for DDPM-3960 used at the dose of 2.5 mg/kg in the four-plate test, it seems that DDPM-3960 at the dose of 2.5 mg/kg has anxiolytic-like properties in mice. This effect might be due to its effect on GAT, in particular mGAT4.^18^

This anxiolytic-like effect was also confirmed in the elevated plus maze test which is a complementary assay for the four-plate test. The elevated plus maze test lasts for 5 min and it should be noted here that for the dose of 2.5 mg/kg no unwanted effects of DDPM-3960 on animals’ locomotor activity were observed at this time point. Hence, potential false positive results of the elevated plus maze test should be excluded. Taken together, both the four-plate test and the elevated plus maze test demonstrated statistically significant anxiolytic-like properties of DDPM-3960 used at the dose of 2.5 mg/kg in mice.

##### Antidepressant-like activity

###### Forced swim test

In the forced swim test an overall effect of treatment with DDPM-3960 was not observed (F[3,28]=0.9800, p > 0.05). The test compound did not show antidepressant-like properties at any dose tested (Fig. 7).

**Fig. 7.**
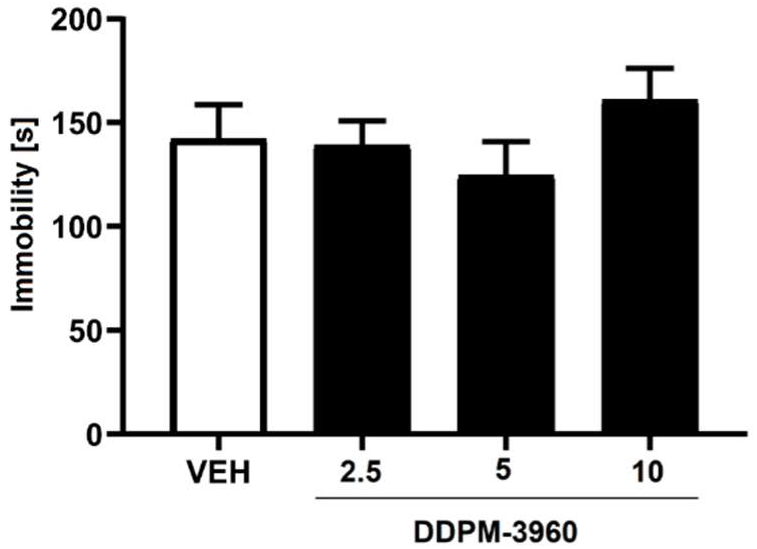
Antidepressant-like activity of the compound DDPM-3960 used at doses 2.5, 5 and 10 mg/kg in the mouse forced swim test. Results are shown as the mean duration of immobility (±SEM) for n = 8. Statistical analysis: one-way ANOVA followed by Dunnett’s *post hoc* comparison. Significance vs. vehicle-treated group: p > 0.05.

Although previous studies reported the involvement of mGAT4^18^ and mGAT1^55, 56^ in depression-like behavior in mice, the forced swim test did not reveal antidepressant-like properties of DDPM-3960 used at doses 2.5, 5 and 10 mg/kg.

##### Analgesic activity

###### Acute, thermally-induced pain (hot plate test)

In the hot plate test repeated-measures ANOVA did not reveal a statistically significant effect of treatment on the latency to pain reaction (F[2,50]=1.629, p > 0.05). Time effect and drug x time interaction were not significant, either (F[1,50]=0.4443, p > 0.05 and F[2,50]=0.01686, p > 0.05, respectively). *Post hoc* analysis showed that DDPM-3960 (10, 30 mg/kg) was not effective in this assay (Fig. 8).

**Fig. 8.**
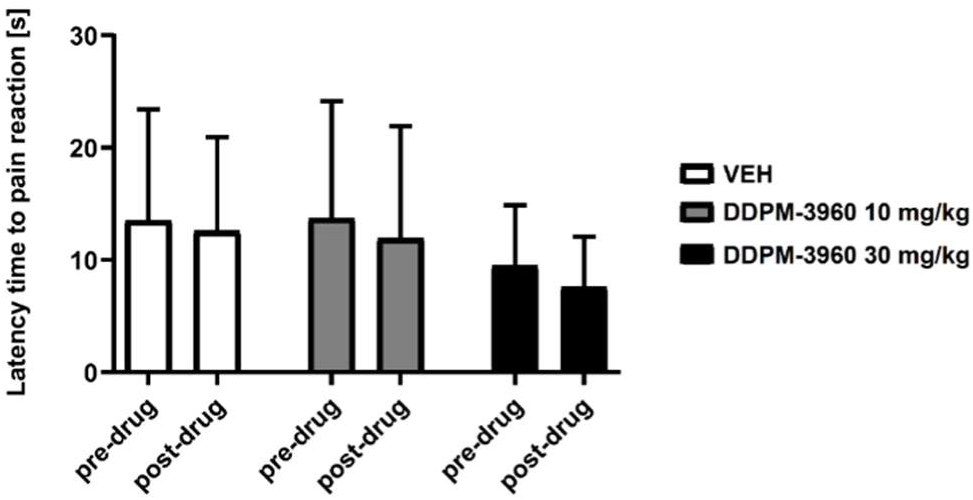
Antinociceptive activity of the compound DDPM-3960 at doses 10 and 30 mg/kg in the hot plate test. Results are shown as mean latency to pain reaction (±SEM) in response to thermal stimulus (55 °C) for n = 9-10. Statistical analysis: repeated-measures ANOVA followed by Dunnett’s *post hoc* comparison. Significance vs. vehicle-treated group at the respective time point: p > 0.05.

###### Neurogenic pain model (capsaicin test)

In the capsaicin test one-way ANOVA revealed an overall effect of treatment on the duration of pain reaction (F[2,19]=15.40, p < 0.0001). *Post hoc* analysis showed that DDPM-3960 at the dose of 30 mg/kg significantly (p < 0.0001) reduced the duration of the licking response in capsaicin-treated mice (Fig. 9).

**Fig. 9.**
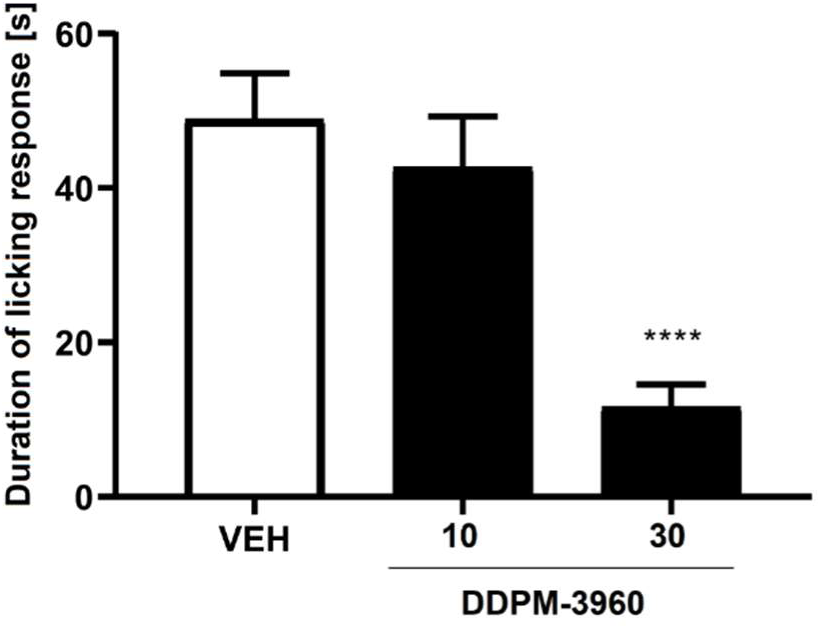
Antinociceptive activity of the compound DDPM-3960 used at doses 10 and 30 mg/kg measured in the mouse capsaicin test. Results are shown as the mean duration of the licking/biting response (±SEM) for n = 6-8. Statistical analysis: one-way ANOVA followed by Dunnett’s *post hoc* comparison. Significance vs. vehicle-treated group: **** p < 0.0001.

###### Tonic pain model (formalin test)

In the formalin test antinociceptive properties of DDPM-3960 at doses 10 and 30 mg/kg were tested. Its influence on neurogenic and inflammatory pain were assessed in the early phase (phase I) and in the late phase (phase II) of this test, respectively.

In the neurogenic phase, an overall effect of treatment on the duration of pain reaction was observed (F[2,21]=12.35, p < 0.001). In this phase the dose 30 mg/kg of DDPM- 3960 significantly (p < 0.01) reduced the nocifensive response of formalin-treated mice (Fig. 10A). One-way ANOVA also revealed a significant effect of treatment in the second phase of the formalin test (F[2,16]=48.63, p < 0.0001). *Post hoc* analysis showed that in this phase DDPM-3960 at the dose of 30 mg/kg reduced the duration of licking/biting response (significant at p < 0.0001, Fig. 10B).

**Fig. 10.**
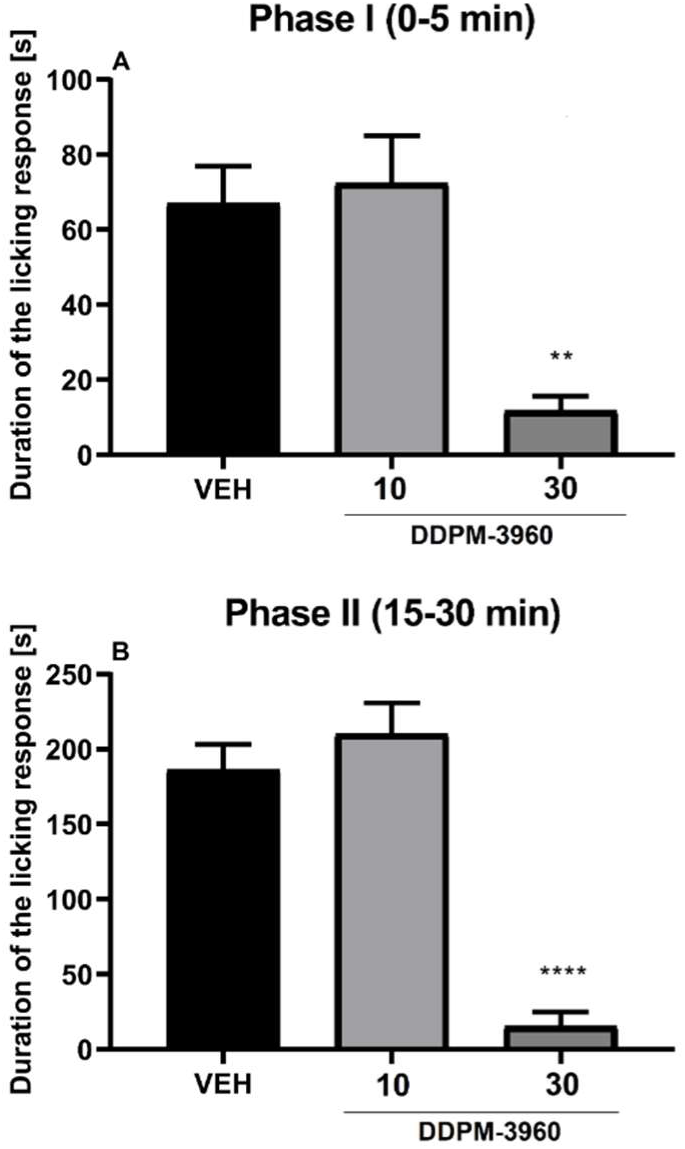
Antinociceptive activity of the compound DDPM-3960 in the mouse formalin test. Results are shown as the mean duration of the licking/biting response (±SEM) in the formalin-injected paw during the early phase (0-5 min) and the late phase (15-30 min) of the test for n = 6-8. Statistical analysis: one-way ANOVA followed by Dunnett’s *post hoc* comparison. Significance vs. vehicle-treated group in the respective phase: ** p < 0.01, **** p < 0.0001.

In order to assess the effect of DDPM-3960 on pain threshold in mice we carried out three behavioral tests in which pain was induced by thermal (hot plate test) or chemical (capsaicin test, formalin test) stimuli. In the hot plate test which is a model of acute pain analgesic effects of DDPM-3960 were not shown. In the hot plate test centrally-acting analgesics can be detected. In contrast to this, DDPM-3960 showed analgesic properties in neurogenic pain (capsaicin test, the first phase of the formalin test). It should be however noted that: (1) this effect was shown only for the dose 30 mg/kg (and not for 10 mg/kg) and (2) this effect of DDPM-3960 in pain tests was assessed 5 min after the test started. Comparing the results obtained in pain tests for the dose 30 mg/kg of DDPM-3960 with those from the locomotor activity test (time point: 5 min), it can be concluded that results obtained in both pain tests (capsaicin test, early phase of the formalin test) are rather false positive ones (in the locomotor activity test a significant decrease of locomotor activity of mice was noted for the dose 30 mg/kg 5 min after the test started). Similar conclusions can be drawn for DDPM-3960 at the dose of 30 mg/kg referring to the locomotor activity results obtained for the time point 30 min and results obtained in the second phase of the formalin test. Although in the late phase of the formalin test the dose 30 mg/kg significantly reduced pain behavior in mice, these results seem to be false positive as the observed decrease in licking/biting behavior should rather be attributed to strong sedative properties of DDPM-3960 30 mg/kg.

## Conclusion

In summary, a new series of nipecotic acid derived GABA uptake inhibitors, characterized by an allenic four- and five-carbon atom spacer, bearing lipophilic aryl residues at the spacer terminus, has been synthesized and biologically characterized. Compounds equipped with the shorter four-carbon atom allenic spacer have been identified to possess higher biological activity than those with an allenic five-carbon atom spacer. The synthesis of nipecotic acid derivatives with terminally double-substituted allenic spacers was accomplished by two efficient cross-coupling reactions, whereby diaryl ketones as starting material have been implemented in Cu^I^-catalyzed cross-couplings of related ketone derived diaryldiazomethanes with nipecotic acid derived terminal alkynes, and substrate 6-methoxy-1-tetralone has been transformed in an analogue cross-coupling reaction via the corresponding *N*-tosylhydrazone. Among the biologically investigated compounds derivative DDPM-3960 [(*S*)-**8d**], exhibiting two 4-chlorophenyl residues connected *via* the four-carbon atom allenic spacer with the amino nitrogen of the polar nipecotic acid head, was identified as highly potent mGAT1 and mGAT4 inhibitor. The *R* enantiomer (*R*)-**8d** displays an inhibitory potency at mGAT1 that is in the range of the known inhibitor tiagabine (**3**). For the enantiopure *S* isomer DDPM-3960 [(*S*)-**8d**] pIC_50_ values of 6.59 ± 0.01 at mGAT4 and 6.49 ± 0.10 for the human equivalent hGAT-3 have been determined, which are significantly higher than that of the benchmark mGAT4 inhibitor (*S*)-SNAP-5114 (**5**). The combination of a rigid allenyl spacer with a diaryl residue as lipophilic domain in compound DDPM-3960 possesses a novel structural scaffold for this kind of bioactive compounds. Unfortunately, the subtype selectivity of DDPM-3960 for mGAT4 over the other three subtypes is not very pronounced.

*In vivo,* DDPM-3960 was more effective as an anticonvulsant compound as compared to previously tested tiagabine. In our earlier study^45^ tiagabine demonstrated anticonvulsant properties in chemically induced seizures (PTZ and pilocarpine seizures). At the dose of 100 mg/kg tiagabine also elevated the seizure threshold for electrically induced seizures by 31.6%, but it had no activity in the MES test. Similarly to tiagabine, DDPM-3960 showed anxiolytic-like properties in mice but in contrast to tiagabine it did not display antidepressant-like effects. Although it apparently reduced animals’ nociceptive responses in some pain tests, these activities should rather be regarded as false positive outcomes resulting from its sedative properties shown in the locomotor activity test.

The observed *in vivo* differences between DDPM-3960 and tiagabine are likely to be due to their distinct profile of GAT affinity. A broader (compared to tiagabine) anticonvulsant profile in mouse tests can be attributed to DDPM-3960’s synergistic effect on mGAT4 and mGAT1. Other (GAT-independent) mechanisms should also be considered.

Overall, the herein described compounds represent a new structural motif for mGAT4 inhibitors, which could serve as useful starting points for the further expansion of structure-activity relationships of mGAT4 inhibitors, facilitate the design of new ligands with higher potencies and better selectivities at this molecular target and even higher efficacy in animal models.

## Experimental Section

All commercially available reagents were used without further purification. Unless otherwise noted, all reactions were performed in oven dried glassware under moisture-free conditions and argon- or nitrogen atmosphere. Toluene and 1,4-dioxane were dried over sodium and distilled under nitrogen. Chlorobenzene was dried over CaCl_2_, distilled under nitrogen atmosphere and stored over molecular sieves (4 Å) under nitrogen atmosphere prior to use. For chromatographic purposes only distilled solvents were used (EtOAc, *iso*-hexane/PE 42-62 °C, dichloromethane/DCM, MeOH, *n*- pentane, Et_2_O, H_2_O). Flash column chromatography was performed using silica gel (grading 0.035-0.070 mm). Thin layer chromatography (TLC) was carried out on precoated silica gel F_254_ glass plates. Preparative MPLC was performed using a Büchi instrument (C-605 binary pump system, C-630 UV detector at 254 nm and C-660 fraction collector) and a Sepacore glass column B-685 (26×230 mm) equipped with YMC Gel SIL-HG (12 nm, 5-20 µm) for straight phase and YMC Gel Triart Prep C18- S (12 nm, S-20 µm) for reverse phase. NMR spectra were recorded on a JNMR-GX (JEOL) at room temperature (unless otherwise noted) on 400 MHz (^1^H NMR: 400 MHz, ^13^C NMR: 101 MHz) and 500 MHz (^1^H NMR: 500 MHz, ^13^C NMR: 126 MHz) spectrometers. These NMR spectrometers were also used for DEPT, HMQC, HMBC and COSY experiments. ^1^H and ^13^C NMR chemical shifts were referenced to the deuterated solvent signals and the coupling constants were stated with an accuracy of 0.5 Hz. MestreNova software was used for further analysis of the NMR data. Broadened signals in ^1^H NMR spectra were supplemented by the index “br” (s_br_, d_br_, t_br_) and signals from mixtures of diastereomeric compounds in racemic form were labeled as “dia1” or “dia2”. IR spectra were recorded with a FT-IR spectrometer and Spectrum v2.00 software was used for analysis. Samples were measured either as KBr pellets or as films on NaCl plates. High resolution mass spectrometry was carried out with a LTQ FT Ultra mass spectrometer (ThermoFinnigan). Optical rotations were determined by a 241 MC Polarimeter ADP440+ at λ=589 cm^-1^. Purity testing of biologically tested compounds *rac*-**11a** and *rac*-**11d** was done by Quantitative ^1^H NMR (qHNMR)^57, 58^ and purity was determined as ≥95%. QHNMR data based on peak area ratios are determined under conditions that assure complete relaxation. For qHNMR the internal standard maleic acid (TraceCERT^®^, Sigma Aldrich, Lot#BCBM8127V: purity 99.94%) was dissolved in MeOD. The purity was calculated using the purity calculator of MestreNova NMR software (MestreLab Research S.L.). For compounds *rac-***8a-d**, (*R*)-**8d**, (*S*)-**8d**, *rac*-(*3R*,*R_a_*)-**8f**/*rac*-(*3R*,*S_a_*)-**8f**, *rac*-**11b**-**c**, *rac*-(*3R*,*R_a_*)- **11f**/*rac*-(*3R*,*S_a_*)-**11f** and *rac*-(*3R*,*R_a_*)-**8g**/*rac*-(*3R*,*S_a_*)-**8g** purity testing was done by means of analytical HPLC on an Agilent 1100 instrument (G1329A ALS autosampler, G1316A column compartment, G1314A VWD detector, G1312A binary pump, G1379A degasser), equipped with a Lichrospher 100 RP-18 (5 μm) in a LiChroCART 250-4 column, with elution at 0.5 mL/min with ammonium formate buffer (10 mM, pH 6.8) to MeOH 20:80 and purity was determined as ≥95%.

### General Procedure for the N-Alkylation (GP1)

N-alkylation was performed employing a synthesis route similar to a procedure described in literature.^59^ Alkyne (1.00 equiv) and amine (1.20 equiv) were dissolved in acetone (2 mL/mmol) and Na_2_CO_3_ (2.50 equiv) and NaI (0.50 equiv) were added. The reaction mixture was refluxed for 72 h and the reaction was monitored by TLC. For quenching, DCM (5 mL/mmol) and water (5 mL/mmol) were added and the product was extracted with DCM. The combined organic phases were then dried over Na_2_SO_4_ and concentrated under vacuum. The crude product was purified by flash column chromatography to afford the desired alkyne.

### General Procedure for the Synthesis of Tosylhydrazones (18) by Literature Procedure^60^ (GP2)

A solution of pure TsNHNH_2_ (1 equiv) in methanol (1 mL/mmol) was stirred and heated to 60 °C until the TsNHNH_2_ dissolved. The mixture was cooled to rt. Then the corresponding aldehyde (1 equiv) was dropped slowly to the mixture. The crude product could be obtained as needle shaped crystals. The colorless crystals were collected by filtration and washed with PE and were then dried under vacuum to afford the desired product.

### General Procedure for the Preparation of Diaryldiazomethanes 13a-f by Literature Procedure^30^ (GP3)

Hydrazine monohydrate (80% purity) was added to a solution of diarylmethanone in ethanol. Then, aqueous HCl (37%, 0.50 mL/20 mmol ketone) was added and the mixture was heated to reflux overnight. After cooling to room temperature, the diarylmethanone hydrazone was precipitated as needle shaped crystals. Filtration of the crude mixture gave pure diarylmethanone hydrazones. A mixture of diarylmethanone hydrazone (10.0 mmol, 1.0 equiv), anhydrous MgSO_4_ (1.00 g) and 30.0 mL DCM was cooled to 0 °C. To this rapidly stirred mixture activated MnO_2_^[61]^ (3.04 g, 35.0 mmol, 3.5 equiv) was added in one portion. The reaction mixture was warmed to room temperature and kept stirring for 8 h and then the solid was filtered off with a celite pad and washed with DCM. After removal of the solvent under reduced pressure, the residual was purified by column chromatography (pretreated with PE/Et_3_N 10:1) with PE/Et_3_N 20:1 as the eluent to afford diaryldiazomethanes **13a**- **f** as purple solid or oil.

### General Procedure for the Synthesis of Allenes via *N*-Tosylhydrazones (GP4):^28^

Under argon atmosphere, the alkyne (1 equiv) was added to a mixture of CuI (0.2 equiv), Li*t*OBu (3.5 equiv), and *N*-tosylhydrazone (2.2 equiv) in 1,4-dioxane (5 mL/0.4 mmol). The solution was stirred at 90 °C for 1 h, and the progress of the reaction was monitored by TLC. Upon completion of the reaction, the mixture was cooled to rt and was filtered through a short silica gel column eluting with EtOAc. The solvent was removed in vacuum to leave a crude product mixture, which was purified by column chromatography to afford the pure allene.

### General Procedure for the Cu^I^-Catalyzed Cross-Coupling of Diaryldiazomethanes and Terminal Alkynes (GP5):^30^

Under nitrogen atmosphere, diaryldiazomethane (1.0 equiv) was added to a mixture of CuI (0.2 equiv), *i*-Pr_2_NH (1.1 equiv) and terminal alkyne (1.0 equiv) in 1,4-dioxane (1.0 mL). The solution was stirred at 30 °C for 1 h and the progress of the reaction was monitored by TLC. Upon completion of the reaction, indicated by a color change from purple to yellow/brown, the reaction mixture was cooled down to room temperature and filtered through a short pad of aluminum oxide by using EtOAc as the eluent. The solvent was removed in vacuum to leave a crude mixture, which was purified by column chromatography to afford the desired allenic product.

### General Procedure for the Ester Hydrolysis with NaOH (GP6):^62^

The ester was dissolved in EtOH (5.0 mL/mmol) and 2N NaOH (1.5 mL/mmol, 3 equiv) was added. The mixture was stirred at rt and the reaction progress monitored by TLC (EtOAc). The solvent was then completely removed under reduced pressure. Phosphate buffer (pH 7) was then added to the solid residue until the pH was adjusted to 7 (indicator paper). After freeze drying, Et_2_O was added to the solid residue and the resulting suspension was filtrated. Subsequently, the filter cake was washed several times with Et_2_O. The solvent was then completely removed under reduced pressure and the remaining oil was dissolved in water and freeze dried, to obtain the free amino acid as white to yellow amorphous powder.

### General Procedure for the Ester Hydrolysis with Ba(OH)_2_ (GP7):^63^

The ester (1 equiv) was dissolved in EtOH/H_2_O (2:1) and Ba(OH)_2_x8H_2_O (2 equiv) was added. The suspension was then stirred at rt overnight. After completion of the reaction CO_2_ was bubbled through the suspension until no further BaCO_3_ precipitated. The suspension was then filtered through a syringe filter (25 mm) and the filtrate was purified by RP-MPLC. After freeze drying the corresponding free amino acid could be obtained as white to yellow amorphous powder.

### Ethyl 1-(prop-2-yn-1-yl)-(*3S*)-piperidine-3-carboxylate [(*S*)-12]

GP1 was followed applying propargyl bromide solution (80 wt% in xylene, 0.56 mL, 5.0 mmol), ethyl (*S*)- piperidine-3-carboxylate (943 mg, 6.0 mmol), acetone (10 mL), Na_2_CO_3_ (1.33 g, 12.5 mmol) and NaI (375 mg, 2.50 mmol). The crude product was purified by column chromatography (PE/EtOAc 8:2) to afford the desired alkyne *S*-**9** as pale yellow oil (973 mg, >99%): *R*_f_=0.18 (PE/EtOAc 8:2); [*α*]_D_^22^=-7.2 (c=1.76 g/100 mL in chloroform); ^1^H NMR (400 MHz, CD_2_Cl_2_): *δ*=1.23 (t, *J*=7.1 Hz, 3H, OCH_2_C*H_3_*), 1.41 (qd, *J*=11.7/3.9 Hz, 1H, NCH_2_CHC*H_ax_*H_eq_), 1.49–1.62 (m, 1H, NCH_2_C*H_ax_*H_eq_CH_2_CH), 1.69–1.78 (m, 1H, NCH_2_CH_ax_*H_eq_*CH_2_CH), 1.83–1.94 (m, 1H, NCH_2_CHCH_ax_*H_eq_*), 2.20 (td, *J*=10.9/3.1 Hz, 1H, NC*H_ax_*H_eq_CH_2_CH_2_CH), 2.27 (t, *J*=2.4 Hz, 1H, NCH_2_C≡C*H*), 2.34 (t, *J*=10.5 Hz, 1H, NC*H_ax_*H_eq_CHCH_2_), 2.53 (tt, *J*=10.4/3.9 Hz, 1H, NCH_2_C*H_ax_*CH_2_), 2.71 (dt_br_, *J*=11.2/4.2 Hz, 1H, NCH_ax_*H_eq_*CH_2_CH_2_CH), 2.94 (dd_br_, *J*=10.9/4.0 Hz, 1H, NCH_ax_*H_eq_*CHCH_2_), 3.29 (d, *J*=2.4 Hz, 2H, NC*H_2_*C≡CH), 4.09 (q, *J*=7.1 Hz, 2H, OC*H_2_*CH_3_); ^13^C NMR (101 MHz, CD_2_Cl_2_): *δ* = 14.6, 25.1, 27.0, 42.4, 47.7, 52.8, 54.8, 60.8, 73.2, 79.6, 174.4; IR (film): ṽ=3291, 2980, 2941, 2860, 2807, 1731, 1468, 1450, 1393, 1368, 1348, 1311, 1262, 1223, 1183, 1153, 1134, 1095, 1047, 1031, 1003, 979, 956, 936, 900, 864, 792, 625 cm^-1^; HRMS-ESI *m/z* [*M*+H]^+^ calcd for C_11_H_17_NO_2_: 196.1338, found: 196.1331.

### *rac*-[Ethyl 1-(but-3-yn-1-yl)piperidine-3-carboxylate]^41^ (*rac*-15)

GP1 was followed applying 4-bromo-1-butin (1.00 eq, 1.94 mL, 20.0 mmol), ethyl nipecotate (1.20 eq, 3.74 mL, 24.0 mmol), acetone (40 mL), Na_2_CO_3_ (2.50 eq, 5.30 g, 50.0 mmol) and NaI (0.50 eq, 1.50 g, 10.0 mmol). The crude product was purified by column chromatography (PE/EtOAc 8:2) to afford the desired alkyne *rac*-**15** as pale yellow oil (3.22 g, 77%): *R*_f_=0.20 (PE/EtOAc 8:2); ^1^H NMR (500 MHz, CDCl_3_): *δ*=1.25 (t, *J*=7.1 Hz, 3H, OCH_2_C*H_3_*), 1.38–1.51 (m, 1H, NCH_2_CHC*H_ax_*H_eq_), 1.51–1.63 (m, 1H, NCH_2_C*H_ax_*H_eq_CH_2_CH), 1.68–1.77 (m, 1H, NCH_2_CH_ax_*H_eq_*CH_2_CH), 1.88–1.96 (m, 1H, NCH_2_CHCH_ax_*H_eq_*), 1.97 (t, *J*=2.7 Hz, 1H, NCH_2_CH_2_CC*H*), 2.07 (td, *J*=11.0/3.1 Hz, 1H, NC*H_ax_*H_eq_CH_2_CH_2_CH), 2.23 (t, *J*=10.6 Hz, 1H, NC*H_ax_*H_eq_CHCH_2_), 2.34–2.41 (m, 2H, NCH_2_C*H_2_*CCH), 2.55 (tt, *J*=10.6/3.9 Hz, 1H, NCH_2_C*H_ax_*CH_2_), 2.57–2.66 (m, 2H, NC*H_2_*CH_2_CCH), 2.73–2.81 (m, 1H, NCH_ax_*H_eq_*CH_2_CH_2_CH), 2.94–3.02 (m, 1H, NCH_ax_*H_eq_*CHCH_2_), 4.13 (q, *J*=7.1 Hz, 2H, OC*H_2_*CH_3_); ^13^C NMR (126 MHz, CDCl_3_): *δ*=14.2 (1C, OCH_2_*C*H_3_), 16.7 (1C, NCH_2_*C*H_2_CCH), 24.5 (1C, NCH_2_*C*H_2_CH_2_CH), 26.9 (1C, NCH_2_CH_2_*C*H_2_CH), 41.8 (1C, NCH_2_CH_2_CH_2_*C*H), 53.4 (1C, N*C*H_2_CH_2_CH_2_CH), 55.1 (1C, N*C*H_2_CHCH_2_), 57.3 (1C, N*C*H_2_CH_2_CCH), 60.3 (1C, O*C*H_2_CH_3_), 69.0 (1C, NCH_2_CH_2_C*C*H), 82.8 (1C, NCH_2_CH_2_*C*CH), 174.1 (1C, *C*O); IR (film): ṽ=3297, 2945, 2856, 2812, 2118, 1731, 1469, 1447, 1371, 1311, 1273, 1214, 1181, 1154, 1134, 1102, 1033 cm^-1^; HRMS-ESI *m/z* [*M*+H]^+^ calcd for C_12_H_19_NO_2_: 210.1494, found: 210.1489.

### *rac*-[Ethyl 1-(4,4-diphenylbuta-2,3-dien-1-yl)piperidine-3-carboxylate] (*rac*-14a)

GP5 was followed using alkyne *rac*-**12**^2626^ (98 mg, 0.50 mmol), CuI (19 mg, 0.10 mmol), *i*-Pr_2_NH (0.08 mL, 0.55 mmol) and (diazomethylene)dibenzene (97 mg, 0.50 mmol) in 1,4-dioxane (2.5 mL). After purification by column chromatography (PE/EtOAc 8:2) pure *rac*-**14a** was afforded as yellow viscous oil (167 mg, 92%): *R*_f_=0.18 (PE/EtOAc 8:2); ^1^H NMR (400 MHz, CD_2_Cl_2_): *δ*=1.21 (t, *J*=7.1 Hz, 3H, OCH_2_C*H*_3_), 1.42 (qd, *J*=11.5/4.1 Hz, 1H, NCH_2_CHC*H_ax_*H_eq_), 1.47–1.62 (m, 1H, NCH_2_C*H_ax_*H_eq_CH_2_CH), 1.65–1.75 (m, 1H, NCH_2_CH_ax_*H_eq_*CH_2_CH), 1.83–1.94 (m, 1H, NCH_2_CHCH_ax_*H_eq_*), 2.10 (td, *J*=10.9/3.0 Hz, 1H, NC*H_ax_*H_eq_CH_2_CH_2_CH), 2.25 (t, *J*=10.5 Hz, 1H, NC*H_ax_*H_eq_CHCH_2_), 2.52 (tt, *J*=10.4/3.9 Hz, 1H, NCH_2_C*H_ax_*CH_2_), 2.73–2.85 (m, 1H, NCH_ax_*H_eq_*CH_2_CH_2_CH), 3.02 (dd_br_, *J*=11.2/3.6 Hz, 1H, NCH_ax_*H_eq_*CHCH_2_), 3.23 (d, *J*=7.0 Hz, 2H, NC*H_2_*CHCC), 4.07 (q, *J*=7.1 Hz, 2H, OC*H_2_*CH_3_), 5.72 (t, *J*=7.0 Hz, 1 H, NCH_2_C*H*CC), 7.24–7.31 (m, 2H, ArH), 7.33 (d, *J*=4.5 Hz, 8 H, ArH); ^13^C NMR (101 MHz, CD_2_Cl_2_): *δ*=14.6 (1C, OCH_2_*C*H_3_), 25.2 (1C, NCH_2_*C*H_2_CH_2_CH), 27.4 (1C, NCH_2_CH_2_*C*H_2_CH), 42.6 (1C, NCH_2_CH_2_CH_2_*C*H), 53.8 (1C, N*C*H_2_CH_2_CH_2_CH), 55.8 (1C, N*C*H_2_CHCH_2_), 58.4 (1C, N*C*H_2_CHCC), 60.7 (1C, O*C*H_2_CH_3_), 91.8 (1C, NCH_2_*C*HCC), 110.4 (1C, NCH_2_CHC*C*), 127.7 (2C, ArC), 128.9 (4C, ArC), 129.0 (2C, ArC), 129.0 (2C, ArC), 137.4 (1C, ArC_q_), 137.4 (1C, ArC_q_), 174.5 (1C, *C*O), 206.7 (1C, NCH_2_CH*C*C); IR (film): ṽ=3057, 3026, 2939, 2866, 2807, 1943, 1731, 1597, 1492, 1466, 1451, 1442, 1367, 1310, 1223, 1180, 1151, 1133, 1095, 1074, 1030, 998, 921, 902, 863, 768, 695 cm^-1^; HRMS-ESI *m/z* [*M*+H]^+^ calcd for C_24_H_27_NO_2_: 362.2120, found: 362.2113.

### *rac*-[Ethyl 1-(4,4-di-*p*-tolylbuta-2,3-dien-1-yl)piperidine-3-carboxylate] (*rac*-14b)

GP5 was followed using alkyne *rac*-**12**^26^ (98 mg, 0.50 mmol), CuI (19 mg, 0.10 mmol), *i*-Pr_2_NH (0.08 mL, 0.55 mmol) and 4,4’-(diazomethylene)bis(methylbenzene) (111 mg, 0.50 mmol) in 1,4-dioxane (2.5 mL) for 30 min. After purification by column chromatography (PE/EtOAc 8:2) pure *rac*-**14b** was afforded as yellow viscous oil (103 mg, 52%): *R*_f_=0.27 (PE/EtOAc 8:2); ^1^H NMR (400 MHz, C_2_D_2_Cl_4_): *δ*=1.15 (t, *J*=7.1 Hz, 3H, OCH_2_C*H*_3_), 1.33 (qd, *J*=11.8/3.8 Hz, 1H, NCH_2_CHC*H_ax_*H_eq_), 1.41–1.55 (m, 1H, NCH_2_C*H_ax_*H_eq_CH_2_CH), 1.58–1.71 (m, 1H, NCH_2_CH_ax_*H_eq_*CH_2_CH), 1.76–1.93 (m, 1H, NCH_2_CHCH_ax_*H_eq_*), 2.02 (t_br_, *J*=10.9 Hz, 1H, NC*H_ax_*H_eq_CH_2_CH_2_CH), 2.16 (t, *J*=10.6 Hz, 1H, NC*H_ax_*H_eq_CHCH_2_), 2.28 (s, 6H, C*H_3_*), 2.39–2.62 (m, 1H, NCH_2_C*H_ax_*CH_2_), 2.76 (d_br_, *J*=7.8 Hz, 1H, NCH_ax_*H_eq_*CH_2_CH_2_CH), 2.99 (d_br_, *J*=10.0 Hz, 1H, NCH_ax_*H_eq_*CHCH_2_), 3.16 (d, *J*=7.1 Hz, 2H, NC*H_2_*CHCC), 4.02 (q, *J*=7.1 Hz, 2H, OC*H_2_*CH_3_), 5.62 (t, *J*=7.1 Hz, 1H, NCH_2_C*H*CC), 7.07 (d, *J*=8.2 Hz, 4H, ArH), 7.15 (d, *J*=8.2 Hz, 4H, ArH); ^13^C NMR (101 MHz, C_2_D_2_Cl_4_): *δ*=14.6 (1C, OCH_2_*C*H_3_), 21.6 (2C, *C*H_3_), 24.9 (1C, NCH_2_*C*H_2_CH_2_CH), 27.1 (1C, NCH_2_CH_2_*C*H_2_CH), 42.1 (1C, NCH_2_CH_2_CH_2_*C*H), 53.4 (1C, N*C*H_2_CH_2_CH_2_CH), 55.2 (1C, N*C*H_2_CHCH_2_), 58.2 (1C, N*C*H_2_CHCC), 60.7 (1C, O*C*H_2_CH_3_), 91.0 (1C, NCH_2_*C*HCC), 109.8 (1C, NCH_2_CHC*C*), 128.7 (2C, ArC), 128.7 (2C, ArC), 129.4 (4C, ArC), 134.0 (2C, ArC_q_), 137.2 (2C, ArC_q_), 174.5 (1C, *C*O), 206.2 (1C, NCH_2_CH*C*C); IR (film): ṽ=3023, 2940, 2866, 2804, 2237, 1941, 1731, 1509, 1466, 1451, 1368, 1310, 1278, 1223, 1181, 1151, 1133, 1095, 1031, 1005, 949, 911, 887, 863, 822, 781, 745, 720, 702 cm^-1^; HRMS-ESI *m/z* [*M*+H]^+^ calcd for C_26_H_31_NO_2_: 390.2433, found: 390.2424.

### *rac*-{Ethyl 1-[4,4-bis(4-fluorophenyl)buta-2,3-dien-1-yl]piperidine-3-carboxylate} (*rac*-14c)

GP5 was followed using alkyne *rac*-**12**^26^ (98 mg, 0.50 mmol), CuI (19 mg, 0.10 mmol), *i*-Pr_2_NH (0.08 mL, 0.55 mmol) and 4,4’-(diazomethylene)bis(fluorobenzene) (115 mg, 0.500 mmol) in 1,4-dioxane (2.5 mL). After purification by column chromatography (PE/EtOAc 8:2) pure *rac*-**14c** was afforded as pale yellow oil (192 mg, 97%): *R*_f_=0.43 (PE/EtOAc 7:3); ^1^H NMR (400 MHz, C_2_D_2_Cl_4_): *δ*=1.14 (t, *J*=7.1 Hz, 3H, OCH_2_C*H*_3_), 1.32 (qd, *J*=11.9/3.9 Hz, 1H, NCH_2_CHC*H_ax_*H_eq_), 1.40–1.54 (m, 1H, NCH_2_C*H_ax_*H_eq_CH_2_CH), 1.55–1.70 (m, 1H, NCH_2_CH_ax_*H_eq_*CH_2_CH), 1.78–1.90 (m, 1H, NCH_2_CHCH_ax_*H_eq_*), 2.00 (td, *J*=11.1/3.1 Hz, 1H, NC*H_ax_*H_eq_CH_2_CH_2_CH), 2.14 (t, *J*=10.6 Hz, 1H, NC*H_ax_*H_eq_CHCH_2_), 2.45 (tt, *J*=10.6/3.8 Hz, 1H, NCH_2_C*H_ax_*CH_2_), 2.70 (d_br_, *J*=11.2 Hz, 1H, NCH_ax_*H_eq_*CH_2_CH_2_CH), 2.93 (dd_br_, *J*=11.1/3.4 Hz, 1H, NCH_ax_*H_eq_*CHCH_2_), 3.15 (d, *J*=7.1 Hz, 2H, NC*H_2_*CHCC), 4.01 (q, *J*=7.1 Hz, 2H, OC*H_2_*CH_3_), 5.65 (t, *J*=7.1 Hz, 1H, NCH_2_C*H*CC), 6.90–7.02 (m, 4H, FCC*H*), 7.18–7.25 (m, 4H, FCCHC*H*); ^13^C NMR (101 MHz, C_2_D_2_Cl_4_): *δ*=14.6 (1C, OCH_2_*C*H_3_), 24.9 (1C, NCH_2_*C*H_2_CH_2_CH), 27.1 (1C, NCH_2_CH_2_*C*H_2_CH), 42.1 (1C, NCH_2_CH_2_CH_2_*C*H), 53.4 (1C, N*C*H_2_CH_2_CH_2_CH), 55.2 (1C, N*C*H_2_CHCH_2_), 58.0 (1C, N*C*H_2_CHCC), 60.7 (1C, O*C*H_2_CH_3_), 91.6 (1C, NCH_2_*C*HCC), 108.4 (1C, NCH_2_CHC*C*), 115.7 (d*_CF_*, *^2^J_CF_*=21.5 Hz, 4C, FC*C*H), 130.2 (d*_CF_*, *^3^J_CF_*=3.9 Hz, 2C, FCCH*C*H), 130.3 (d*_CF_*, *^3^J_CF_*=3.9 Hz, 2C, FCCH*C*H), 132.8 (d*_CF_*, *^4^J_CF_*=1.8 Hz, 1C, FCCHCH*C*), 132.9 (d*_CF_*, *^4^J_CF_*=1.8 Hz, 1C, FCCHCH*C*), 162.27 (d*_CF_*, *^1^J_CF_*=246.5 Hz, 2C, F*C*), 174.4 (1C, *C*O), 206.2 (1C, NCH_2_CH*C*C); ^19^F NMR (376 MHz, C_2_D_2_Cl_4_): *δ*=-114.59–-114.33 (m); IR (film): ṽ= 2940, 2803, 1941, 1731, 1601, 1505, 1469, 1455, 1372, 1298, 1281, 1223, 1180, 1155, 1133, 1094, 1030, 911, 838, 800, 723 cm^-1^; HRMS-ESI *m/z* [*M*+H]^+^ calcd for C_24_H_25_F_2_NO_2_: 398.1932, found: 398.1919.

### *rac*-{Ethyl 1-[4,4-bis(4-chlorophenyl)buta-2,3-dien-1-yl]piperidine-3-carboxylate} (*rac*-14d)

GP5 was followed using alkyne *rac*-**12**^26^ (98 mg, 0.50 mmol), CuI (19 mg, 0.10 mmol), *i*-Pr_2_NH (0.08 mL, 0.55 mmol) and 4,4’- (diazomethylene)bis(chlorobenzene) (203 mg, crude compound) in 1,4-dioxane (2.5 mL). After purification by column chromatography (PE/EtOAc 8:2) pure *rac*-**14d** was afforded as yellow viscous oil (204 mg, 95%): *R*_f_=0.57 (PE/EtOAc 6:4); ^1^H NMR (400 MHz, CD_2_Cl_2_): *δ*=1.21 (t, *J*=7.1 Hz, 3H, OCH_2_C*H*_3_), 1.41 (qd, *J*=11.3/3.7 Hz, 1H, NCH_2_CHC*H_ax_*H_eq_), 1.47–1.60 (m, 1H, NCH_2_C*H_ax_*H_eq_CH_2_CH), 1.65–1.75 (m, 1H, NCH_2_CH_ax_*H_eq_*CH_2_CH), 1.84–1.94 (m, 1H, NCH_2_CHCH_ax_*H_eq_*), 2.09 (td, *J*=10.8/3.0 Hz, 1H, NC*H_ax_*H_eq_CH_2_CH_2_CH), 2.23 (t, *J*=10.5 Hz, 1H, NC*H_ax_*H_eq_CHCH_2_), 2.50 (tt, *J*=10.3/3.9 Hz, 1H, NCH_2_C*H_ax_*CH_2_), 2.71–2.80 (m, 1H, NCH_ax_*H_eq_*CH_2_CH_2_CH), 2.98 (dd_br_, *J*=11.2/3.6 Hz, 1H, NCH_ax_*H_eq_*CHCH_2_), 3.22 (dd, *J*=7.0/1.6 Hz, 2H, NC*H_2_*CHCC), 4.08 (q, *J*=7.1 Hz, 2H, OC*H_2_*CH_3_), 5.75 (t, *J*=7.0 Hz, 1H, NCH_2_C*H*CC), 7.23–7.29 (m, 4H, ArH), 7.29–7.35 (m, 4H, ArH); ^13^C NMR (101 MHz, CD_2_Cl_2_): *δ*=14.6 (1C, OCH_2_*C*H_3_), 25.2 (1C, NCH_2_*C*H_2_CH_2_CH), 27.4 (1C, NCH_2_CH_2_*C*H_2_CH), 42.5 (1C, NCH_2_CH_2_CH_2_*C*H), 53.8 (1C, N*C*H_2_CH_2_CH_2_CH), 55.7 (1C, N*C*H_2_CHCH_2_), 58.1 (1C, N*C*H_2_CHCC), 60.7 (1C, O*C*H_2_CH_3_), 92.6 (1C, NCH_2_*C*HCC), 108.8 (1C, NCH_2_CHC*C*), 129.1 (4C, ArC), 130.2 (2C, ArC), 130.3 (2C, ArC), 133.5 (1C, ArC_q_), 133.6 (1C, ArC_q_), 135.6 (1C, ArC_q_), 135.7 (1C, ArC_q_), 174.4 (1C, *C*O), 206.6 (1C, NCH_2_CH*C*C); IR (film): ṽ=2941, 2867, 2801, 2238, 1941, 1728, 1590, 1489, 1467, 1452, 1415, 1396, 1368, 1309, 1274, 1223, 1182, 1152, 1134, 1091, 1047, 1030, 1014, 949, 909, 887, 832, 746, 702 cm^-1^; HRMS-ESI *m/z* [*M*+H]^+^ calcd for C_24_H_25_Cl_2_NO_2_: 430.1341, found: 430.1338.

### *rac*-{Ethyl 1-[4,4-bis(4-methoxyphenyl)buta-2,3-dien-1-yl]piperidine-3- carboxylate} (*rac*-14e)

GP5 was followed using alkyne *rac*-**12**^26^ (98 mg, 0.50 mmol), CuI (19 mg, 0.10 mmol), *i*-Pr_2_NH (0.080 mL, 0.55 mmol) and 4,4’- (diazomethylene)bis(methoxybenzene) (650 mg, crude compound) in 1,4-dioxane (2.5 mL). After purification by column chromatography (PE/EtOAc 7:3) pure ***rac*-14e** was afforded as yellow viscous oil (148 mg, 70%): *R*_f_=0.34 (PE/EtOAc 6:4); ^1^H NMR (400 MHz, C_2_D_2_Cl_4_): *δ*=1.22 (t, *J*=7.1 Hz, 3H, OCH_2_C*H*_3_), 1.40 (qd, *J*=12.1/3.9 Hz, 1H, NCH_2_CHC*H_ax_*H_eq_), 1.49–1.63 (m, 1H, NCH_2_C*H_ax_*H_eq_CH_2_CH), 1.66–1.78 (m, 1H, NCH_2_CH_ax_*H_eq_*CH_2_CH), 1.87–1.98 (m, 1H, NCH_2_CHCH_ax_*H_eq_*), 2.02–2.15 (m, 1H, NC*H_ax_*H_eq_CH_2_CH_2_CH), 2.24 (t, *J*=10.7 Hz, 1H, NC*H_ax_*H_eq_CHCH_2_), 2.47–2.66 (m, 1H, NCH_2_C*H_ax_*CH_2_), 2.82 (d_br_, *J*=11.2 Hz, 1H, NCH_ax_*H_eq_*CH_2_CH_2_CH), 3.06 (d_br_, *J*=10.9 Hz, 1H, NCH_ax_*H_eq_*CHCH_2_), 3.23 (d, *J*=7.0 Hz, 2H, NC*H_2_*CHCC), 3.81 (s, 6H, OC*H_3_*), 4.09 (q, *J*=7.1 Hz, 2H, OC*H_2_*CH_3_), 5.69 (t, *J*=7.0 Hz, 1H, NCH_2_C*H*CC), 6.80–6.96 (m, 4H, ArH), 7.17–7.35 (m, 4H, ArH); ^13^C NMR (101 MHz, C_2_D_2_Cl_4_): *δ*=14.2 (1C, OCH_2_*C*H_3_), 24.4 (1C, NCH_2_*C*H_2_CH_2_CH), 26.7 (1C, NCH_2_CH_2_*C*H_2_CH), 41.7 (1C, NCH_2_CH_2_CH_2_*C*H), 52.9 (1C, N*C*H_2_CH_2_CH_2_CH), 54.8 (1C, N*C*H_2_CHCH_2_), 55.3 (2C, O*C*H_3_), 57.9 (1C, N*C*H_2_CHCC), 60.3 (1C, O*C*H_2_CH_3_), 90.4 (1C, NCH_2_*C*HCC), 108.8 (1C, NCH_2_CHC*C*), 113.7 (4C, ArC), 128.9 (2C, ArC_q_), 129.4 (2C, ArC), 129.4 (2C, ArC), 158.6 (2C, *C*OCH_3_), 174.0 (1C, *C*O), 205.6 (1C, NCH_2_CH*C*C); IR (film): ṽ=2938, 2835, 1729, 1606, 1578, 1508, 1464, 1442, 1368, 1295, 1248, 1175, 1151, 1134, 1095, 1034, 833 cm^-1^; HRMS-ESI *m/z* [*M*+H]^+^ calcd for C_26_H_31_NO_4_: 422.2331, found: 422.2328.

### *rac*-{Ethyl (*R_a_*)-1-[4-(1,1’-biphenyl-4-yl)-4-phenylbuta-2,3-dien-1-yl]-(*3R*)- piperidine-3-carboxylate} [*rac*-(*3R*,*R_a_*)-14f] and *rac*-{ethyl (*S_a_*)-1-[4-(1,1’-biphenyl-4-yl)-4-phenylbuta-2,3-dien-1-yl]-(*3R*)-piperidine-3-carboxylate} [*rac*-(*3R*,*S_a_*)-14f]

GP5 was followed using alkyne *rac*-**12**^26^ (98 mg, 0.50 mmol), CuI (19 mg, 0.10 mmol), *i*-Pr_2_NH (0.08 mL, 0.55 mmol) and 4-[diazo(phenyl)methyl]-1,1’-biphenyl (135 mg, 0.50 mmol) in 1,4-dioxane (2.5 mL) for 40 min. After purification by column chromatography (gradient elution PE/EtOAc 9:1 to PE/EtOAc 7:3) pure *rac*-(*3R*,*R_a_*)-**14f** and *rac*-(*3R*,*S_a_*)-**14f** were obtained as ∼ 1:1 mixture of racemic diastereomers as yellow viscous oil (205 mg, 94%): *R*_f_=0.19 (PE/EtOAc 8:2); ^1^H NMR (400 MHz, C_2_D_2_Cl_4_): *δ*=1.14 (t, *J*=7.1 Hz, 1.5H, OCH_2_C*H_3_*, dia1 or dia2), 1.14 (t, *J*=7.1 Hz, 1.5H, OCH_2_C*H_3_*, dia1 or dia2), 1.34 (qd, *J*=12.0/3.5 Hz, 1H, NCH_2_CHC*H_ax_*H_eq_), 1.43–1.53 (m, 1H, NCH_2_C*H_ax_*H_eq_CH_2_CH), 1.60–1.71 (m, 1H, NCH_2_CH_ax_*H_eq_*CH_2_CH), 1.78–1.91 (m, 1H, NCH_2_CHCH_ax_*H_eq_*), 2.04 (t_br_, *J*=11.1 Hz, 1H, NC*H_ax_*H_eq_CH_2_CH_2_CH), 2.19 (t, *J*=10.6 Hz, 1H, NC*H_ax_*H_eq_CHCH_2_), 2.40–2.55 (m, 1H, NCH_2_C*H_ax_*CH_2_), 2.76 (d_br_, *J*=9.1 Hz, 1H, NCH_ax_*H_eq_*CH_2_CH_2_CH), 2.99 (d_br_, *J*=11.0 Hz, 1H, NCH_ax_*H_eq_*CHCH_2_), 3.21 (d, *J*=7.0 Hz, 2H, NC*H_2_*CHCC), 4.01 (q, *J*=7.1 Hz, 1H, OC*H_2_*CH_3_, dia1 or dia2), 4.01 (q, *J*=7.1 Hz, 1H, OC*H_2_*CH_3_, dia1 or dia2), 5.71 (t, *J*=7.0 Hz, 1H, NCH_2_C*H*CC), 7.26–7.41 (m, 10H, ArH), 7.48–7.59 (m, 4H, ArH); ^13^C NMR (101 MHz, C_2_D_2_Cl_4_): *δ*=14.6 (1C, OCH_2_*C*H_3_), 24.8 (1C, NCH_2_*C*H_2_CH_2_CH), 27.1 (1C, NCH_2_CH_2_*C*H_2_CH), 42.1 (1C, NCH_2_CH_2_CH_2_*C*H), 53.3 (0.5C, N*C*H_2_CH_2_CH_2_CH, dia1 or dia2), 53.4 (0.5C, N*C*H_2_CH_2_CH_2_CH, dia1 or dia2), 55.2 (1C, N*C*H_2_CHCH_2_), 58.1 (0.5C, N*C*H_2_CHCC, dia1 or dia2), 58.1 (0.5C, N*C*H_2_CHCC, dia1 or dia2), 60.7 (1C, O*C*H_2_CH_3_), 91.3 (0.5C, NCH_2_*C*HCC, dia1 or dia2), 91.4 (0.5C, NCH_2_*C*HCC, dia1 or dia2), 109.9 (1C, NCH_2_CHC*C*), 127.2 (2C, ArC), 127.3 (2C, ArC), 127.7 (1C, ArC), 127.7 (1C, ArC), 128.8 (2C, ArC), 128.9 (1C, ArC), 128.9 (1C, ArC), 129.1 (1C, ArC), 129.1 (1C, ArC), 129.2 (2C, ArC), 135.9 (1C, ArC_q_), 136.8 (1C, ArC_q_), 140.0 (1C, ArC_q_), 140.7 (1C, ArC_q_), 174.4 (1C, *C*O), 206.6 (1C, NCH_2_CH*C*C); IR (film): ṽ=3424, 3057, 3028, 2939, 2803, 1943, 1730, 1598, 1486, 1466, 1447, 1367, 1310, 1222, 1180, 1151, 1133, 1094, 1030, 1007, 906, 841, 767, 729, 697 cm^-1^; HRMS-ESI *m/z* [*M*+H]^+^ calcd for C_30_H_31_NO_2_: 438.2433, found: 438.2421.

### *rac*-[1-(4,4-Diphenylbuta-2,3-dien-1-yl)piperidine-3-carboxylic acid] (*rac*-8a)

GP7 was followed using nipecotic acid ester *rac*-**14a** (171 mg, 0.472 mmol) and Ba(OH)_2_x8H_2_O (298 mg, 0.944 mmol) in 3.4 mL EtOH/H_2_O (2:1) overnight. After purification by RP-MPLC (MeOH/H_2_O 7:3) the desired amino acid *rac*-**8a** was afforded as pale yellow amorphous solid (43.6 mg, 28%). ^1^H NMR (500 MHz, MeOD, NaOD): *δ*=1.24 (qd, *J*=12.5/3.1 Hz, 1H, NCH_2_CHC*H_ax_*H_eq_), 1.39–1.53 (m, 1H, NCH_2_C*H_ax_*H_eq_CH_2_CH), 1.53–1.63 (m, 1H, NCH_2_CH_ax_*H_eq_*CH_2_CH), 1.80–1.95 (m, 2H, NCH_2_CHCH_ax_*H_eq_* and NC*H_ax_*H_eq_CH_2_CH_2_CH), 2.08 (t, *J*=11.1 Hz, 1H, NC*H_ax_*H_eq_CHCH_2_), 2.30 (t_br_, *J*=7.3 Hz, 1H, NCH_2_C*H_ax_*CH_2_), 2.82 (d, *J*=11.2 Hz, 1H, NCH_ax_*H_eq_*CH_2_CH_2_CH), 3.01–3.18 (m, 3H, NCH_ax_*H_eq_*CHCH_2_ and NC*H_2_*CHCC), 5.65 (t, *J*=7.3 Hz, 1H, NCH_2_C*H*CC), 7.19 (m, 10H, ArH); ^13^C NMR (126 MHz, MeOD, NaOD): *δ*=26.0 (1C, NCH_2_*C*H_2_CH_2_CH), 29.3 (1C, NCH_2_CH_2_*C*H_2_CH), 46.6 (1C, NCH_2_CH_2_CH_2_*C*H), 54.4 (1C, N*C*H_2_CH_2_CH_2_CH), 58.1 (1C, N*C*H_2_CHCH_2_), 59.1 (1C, N*C*H_2_CHCC), 91.1 (1C, NCH_2_*C*HCC), 111.4 (1C, NCH_2_CHC*C*), 128.4 (1C, ArC), 128.4 (1C, ArC), 129.4–129.6 (m, 8C, ArC), 137.9 (1C, ArC_q_), 138.0 (1C, ArC_q_), 182.6 (1C, *C*O), 207.6 (1C, NCH_2_CH*C*C); IR (KBr): ṽ=3418, 3057, 3027, 2937, 2858, 2804, 1945, 1560, 1492, 1466, 1451, 1442, 1407, 1333, 1155, 1093, 1074, 1030, 1000, 961, 922, 903, 769, 695, 630, 611 cm^-1^; HRMS-ESI *m/z* [*M*+H]^+^ calcd for C_22_H_23_NO_2_: 334.1807, found: 334.1801.

### *rac*-[1-(4,4-Di-*p*-tolylbuta-2,3-dien-1-yl)piperidine-3-carboxylic acid] (*rac*-8b)

GP7 was followed using nipecotic acid ester *rac*-**14b** (92.3 mg, 0.237 mmol) and Ba(OH)_2_x8H_2_O (150 mg, 0.474 mmol) in 1.5 mL EtOH/H_2_O (2:1) for 6 h. After purification by RP-MPLC (MeOH/H_2_O 7:3) the desired amino acid *rac*-**8b** was afforded as white amorphous solid (19.2 mg, 22%): mp: 101°C; ^1^H NMR (400 MHz, MeOD): *δ*=1.36 (qd, *J*=12.7/4.4 Hz, 1H, NCH_2_CHC*H_ax_*H_eq_), 1.51–1.65 (m, 1H, NCH_2_C*H_ax_*H_eq_CH_2_CH), 1.65–1.75 (m, 1H, NCH_2_CH_ax_*H_eq_*CH_2_CH), 1.91–2.05 (m, 2H, NCH_2_CHCH_ax_*H_eq_* and NC*H_ax_*H_eq_CH_2_CH_2_CH), 2.19 (t, *J*=11.2 Hz, 1H, NC*H_ax_*H_eq_CHCH_2_), 2.33 (s, 3H, C*H_3_*), 2.34 (s, 3H, C*H_3_*), 2.42 (tt, *J*=11.6/3.8 Hz, 1H, NCH_2_C*H_ax_*CH_2_), 2.93 (d_br_, *J*=11.4 Hz, 1H, NCH_ax_*H_eq_*CH_2_CH_2_CH), 3.13–3.30 (m, 3H, NCH_ax_*H_eq_*CHCH_2_ and NC*H_2_*CHCC), 5.71 (t, *J*=7.3 Hz, 1H, NCH_2_C*H*CC), 6.99–7.31 (m, 8H, ArH); ^13^C NMR (126 MHz, MeOD): *δ*=21.2 (2C, *C*H_3_), 26.0 (1C, NCH_2_*C*H_2_CH_2_CH), 29.3 (1C, NCH_2_CH_2_*C*H_2_CH), 46.6 (1C, NCH_2_CH_2_CH_2_*C*H), 54.4 (1C, N*C*H_2_CH_2_CH_2_CH), 58.1 (1C, N*C*H_2_CHCH_2_), 59.3 (1C, N*C*H_2_CHCC), 90.8 (1C, NCH_2_*C*HCC), 111.1 (1C, NCH_2_CHC*C*), 129.4 (2C, ArC), 129.4 (2C, ArC), 130.1 (2C, ArC), 130.1 (2C, ArC), 135.0 (1C, ArC_q_), 135.1 (1C, ArC_q_), 138.2 (1C, ArC_q_), 138.3 (1C, ArC_q_), 182.7 (1C, *C*O), 207.4 (1C, NCH_2_CH*C*C); IR (KBr): ṽ=3387, 3026, 2922, 1932, 1603, 1508, 1447, 1374, 1362, 1336, 1309, 1279, 1258, 1210, 1178, 1149, 1108, 1059, 1017, 959, 938, 912, 872, 855, 829, 820, 783, 725, 668, 647, 610, 591, 526 cm^-1^; HRMS-ESI *m/z* [*M*+H]^+^ calcd for C_24_H_27_NO_2_: 362.2120, found: 362.2112.

### *rac*-{1-[4,4-Bis(4-fluorophenyl)buta-2,3-dien-1-yl]piperidine-3-carboxylic acid} (*rac*-8c)

GP6 was followed using nipecotic acid ester *rac*-**14c** (0.425 mmol, 169 mg), EtOH (2.1 mL) and 2M NaOH (0.64 mL) for 40 min. The desired amino acid *rac*-**8c** was obtained as pale yellow amorphous solid (40.4 mg, 26%): mp: 77 °C; ^1^H NMR (400 MHz, MeOD, NaOD): *δ*=1.11 (qd, *J*=12.7/4.1 Hz, 1H, NCH_2_CHC*H_ax_*H_eq_), 1.33 (qt, *J*=13.1/4.0 Hz, 1H, NCH_2_C*H_ax_*H_eq_CH_2_CH), 1.40–1.52 (m, 1H, NCH_2_CH_ax_*H_eq_*CH_2_CH), 1.67–1.81 (m, 2H, NCH_2_CHCH_ax_*H_eq_* and NC*H_ax_*H_eq_CH_2_CH_2_CH), 1.92 (t, *J*=11.3 Hz, 1H, NC*H_ax_*H_eq_CHCH_2_), 2.16 (tt, *J*=11.8/3.8 Hz, 1H, NCH_2_C*H_ax_*CH_2_), 2.66 (d_br_, *J*=11.2 Hz, 1H, NCH_ax_*H_eq_*CH_2_CH_2_CH), 2.90–2.97 (m, 1H, NCH_ax_*H_eq_*CHCH_2_), 2.99 (dd, *J*=7.3/2.0 Hz, 2H, NC*H_2_*CHCC), 5.53 (t, *J*=7.3 Hz, 1H, NCH_2_C*H*CC), 6.76–6.92 (m, 4H, FCC*H*), 6.98–7.12 (m, 4H, FCCHC*H*); ^13^C NMR (101 MHz, MeOD, NaOD): *δ*=26.0 (1C, NCH_2_*C*H_2_CH_2_CH), 29.3 (1C, NCH_2_CH_2_*C*H_2_CH), 46.6 (1C, NCH_2_CH_2_CH_2_*C*H), 54.5 (1C, N*C*H_2_CH_2_CH_2_CH), 58.0 (1C, N*C*H_2_CHCH_2_), 59.0 (1C, N*C*H_2_CHCC), 91.6 (1C, NCH_2_*C*HCC), 109.6 (1C, NCH_2_CHC*C*), 116.4 (dd*_CF_*, *^2^J_CF_*=21.9/2.0 Hz, 4C, FC*C*H), 131.2 (dd*_CF_*, *^3^J_CF_*=8.1/3.0 Hz, 4C, FCCH*C*H), 134.0 (t*_CF_*, *^4^J_CF_*=3.8 Hz, 2C, FCCHCH*C*), 163.7 (d*_CF_*, *^1^J_CF_*=245.5 Hz, 2C, F*C*), 182.7 (1C, *C*O), 207.4 (1C, NCH_2_CH*C*C); ^19^F NMR (376 MHz, MeOD, NaOD): *δ*=-116.9, -117.0; IR (KBr): ṽ=3447, 3048, 2940, 2863, 2802, 1944, 1715, 1601, 1505, 1467, 1451, 1400, 1338, 1298, 1282, 1224, 1156, 1095, 1013, 911, 838, 800, 724, 606, 584 cm^-1^; HRMS-ESI *m/z* [*M*+H]^+^ calcd for C_22_H_21_F_2_NO_2_: 370.1619, found: 370.1610.

### *rac*-{1-[4,4-Bis(4-chlorophenyl)penta-3,4-dien-1-yl]piperidine-3-carboxylic acid} (*rac*-8d)

GP6 was followed using nipecotic acid ester *rac*-**14d** (0.470 mmol, 202 mg), EtOH (2.35 mL) and 2M NaOH (0.71 mL) for 1.5 h. The desired amino acid *rac*-**8d** was obtained as pale yellow amorphous solid (46.8 mg, 25%): mp: 91 °C; ^1^H NMR (500 MHz, MeOD, NaOD): *δ*=1.36 (qd, *J*=12.7/4.1 Hz, 1H, NCH_2_CHC*H_ax_*H_eq_), 1.58 (qt, *J*=13.0/3.8 Hz, 1H, NCH_2_C*H_ax_*H_eq_CH_2_CH), 1.67–1.76 (m, 1H, NCH_2_CH_ax_*H_eq_*CH_2_CH), 1.94–2.06 (m, 2H, NCH_2_CHCH_ax_*H_eq_* and NC*H_ax_*H_eq_CH_2_CH_2_CH), 2.18 (t, *J*=11.3 Hz, 1H, NC*H_ax_*H_eq_CHCH_2_), 2.41 (tt, *J*=11.7/3.8 Hz, 1H, NCH_2_C*H_ax_*CH_2_), 2.90 (d_br_, *J*=11.2 Hz, 1H, NCH_ax_*H_eq_*CH_2_CH_2_CH), 3.14–3.22 (m, 1H, NCH_ax_*H_eq_*CHCH_2_), 3.26 (dd, *J*=7.3/2.0 Hz, 2H, NC*H_2_*CHCC), 5.83 (t, *J*=7.3 Hz, 1H, NCH_2_C*H*CC), 7.21–7.32 (m, 4H, ArH), 7.32–7.42 (m, 4H, ArH); ^13^C NMR (126 MHz, MeOD, NaoD): *δ*=26.0 (1C, NCH_2_*C*H_2_CH_2_CH), 29.3 (1C, NCH_2_CH_2_*C*H_2_CH), 46.5 (1C, NCH_2_CH_2_CH_2_*C*H), 54.5 (1C, N*C*H_2_CH_2_CH_2_CH), 58.0 (1C, N*C*H_2_CHCH_2_), 58.7 (1C, N*C*H_2_CHCC), 92.1 (1C, NCH_2_*C*HCC), 109.7 (1C, NCH_2_CHC*C*), 129.8 (2C, ArC), 129.8 (2C, ArC), 130.9 (2C, ArC), 130.9 (2C, ArC), 134.4 (1C, ArC_q_), 134.4 (1C, ArC_q_), 136.3 (1C, ArC_q_), 136.3 (1C, ArC_q_), 182.7 (1C, *C*O), 207.5 (1C, CH_2_CH*C*C); IR (KBr): ṽ=3433, 3030, 2938, 2860, 2796, 2522, 1942, 1712, 1589, 1489, 1451, 1396, 1334, 1300, 1222, 1191, 1153, 1090, 1044, 1013, 959, 909, 868, 831, 783, 747, 722, 702, 638, 612, 535, 509, 460 cm^-1^; HRMS-ESI *m/z* [*M*+H]^+^ calcd for C_22_H_21_Cl_2_NO_2_: 402.1028, found: 402.1020.

### 1-[4,4-Bis(4-chlorophenyl)penta-3,4-dien-1-yl]-(*R*)-piperidine-3-carboxylic acid [(*R*)-8d]

A) Synthesis of starting material (*R*)-**14d**: GP5 was followed using alkyne (*R*)- **12**^25^ (98 mg, 0.50 mmol), CuI (19 mg, 0.10 mmol), *i*-Pr_2_NH (0.08 mL, 0.55 mmol) and 4,4’-(diazomethylene)bis(chlorobenzene) (132 mg, 0.50 mmol) in 1,4-dioxane (2.5 mL). After purification by column chromatography (PE/EtOAc 8:2) pure (*R*)-**14d** was afforded as yellow viscous oil (208 mg, 97%): *R*_f_=0.21 (PE/EtOAc 8:2). B) Synthesis of (*R*)-**8d**: GP6 was followed using nipecotic acid ester (*R*)-**14d** (0.480 mmol, 207 mg), EtOH (2.4 mL) and 2M NaOH (0.72 mL) for 1.5 h. The desired amino acid (*R*)-**8d** was obtained as pale yellow amorphous solid (99.0 mg, 51%): mp: 85 °C; [*α*]_D_^22^=+15.2 (c=0.90 g/100 mL in chloroform); ^1^H NMR (500 MHz, MeOD, NaOD): *δ*=1.36 (qd, *J*=12.7/3.9 Hz, 1H, NCH_2_CHC*H_ax_*H_eq_), 1.53–1.64 (m, 1H, NCH_2_C*H_ax_*H_eq_CH_2_CH), 1.66–1.75 (m, 1H, NCH_2_CH_ax_*H_eq_*CH_2_CH), 1.94–2.06 (m, 2H, NCH_2_CHCH_ax_*H_eq_* and NC*H_ax_*H_eq_CH_2_CH_2_CH), 2.18 (t, *J*=11.3 Hz, 1H, NC*H_ax_*H_eq_CHCH_2_), 2.41 (tt, *J*=11.7/3.3 Hz, 1H, NCH_2_C*H_ax_*CH_2_), 2.90 (d_br_, *J*=11.2 Hz, 1H, NCH_ax_*H_eq_*CH_2_CH_2_CH), 3.19 (d_br_, *J*=11.7 Hz, 1H, NCH_ax_*H_eq_*CHCH_2_), 3.25 (dd, *J*=7.3/1.3 Hz, 2H, NC*H_2_*CHCC), 5.83 (t, *J*=7.3 Hz, 1H, NCH_2_C*H*CC), 7.24–7.31 (m, 4H, ArH), 7.32–7.41 (m, 4H, ArH); ^13^C NMR (126 MHz, MeOD, NaOD): *δ*=26.0 (1C, NCH_2_*C*H_2_CH_2_CH), 29.3 (1C, NCH_2_CH_2_*C*H_2_CH), 46.6 (1C, NCH_2_CH_2_CH_2_*C*H), 54.5 (1C, N*C*H_2_CH_2_CH_2_CH), 58.0 (1C, N*C*H_2_CHCH_2_), 58.7 (1C, N*C*H_2_CHCC), 92.1 (1C, NCH_2_*C*HCC), 109.7 (1C, NCH_2_CHC*C*), 129.8 (2C, ArC), 129.8 (2C, ArC), 130.9 (2C, ArC), 130.9 (2C, ArC), 134.4 (1C, ArC_q_), 134.4 (1C, ArC_q_), 136.3 (1C, ArC_q_), 136.3 (1C, ArC_q_), 182.6 (1C, *C*O), 207.5 (1C, CH_2_CH*C*C); IR (KBr): ṽ=3426, 3030, 2938, 2862, 2797, 2531, 1942, 1711, 1589, 1489, 1451, 1396, 1333, 1300, 1222, 1192, 1152, 1090, 1044, 1013, 959, 909, 868, 831, 769, 747, 722, 702, 638, 612, 534, 508, 460 cm^-1^; HRMS-ESI *m/z* [*M*+H]^+^ calcd for C_22_H_21_Cl_2_NO_2_: 402.1028, found: 402.1021.

### 1-[4,4-Bis(4-chlorophenyl)penta-3,4-dien-1-yl]-(*S*)-piperidine-3-carboxylic acid [(*S*)-8d]

A) Synthesis of starting material (*S*)-**14d**: GP5 was followed using alkyne (*S*)- **12** (98 mg, 0.50 mmol), CuI (19 mg, 0.10 mmol), *i*-Pr_2_NH (0.08 mL, 0.55 mmol) and 4,4’-(diazomethylene)bis(chlorobenzene) (132 mg, 0.50 mmol) in 1,4-dioxane (2.5 mL). After purification by column chromatography (PE/EtOAc 8:2) pure (*S*)-**14d** was afforded as yellow viscous oil (207 mg, 96%): *R*_f_=0.17 (PE/EtOAc 8:2). B) Synthesis of (*S*)-**8d**: GP6 was followed using nipecotic acid ester (*S*)-**14d** (0.480 mmol, 207 mg), EtOH (2.4 mL) and 2M NaOH (0.72 mL) for 1.5 h. The desired amino acid (*S*)-**8d** was obtained as pale yellow amorphous solid (130 mg, 68%): mp: 86 °C; [*α*]_D_^22^=-18.4 (c=0.90 g/100 mL in chloroform); ^1^H NMR (400 MHz, MeOD, NaOD): *δ*=1.36 (qd, *J*=12.6/4.2 Hz, 1H, NCH_2_CHC*H_ax_*H_eq_), 1.58 (qt, *J*=12.8/4.0 Hz, 1H, NCH_2_C*H_ax_*H_eq_CH_2_CH), 1.66–1.76 (m, 1H, NCH_2_CH_ax_*H_eq_*CH_2_CH), 1.93–2.06 (m, 2H, NCH_2_CHCH_ax_*H_eq_* and NC*H_ax_*H_eq_CH_2_CH_2_CH), 2.18 (t, *J*=11.2 Hz, 1H, NC*H_ax_*H_eq_CHCH_2_), 2.41 (tt, *J*=11.6/3.7 Hz, 1H, NCH_2_C*H_ax_*CH_2_), 2.90 (d_br_, *J*=11.2 Hz, 1H, NCH_ax_*H_eq_*CH_2_CH_2_CH), 3.18 (dd_br_, *J*=11.1/3.5 Hz, 1H, NCH_ax_*H_eq_*CHCH_2_), 3.25 (dd, *J*=7.3/1.3 Hz, 2H, NC*H_2_*CHCC), 5.83 (t, *J*=7.3 Hz, 1H, NCH_2_C*H*CC), 7.24–7.31 (m, 4H, ArH), 7.31–7.40 (m, 4H, ArH); ^13^C NMR (126 MHz, MeOD, NaOD): *δ*=26.0 (1C, NCH_2_*C*H_2_CH_2_CH), 29.3 (1C, NCH_2_CH_2_*C*H_2_CH), 46.5 (1C, NCH_2_CH_2_CH_2_*C*H), 54.5 (1C, N*C*H_2_CH_2_CH_2_CH), 57.9 (1C, N*C*H_2_CHCH_2_), 58.7 (1C, N*C*H_2_CHCC), 92.1 (1C, NCH_2_*C*HCC), 109.7 (1C, NCH_2_CHC*C*), 129.8 (2C, ArC), 129.8 (2C, ArC), 130.9 (2C, ArC), 130.9 (2C, ArC), 134.4 (1C, ArC_q_), 134.4 (1C, ArC_q_), 136.3 (1C, ArC_q_), 136.3 (1C, ArC_q_), 182.7 (1C, *C*O), 207.5 (1C, CH_2_CH*C*C); IR (KBr): ṽ=3428, 3030, 2936, 2858, 2797, 2528, 1943, 1708, 1589, 1489, 1450, 1396, 1333, 1300, 1276, 1222, 1192, 1152, 1090, 1044, 1013, 959, 909, 867, 831, 770, 746, 722, 702, 670, 638, 612, 535, 509, 462 cm^-1^; HRMS-ESI *m/z* [*M*+H]^+^ calcd for C_22_H_21_Cl_2_NO_2_: 402.1028, found: 402.1022.

### *rac*-{(*R_a_*)-1-[4-(1,1’-biphenyl-4-yl)-4-phenylbuta-2,3-dien-1-yl]-(*3R*)-piperidine-3-carboxylic acid} [*rac*-(*3R*,*R_a_*)-8f] and *rac*-{(*S_a_*)-1-[4-(1,1’-biphenyl-4-yl)-4- phenylbuta-2,3-dien-1-yl]-(*3R*)-piperidine-3-carboxylic acid} [*rac*-(*3R*,*S_a_*)-8f]

GP6 was followed using nipecotic acid esters *rac*-(*3R*,*R_a_*)-**14f** and *rac*-(*3R*,*S_a_*)-**14f** (0.423 mmol, 185 mg), EtOH (2.1 mL) and 2M NaOH (0.63 mL) for 2 h. The desired amino acids *rac*-(*3R*,*R_a_*)-**8f** and *rac*-(*3R*,*S_a_*)-**8f** were obtained as ∼ 1:1 mixture of racemic diastereomers as pale yellow amorphous solid (16.2 mg, 9%): mp: 109 °C; ^1^H NMR (400 MHz, MeOD, NaOD): *δ*=1.37 (qd, *J*=12.7/4.2 Hz, 1H, NCH_2_CHC*H_ax_*H_eq_), 1.51–1.67 (m, 1H, NCH_2_C*H_ax_*H_eq_CH_2_CH), 1.67–1.79 (m, 1H, NCH_2_CH_ax_*H_eq_*CH_2_CH), 1.93–2.10 (m, 2H, NCH_2_CHCH_ax_*H_eq_* and NC*H_ax_*H_eq_CH_2_CH_2_CH), 2.21 (t, *J*=11.3 Hz, 0.5H, NC*H_ax_*H_eq_CHCH_2_, dia1 or dia2), 2.23 (t, *J*=11.3 Hz, 0.5H, NC*H_ax_*H_eq_CHCH_2_, dia1 or dia2), 2.43 (tt, *J*=11.6/3.8 Hz, 1H, NCH_2_C*H_ax_*CH_2_), 2.96 (d_br_, *J*=11.3 Hz, 1H, NCH_ax_*H_eq_*CH_2_CH_2_CH), 3.17–3.30 (m, 3H, NCH_ax_*H_eq_*CHCH_2_ and NC*H_2_*CHCC), 5.80 (t, *J*=7.3 Hz, 0.5H, NCH_2_C*H*CC, dia1 or dia2), 5.81 (t, *J*=7.3 Hz, 0.5H, NCH_2_C*H*CC, dia1 or dia2), 7.22–7.47 (m, 10H, ArH), 7.53–7.70 (m, 4H, ArH); ^13^C NMR (126 MHz, MeOD, NaOD): *δ*=26.0 (1C, NCH_2_*C*H_2_CH_2_CH), 29.3 (1C, NCH_2_CH_2_*C*H_2_CH), 46.6 (0.5C, NCH_2_CH_2_CH_2_*C*H, dia1 or dia2), 46.6 (0.5C, NCH_2_CH_2_CH_2_*C*H, dia1 or dia2), 54.4 (0.5C, N*C*H_2_CH_2_CH_2_CH, dia1 or dia2), 54.5 (0.5C, N*C*H_2_CH_2_CH_2_CH, dia1 or dia2), 58.0 (0.5C, N*C*H_2_CHCH_2_, dia1 or dia2), 58.1 (0.5C, N*C*H_2_CHCH_2_, dia1 or dia2), 59.1 (0.5C, N*C*H_2_CHCC, dia1 or dia2), 59.2 (0.5C, N*C*H_2_CHCC, dia1 or dia2), 91.2 (0.5C, NCH_2_*C*HCC, dia1 or dia2), 91.3 (0.5C, NCH_2_*C*HCC, dia1 or dia2), 111.1 (1C, NCH_2_CHC*C*), 127.9 (2C, ArC), 128.0 (1C, ArC), 128.0 (1C, ArC), 128.4 (0.5C, ArC, dia1 or dia2), 128.4 (0.5C, ArC, dia1 or dia2), 128.5 (0.5C, ArC, dia1 or dia2), 128.5 (0.5C, ArC, dia1 or dia2), 129.6 (1C, ArC), 129.6 (1C, ArC), 129.6 (1C, ArC), 129.6 (1C, ArC), 129.9 (1C, ArC), 129.9 (2C, ArC), 129.9 (1C, ArC), 136.9 (0.5C, ArC_q_, dia1 or dia2), 136.9 (0.5C, ArC_q_, dia1 or dia2), 137.9 (0.5C, ArC_q_, dia1 or dia2), 137.9 (0.5C, ArC_q_, dia1 or dia2), 141.5 (0.5C, ArC_q_, dia1 or dia2), 141.5 (0.5C, ArC_q_, dia1 or dia2), 141.9 (1C, ArC_q_), 182.7 (1C, *C*O), 207.7 (0.5C, NCH_2_CH*C*C, dia1 or dia2), 207.7 (0.5C, NCH_2_CH*C*C, dia1 or dia2); IR (KBr): ṽ=3423, 3055, 3027, 2936, 2857, 2800, 1942, 1708, 1597, 1578, 1486, 1466, 1447, 1405, 1334, 1309, 1224, 1191, 1155, 1133, 1094, 1074, 1040, 1007, 963, 907, 841, 767, 729, 697, 630 cm^-1^; HRMS-ESI *m/z* [*M*+H]^+^ calcd for C_28_H_27_NO_2_: 410.2120, found: 410.2110.

### *rac*-[Ethyl 1-(5,5-diphenylpenta-3,4-dien-1-yl)piperidine-3-carboxylate] (*rac*-16a)

GP5 was followed using alkyne *rac*-**15** (105 mg, 0.500 mmol), CuI (19 mg, 0.10 mmol), *i*-Pr_2_NH (0.08 mL, 0.55 mmol) and (diazomethylene)dibenzene (97 mg, 0.50 mmol) in 1,4-dioxane (2.5 mL). After purification by column chromatography (PE/EtOAc 8:2) pure *rac*-**16a** was afforded as yellow viscous oil (151 mg, 80%): *R*_f_=0.28 (PE/EtOAc 7:3); ^1^H NMR (400 MHz, CD_2_Cl_2_): *δ*=1.22 (t, *J*=7.1 Hz, 3H, OCH_2_C*H*_3_), 1.33–1.44 (m, 1H, NCH_2_CHC*H_ax_*H_eq_), 1.45–1.56 (m, 1H, NCH_2_C*H_ax_*H_eq_CH_2_CH), 1.61–1.72 (m, 1H, NCH_2_CH_ax_*H_eq_*CH_2_CH), 1.82–1.92 (m, 1H, NCH_2_CHCH_ax_*H_eq_*), 1.99 (td, *J*=10.9/2.9 Hz, 1H, NC*H_ax_*H_eq_CH_2_CH_2_CH), 2.15 (t, *J*=10.6 Hz, 1H, NC*H_ax_*H_eq_CHCH_2_), 2.36 (q, *J*=6.8 Hz, 2H, NCH_2_C*H_2_*CHCC), 2.41–2.59 (m, 3H, NCH_2_C*H_ax_*CH_2_ and NC*H_2_*CH_2_CHCC), 2.68–2.81 (m, 1H, NCH_ax_*H_eq_*CH_2_CH_2_CH), 2.96 (d_br_, *J*=11.1 Hz, 1H, NCH_ax_*H_eq_*CHCH_2_), 4.07 (q, *J*=7.1 Hz, 2H, OC*H_2_*CH_3_), 5.72 (t, *J*=6.8 Hz, 1H, NCH_2_CH_2_C*H*CC), 7.15–7.27 (m, 2H, ArH), 7.27–7.41 (m, 8H, ArH); ^13^C NMR (101 MHz, CD_2_Cl_2_): *δ*=14.4 (1C, OCH_2_*C*H_3_), 25.1 (1C, NCH_2_*C*H_2_CH_2_CH), 27.1 (1C, NCH_2_*C*H_2_CHCC or NCH_2_CH_2_*C*H_2_CH), 27.4 (1C, NCH_2_*C*H_2_CHCC or NCH_2_CH_2_*C*H_2_CH), 42.3 (1C, NCH_2_CH_2_CH_2_*C*H), 54.1 (1C, N*C*H_2_CH_2_CH_2_CH), 56.0 (1C, N*C*H_2_CHCH_2_), 58.4 (1C, N*C*H_2_CH_2_CHCC), 60.5 (1C, O*C*H_2_CH_3_), 92.9 (1C, NCH_2_CH_2_*C*HCC), 110.1 (1C, NCH_2_CH_2_CHC*C*), 127.4 (2C, ArC), 128.7 (4C, ArC), 128.8 (4C, ArC), 137.5 (2C, ArC_q_), 174.4 (1C, *C*O), 205.9 (1C, CH_2_CH*C*C); IR (film): ṽ=3057, 3025, 2940, 2854, 2806, 1943, 1731, 1597, 1492, 1467, 1452, 1442, 1370, 1310, 1273, 1210, 1178, 1151, 1100, 1031, 768, 695 cm^-1^; HRMS-ESI *m/z* [*M*+H]^+^ calcd for C_25_H_29_NO_2_: 376.2277, found: 376.2272.

### *rac*-[Ethyl 1-(5,5-di-*p*-tolylpenta-3,4-dien-1-yl)piperidine-3-carboxylate] (*rac*- 16b)

GP5 was followed using alkyne *rac*-**15** (105 mg, 0.500 mmol), CuI (19 mg, 0.10 mmol), *i*-Pr_2_NH (0.08 mL, 0.55 mmol) and 4,4’-(diazomethylene)bis(methylbenzene) (111 mg, 0.500 mmol) in 1,4-dioxane (2.5 mL) for 30 min. After purification by column chromatography (PE/EtOAc 8:2) pure *rac*-**16b** was afforded as yellow viscous oil (124 mg, 62%): *R*_f_=0.20 (PE/EtOAc 8:2); ^1^H NMR (400 MHz, C_2_D_2_Cl_4_): *δ*=1.16 (t, *J*=7.1 Hz, 3H, OCH_2_C*H*_3_), 1.32 (qd, *J*=11.9/3.8 Hz, 1H, NCH_2_CHC*H_ax_*H_eq_), 1.38–1.52 (m, 1H, NCH_2_C*H_ax_*H_eq_CH_2_CH), 1.52–1.66 (m, 1H, NCH_2_CH_ax_*H_eq_*CH_2_CH), 1.75–1.87 (m, 1H, NCH_2_CHCH_ax_*H_eq_*), 1.91 (t_br_, *J*=10.4 Hz, 1H, NC*H_ax_*H_eq_CH_2_CH_2_CH), 2.07 (t, *J*=10.6 Hz, 1H, NC*H_ax_*H_eq_CHCH_2_), 2.28 (s, 8H, NCH_2_C*H_2_*CHCC and C*H_3_*), 2.35–2.55 (m, 3H, NCH_2_C*H_ax_*CH_2_ and NC*H_2_*CH_2_CHCC), 2.69 (d_br_, *J*=11.0 Hz, 1H, NCH_ax_*H_eq_*CH_2_CH_2_CH), 2.90 (d_br_, *J*=11.0 Hz, 1H, NCH_ax_*H_eq_*CHCH_2_), 4.02 (q, *J*=7.1 Hz, 2H, OC*H_2_*CH_3_), 5.60 (t, *J*=6.6 Hz, 1H, NCH_2_CH_2_C*H*CC), 7.07 (d, *J*=8.0 Hz, 4H, ArH), 7.17 (d, *J*=8.0 Hz, 4H, ArH); ^13^C NMR (101 MHz, C_2_D_2_Cl_4_): *δ*=14.6 (1C, OCH_2_*C*H_3_), 21.6 (2C, *C*H_3_), 24.9 (1C, NCH_2_*C*H_2_CH_2_CH), 26.9 (1C, NCH_2_*C*H_2_CHCC or NCH_2_CH_2_*C*H_2_CH), 27.3 (1C, NCH_2_*C*H_2_CHCC or NCH_2_CH_2_*C*H_2_CH), 42.1 (1C, NCH_2_CH_2_CH_2_*C*H), 53.8 (1C, N*C*H_2_CH_2_CH_2_CH), 55.6 (1C, N*C*H_2_CHCH_2_), 58.3 (1C, N*C*H_2_CH_2_CHCC), 60.7 (1C, O*C*H_2_CH_3_), 92.5 (1C, NCH_2_CH_2_*C*HCC), 109.7 (1C, NCH_2_CH_2_CHC*C*), 128.6 (2C, ArC), 128.6 (2C, ArC), 129.4 (4C, ArC), 134.3 (2C, ArC_q_), 137.0 (2C, ArC_q_), 174.6 (1C, *C*O), 205.5 (1C, NCH_2_CH_2_CH*C*C); IR (film): ṽ=3022, 2939, 2857, 2805, 1940, 1731, 1508, 1468, 1445, 1370, 1308, 1275, 1210, 1179, 1151, 1133, 1100, 1033, 910, 862, 822, 720 cm^-1^; HRMS-ESI *m/z* [*M*+H]^+^ calcd for C_27_H_33_NO_2_: 404.2590, found: 404.2579.

### *rac*-{Ethyl 1-[5,5-bis(4-fluorophenyl)penta-3,4-dien-1-yl]piperidine-3- carboxylate} (*rac*-16c)

GP5 was followed using alkyne *rac*-**15** (105 mg, 0.500 mmol), CuI (19 mg, 0.10 mmol), *i*-Pr_2_NH (0.08 mL, 0.55 mmol) and 4,4’-(diazomethylene)bis(fluorobenzene) (115 mg, 0.500 mmol) in 1,4-dioxane (2.5 mL). After purification by column chromatography (PE/EtOAc 8:2) pure *rac*-**16c** was afforded as pale yellow oil (194 mg, 94%): *R*_f_=0.21 (PE/EtOAc 7:3); ^1^H NMR (400 MHz, C_2_D_2_Cl_4_): *δ*=1.15 (t, *J*=7.1 Hz, 3H, OCH_2_C*H*_3_), 1.23–1.49 (m, 2H, NCH_2_CHC*H_ax_*H_eq_ and NCH_2_C*H_ax_*H_eq_CH_2_CH), 1.54–1.68 (m, 1H, NCH_2_CH_ax_*H_eq_*CH_2_CH), 1.76–1.86 (m, 1H, NCH_2_CHCH_ax_*H_eq_*), 1.91 (td, *J*=10.6/2.3 Hz, 1H, NC*H_ax_*H_eq_CH_2_CH_2_CH), 2.07 (t, *J*=10.6 Hz, 1H, NC*H_ax_*H_eq_CHCH_2_), 2.29 (q, *J*=7.0 Hz, 2H, NCH_2_C*H_2_*CHCC), 2.34–2.52 (m, 3H, NCH_2_C*H_ax_*CH_2_ and NC*H_2_*CH_2_CHCC), 2.67 (d_br_, *J*=10.9 Hz, 1H, NCH_ax_*H_eq_*CH_2_CH_2_CH), 2.88 (d_br_, *J*=10.3 Hz, 1H, NCH_ax_*H_eq_*CHCH_2_), 4.01 (q, *J*=7.1 Hz, 2H, OC*H_2_*CH_3_), 5.64 (t, *J*=6.6 Hz, 1H, NCH_2_CH_2_C*H*CC), 6.91–7.01 (m, 4H, FCC*H*), 7.16–7.29 (m, 4H, FCCHC*H*); ^13^C NMR (101 MHz, C_2_D_2_Cl_4_): *δ*=14.4 (1C, OCH_2_*C*H_3_), 24.7 (1C, NCH_2_*C*H_2_CH_2_CH), 26.6 (1C, NCH_2_*C*H_2_CHCC), 27.1 (1C, NCH_2_CH_2_*C*H_2_CH), 41.9 (1C, NCH_2_CH_2_CH_2_*C*H), 53.6 (1C, N*C*H_2_CH_2_CH_2_CH), 55.4 (1C, N*C*H_2_CHCH_2_), 57.9 (1C, N*C*H_2_CH_2_CHCC), 60.5 (1C, O*C*H_2_CH_3_), 93.0 (1C, NCH_2_CH_2_*C*HCC), 108.1 (1C, NCH_2_CH_2_CHC*C*), 115.4 (d*_CF_*, *^2^J_CF_*=21.4 Hz, 4C, FC*C*H), 130.0 (d*_CF_*, *^3^J_CF_*=8.0 Hz, 2C, FCCH*C*H), 130.3 (d*_CF_*, *^3^J_CF_*=8.0 Hz, 2C, FCCH*C*H), 133.0 (d*_CF_*, *^4^J_CF_*=2.8 Hz, 2C, FCCHCH*C*), 162.0 (d*_CF_*, *^1^J_CF_*=245.9 Hz, 2C, F*C*), 174.4 (1C, *C*O), 205.3 (1C, CH_2_CH*C*C); ^19^F {^1^H} NMR (376 MHz, C_2_D_2_Cl_4_): *δ*=-114.8; IR (film): ṽ=2941, 2855, 2806, 1942, 1893, 1731, 1601, 1505, 1468, 1446, 1370, 1299, 1279, 1222, 1179, 1156, 1134, 1096, 1032, 1014, 911, 838, 800, 723 cm^-1^; HRMS-ESI *m/z* [*M*+H]^+^ calcd for C_25_H_27_F_2_NO_2_: 412.2088, found: 412.2078.

### *rac*-{Ethyl 1-[5,5-bis(4-chlorophenyl)penta-3,4-dien-1-yl]piperidine-3- carboxylate} (*rac*-16d)

GP5 was followed using alkyne *rac*-**15** (105 mg, 0.500 mmol), CuI (19 mg, 0.10 mmol), *i*-Pr_2_NH (0.08 mL, 0.55 mmol) and 4,4’- (diazomethylene)bis(chlorobenzene) (303 mg, crude compound) in 1,4-dioxane (2.5 mL). After purification by column chromatography (PE/EtOAc 8:2) pure ***rac*-16d** was afforded as yellow viscous oil (203 mg, 91%): *R*_f_=0.50 (PE/EtOAc 6:4); ^1^H NMR (400 MHz, CD_2_Cl_2_): *δ*=1.22 (t, *J*=7.1 Hz, 3H, OCH_2_C*H*_3_), 1.32–1.55 (m, 2H, NCH_2_CHC*H_ax_*H_eq_ and NCH_2_C*H_ax_*H_eq_CH_2_CH), 1.63–1.72 (m, 1H, NCH_2_CH_ax_*H_eq_*CH_2_CH), 1.83–1.92 (m, 1H, NCH_2_CHCH_ax_*H_eq_*), 1.99 (td, *J*=10.9/3.0 Hz, 1H, NC*H_ax_*H_eq_CH_2_CH_2_CH), 2.14 (t, *J*=10.6 Hz, 1H, NC*H_ax_*H_eq_CHCH_2_), 2.29–2.39 (m, 2H, NCH_2_C*H_2_*CHCC), 2.39–2.54 (m, 3H, NCH_2_C*H_ax_*CH_2_ and NC*H_2_*CH_2_CHCC), 2.67–2.79 (m, 1H, NCH_ax_*H_eq_*CH_2_CH_2_CH), 2.94 (d_br_, *J*=11.6 Hz, 1H, NCH_ax_*H_eq_*CHCH_2_), 4.08 (q, *J*=7.1 Hz, 2H, OC*H_2_*CH_3_), 5.76 (t, *J*=6.7 Hz, 1H, NCH_2_CH_2_C*H*CC), 7.24–7.30 (m, 4H, ArH), 7.30–7.35 (m, 4H, ArH); ^13^C NMR (101 MHz, CD_2_Cl_2_): *δ*=14.6 (1C, OCH_2_*C*H_3_), 25.2 (1C, NCH_2_*C*H_2_CH_2_CH), 27.1 (1C, NCH_2_*C*H_2_CHCC), 27.6 (1C, NCH_2_CH_2_*C*H_2_CH), 42.5 (1C, NCH_2_CH_2_CH_2_*C*H), 53.5 (1C, N*C*H_2_CH_2_CH_2_CH), 56.1 (1C, N*C*H_2_CHCH_2_), 58.3 (1C, N*C*H_2_CH_2_CHCC), 60.7 (1C, O*C*H_2_CH_3_), 93.9 (1C, NCH_2_CH_2_*C*HCC), 108.6 (1C, NCH_2_CH_2_CHC*C*), 129.0 (4C, ArC), 130.2 (2C, ArC), 130.2 (2C, ArC), 133.3 (2C, ArC_q_), 136.0 (1C, ArC_q_), 136.0 (1C, ArC_q_), 174.5 (1C, *C*O), 206.1 (1C, CH_2_CH*C*C); IR (film): ṽ=2941, 2855, 2807, 2361, 2230, 1941, 1727, 1590, 1489, 1467, 1444, 1396, 1370, 1303, 1274, 1181, 1152, 1133, 1090, 1032, 1014, 949, 909, 887, 832, 795, 746, 721, 702 cm^-1^; HRMS-ESI *m/z* [*M*+H]^+^ calcd for C_25_H_27_Cl_2_NO_2_: 444.1497, found: 444.1495.

### *rac*-{Ethyl 1-[5,5-bis(4-methoxyphenyl)penta-3,4-dien-1-yl]piperidine-3- carboxylate} (*rac*-16e)

GP5 was followed using alkyne *rac*-**15** (105 mg, 0.500 mmol), CuI (19 mg, 0.10 mmol), *i*-Pr_2_NH (0.08 mL, 0.55 mmol) and 4,4’-(diazomethylene)bis(4- methoxybenzene) (520 mg, crude compound) in 1,4-dioxane (2.5 mL). After purification by column chromatography (PE/EtOAc 8:2) pure *rac*-**16e** was afforded as yellow viscous oil (97.7 mg, 45%): *R*_f_=0.24 (PE/EtOAc 7:3); ^1^H NMR (400 MHz, CDCl_3_): *δ*=1.24 (t, *J*=7.1 Hz, 3H, OCH_2_C*H*_3_), 1.41 (qd, *J*=12.0/4.0 Hz, 1H, NCH_2_CHC*H_ax_*H_eq_), 1.47–1.63 (m, 1H, NCH_2_C*H_ax_*H_eq_CH_2_CH), 1.64–1.76 (m, 1H, NCH_2_CH_ax_*H_eq_*CH_2_CH), 1.87–2.08 (m, 2H, NCH_2_CHCH_ax_*H_eq_* and NC*H_ax_*H_eq_CH_2_CH_2_CH), 2.17 (t, *J*=10.7 Hz, 1H, NC*H_ax_*H_eq_CHCH_2_), 2.36 (q, *J*=7.2 Hz, 2H, NCH_2_C*H_2_*CHCC), 2.45–2.66 (m, 3H, NCH_2_C*H_ax_*CH_2_ and NC*H_2_*CH_2_CHCC), 2.80 (d_br_, *J*=11.3 Hz, 1H, NCH_ax_*H_eq_*CH_2_CH_2_CH), 3.02 (d_br_, *J*=11.2 Hz, 1H, NCH_ax_*H_eq_*CHCH_2_), 3.82 (s, 6H, OC*H*_3_), 4.12 (q, *J*=7.1 Hz, 2H, OC*H_2_*CH_3_), 5.65 (t, *J*=6.6 Hz, 1H, NCH_2_CH_2_C*H*CC), 6.78–6.94 (m, 4H, ArH), 7.17–7.35 (m, 4H, ArH); ^13^C NMR (101 MHz, CDCl_3_): *δ*=14.2 (1C, OCH_2_*C*H_3_), 24.6 (1C, NCH_2_*C*H_2_CH_2_CH), 26.8 (1C, NCH_2_*C*H_2_CHCC), 27.0 (1C, NCH_2_CH_2_*C*H_2_CH), 41.9 (1C, NCH_2_CH_2_CH_2_*C*H), 53.7 (1C, N*C*H_2_CH_2_CH_2_CH), 55.3 (2C, O*C*H_3_), 55.4 (1C, N*C*H_2_CHCH_2_), 58.1 (1C, N*C*H_2_CH_2_CHCC), 60.3 (1C, O*C*H_2_CH_3_), 91.8 (1C, NCH_2_CH_2_*C*HCC), 109.0 (1C, NCH_2_CH_2_CHC*C*), 113.7 (4C, ArC), 129.5 (4C, ArC), 129.6 (2C, ArC_q_), 158.7 (2C, *C*OCH_3_), 174.2 (1C, *C*O), 204.9 (1C, NCH_2_CH_2_CH*C*C); IR (film): ṽ=2938, 2835, 2805, 1940, 1730, 1606, 1578, 1508, 1464, 1442, 1370, 1296, 1247, 1174, 1152, 1106, 1034, 963, 935, 910, 833, 807, 779, 730 cm^-1^; HRMS-ESI *m/z* [*M*+H]^+^ calcd for C_27_H_33_NO_4_: 436.2488, found: 436.2486.

### *rac*-{Ethyl (*R_a_*)-1-[5-(1,1’-biphenyl-4-yl)-5-phenylpenta-3,4-dien-1-yl]-(*3R*)- piperidine-3-carboxylate} [*rac*-(*3R*,*R_a_*)-16f] and *rac*-{ethyl (*S_a_*)-1-[5-(1,1’-biphenyl-4-yl)-5-phenylpenta-3,4-dien-1-yl]-(*3R*)-piperidine-3-carboxylate} [*rac*-(*3R*,*S_a_*)- 16f]

GP5 was followed using alkyne *rac*-**15** (105 mg, 0.500 mmol), CuI (19 mg, 0.10 mmol), *i*-Pr_2_NH (0.08 mL, 0.55 mmol) and 4-[diazo(phenyl)methyl]-1,1’-biphenyl (135 mg, 0.50 mmol) in 1,4-dioxane (2.5 mL) for 1 h. After purification by column chromatography (gradient elution PE/EtOAc 9:1 to PE/EtOAc 7:3) pure *rac*-(*3R*,*R_a_*)- **16f** and *rac*-(*3R*,*S_a_*)-**16f** were obtained as ∼ 1:1 mixture of racemic diastereomers as yellow viscous oil (198 mg, 88%): *R*_f_=0.14 (PE/EtOAc 8:2); ^1^H NMR (400 MHz, C_2_D_2_Cl_4_): *δ*=1.15 (t, *J*=7.1 Hz, 1.5H, OCH_2_C*H_3_*, dia1 or dia2), 1.16 (t, *J*=7.1 Hz, 1.5H, OCH_2_C*H*_3_, dia1 or dia2), 1.32 (qd, *J*=11.7/3.7 Hz, 1H, NCH_2_CHC*H_ax_*H_eq_), 1.38–1.52 (m, 1H, NCH_2_C*H_ax_*H_eq_CH_2_CH), 1.53–1.67 (m, 1H, NCH_2_CH_ax_*H_eq_*CH_2_CH), 1.76–1.88 (m, 1H, NCH_2_CHCH_ax_*H_eq_*), 1.93 (t_br_, *J*=10.8 Hz, 1H, NC*H_ax_*H_eq_CH_2_CH_2_CH), 2.09 (t, *J*=10.6 Hz, 1H, NC*H_ax_*H_eq_CHCH_2_), 2.33 (q, *J*=7.1 Hz, 2H, NCH_2_C*H_2_*CHCC), 2.38–2.56 (m, 3H, NCH_2_C*H_ax_*CH_2_ and NC*H_2_*CH_2_CHCC), 2.71 (d_br_, *J*=11.1 Hz, 1H, NCH_ax_*H_eq_*CH_2_CH_2_CH), 2.92 (d_br_, *J*=11.0 Hz, 1H, NCH_ax_*H_eq_*CHCH_2_), 4.01 (q, *J*=7.1 Hz, 1H, OC*H_2_*CH_3_, dia1 or dia2), 4.02 (q, *J*=7.1 Hz, 1H, OC*H_2_*CH_3_, dia1 or dia2), 5.69 (t, *J*=7.1 Hz, 1H, NCH_2_CH_2_C*H*CC), 7.18–7.45 (m, 10H, ArH), 7.45–7.63 (m, 4H, ArH); ^13^C NMR (101 MHz, C_2_D_2_Cl_4_): *δ*=14.6 (1C, OCH_2_*C*H_3_), 24.9 (1C, NCH_2_*C*H_2_CH_2_CH), 26.8 (1C, NCH_2_*C*H_2_CHCC or NCH_2_CH_2_*C*H_2_CH), 27.3 (1C, NCH_2_*C*H_2_CHCC or NCH_2_CH_2_*C*H_2_CH), 42.1 (1C, NCH_2_CH_2_CH_2_*C*H), 53.8 (1C, N*C*H_2_CH_2_CH_2_CH), 55.6 (1C, N*C*H_2_CHCH_2_), 58.2 (1C, N*C*H_2_CH_2_CHCC), 60.7 (1C, O*C*H_2_CH_3_), 92.9 (1C, NCH_2_CH_2_*C*HCC), 109.8 (1C, NCH_2_CH_2_CHC*C*), 127.3 (2C, ArC), 127.3 (2C, ArC), 127.5 (1C, ArC), 127.7 (1C, ArC), 128.7 (2C, ArC), 128.9 (1C, ArC), 128.9 (1C, ArC), 129.1 (1C, ArC), 129.1 (1C, ArC), 129.2 (2C, ArC), 136.3 (1C, ArC_q_), 137.1 (1C, ArC_q_), 139.8 (1C, ArC_q_), 140.8 (1C, ArC_q_), 174.6 (1C, *C*O), 206.0 (1C, NCH_2_CH_2_CH*C*C); IR (film): ṽ=3057, 3028, 2940, 2854, 2804, 1940, 1730, 1598, 1519, 1486, 1468, 1446, 1370, 1310, 1273, 1210, 1178, 1151, 1133, 1100, 1074, 1030, 1007, 964, 906, 842, 799, 767, 728, 697 cm^-1^; HRMS-ESI *m/z* [*M*+H]^+^ calcd for C_31_H_33_NO_2_: 452.2590, found: 452.2578.

### *rac*-[1-(5,5-Diphenylpenta-3,4-dien-1-yl)piperidine-3-carboxylic acid] (*rac*-11a)

GP7 was followed using nipecotic acid ester *rac*-**16a** (93.9 mg, 0.250 mmol) and Ba(OH)_2_x8H_2_O (158 mg, 0.500 mmol) in 1.8 mL EtOH/H_2_O (2:1) for 25 h. After purification by RP-MPLC (MeOH/H_2_O 7:3) amino acid *rac*-**11a** was afforded as white amorphous solid (45.3 mg, 52%): mp: 41 °C; ^1^H NMR (500 MHz, MeOD, NaOD): *δ*=1.13 (qd, *J*=13.5/4.4 Hz, 1H, NCH_2_CHC*H_ax_*H_eq_), 1.36 (qt, *J*=12.9/3.8 Hz, 1H, NCH_2_C*H_ax_*H_eq_CH_2_CH), 1.43–1.55 (m, 1H, NCH_2_CH_ax_*H_eq_*CH_2_CH), 1.67–1.82 (m, 2H, NC*H_ax_*H_eq_CH_2_CH_2_CH and NCH_2_CHCH_ax_*H_eq_*), 1.86 (t, *J*=11.4 Hz, 1H, NC*H_ax_*H_eq_CHCH_2_), 2.11–2.30 (m, 3H, NCH_2_C*H_ax_*CH_2_ and NCH_2_C*H_2_*CHCC), 2.32–2.43 (m, 2H, NC*H_2_*CH_2_CHCC), 2.71 (d_br_, *J*=11.4 Hz, 1H, NCH_ax_*H_eq_*CH_2_CH_2_CH), 2.95 (d_br_, *J*=11.0 Hz, 1H, NCH_ax_*H_eq_*CHCH_2_), 5.55 (t, *J*=6.7 Hz, 1H, NCH_2_CH_2_C*H*CC), 7.00–7.19 (m, 10H, ArH); ^13^C NMR (101 MHz, MeOD, NaOD): *δ*=26.0 (1C, NCH_2_*C*H_2_CH_2_CH), 27.2 (1C, NCH_2_*C*H_2_CHCC), 29.5 (1C, NCH_2_CH_2_*C*H_2_CH), 46.3 (1C, NCH_2_CH_2_CH_2_*C*H), 54.9 (1C, N*C*H_2_CH_2_CH_2_CH), 58.2 (1C, N*C*H_2_CHCH_2_), 59.4 (1C, N*C*H_2_CH_2_CHCC), 92.9 (1C, NCH_2_CH_2_*C*HCC), 111.5 (1C, NCH_2_CH_2_CHC*C*), 128.2 (2C, ArC), 129.4 (4C, ArC), 129.4 (4C, ArC), 138.3 (2C, ArC_q_), 182.7 (1C, *C*O), 206.7 (1C, CH_2_CH*C*C); IR (KBr): ṽ=3433, 3056, 3026, 2938, 2857, 2805, 2525, 2368, 2345, 1944, 1708, 1597, 1491, 1451, 1442, 1389, 1310, 1278, 1218, 1188, 1155, 1101, 1074, 1030, 921, 902, 770, 696, 630 cm^-1^; HRMS-ESI *m/z* [*M*+H]^+^ calcd for C_23_H_25_NO_2_: 348.1964, found: 348.1958; average purity by qHNMR (internal calibrant: maleic acid Lot#BCBM8127V): 99%.

### *rac*-[1-(5,5-Di-*p*-tolylpenta-3,4-dien-1-yl)piperidine-3-carboxylic acid] (*rac*-11b)

GP7 was followed using nipecotic acid ester *rac*-**16b** (109 mg, 0.269 mmol) and Ba(OH)_2_x8H_2_O (170 mg, 0.538 mmol) in 1.5 mL EtOH/H_2_O (2:1) for 5 h. After purification by RP-MPLC (MeOH/H_2_O 7:3) amino acid *rac*-**11b** was afforded as white amorphous solid (14.6 mg, 15%): mp: 54 °C; ^1^H NMR (500 MHz, MeOD): *δ*=1.33 (qd, *J*=13.0/4.4 Hz, 1H, NCH_2_CHC*H_ax_*H_eq_), 1.51–1.63 (m, 1H, NCH_2_C*H_ax_*H_eq_CH_2_CH), 1.64–1.74 (m, 1H, NCH_2_CH_ax_*H_eq_*CH_2_CH), 1.93 (td, *J*=11.8/2.9 Hz, 1H, NC*H_ax_*H_eq_CH_2_CH_2_CH), 1.97–2.03 (m, 1H, NCH_2_CHCH_ax_*H_eq_*), 2.05 (t, *J*=11.4 Hz, 1H, NC*H_ax_*H_eq_CHCH_2_), 2.33 (s, 6H, C*H_3_*), 2.35–2.45 (m, 3H, NCH_2_C*H_2_*CHCC and NCH_2_C*H_ax_*CH_2_), 2.52–2.61 (m, 2H, NC*H_2_*CH_2_CHCC), 2.90 (d_br_, *J*=11.3 Hz, 1H, NCH_ax_*H_eq_*CH_2_CH_2_CH), 3.16 (d_br_, *J*=11.5 Hz, 1H, NCH_ax_*H_eq_*CHCH_2_), 5.69 (t, *J*=6.6 Hz, 1H, NCH_2_CH_2_C*H*CC), 7.11–7.23 (m, 8H, ArH); ^13^C NMR (126 MHz, MeOD): *δ*=21.2 (2C, *C*H_3_), 26.0 (1C, NCH_2_*C*H_2_CH_2_CH), 27.3 (1C, NCH_2_*C*H_2_CHCC), 29.5 (1C, NCH_2_CH_2_*C*H_2_CH), 46.3 (1C, NCH_2_CH_2_CH_2_*C*H), 54.9 (1C, N*C*H_2_CH_2_CH_2_CH), 58.2 (1C, N*C*H_2_CHCH_2_), 59.5 (1C, N*C*H_2_CH_2_CHCC), 92.5 (1C, NCH_2_CH_2_*C*HCC), 111.2 (1C, NCH_2_CH_2_CHC*C*), 129.3 (4C, ArC), 130.0 (4C, ArC), 135.5 (2C, ArC_q_), 138.0 (2C, ArC_q_), 182.7 (1C, *C*O), 206.4 (1C, NCH_2_CH_2_CH*C*C); IR (KBr): ṽ=3432, 3021, 2922, 2858, 1942, 1702, 1593, 1509, 1449, 1386, 1308, 1181, 1153, 1108, 1020, 910, 822, 796, 783, 720, 668, 611, 584, 514, 462 cm^-1^; HRMS-ESI *m/z* [*M*+H]^+^ calcd for C_25_H_29_NO_2_: 376.2277, found: 376.2266.

### *rac*-{1-[5,5-Bis(4-fluorophenyl)penta-3,4-dien-1-yl]piperidine-3-carboxylic acid} (*rac*-11c)

GP7 was followed using nipecotic acid ester *rac*-**16c** (194 mg, 0.472 mmol) and Ba(OH)_2_x8H_2_O (298 mg, 0.944 mmol) in 3.0 mL EtOH/H_2_O (2:1) overnight. After purification by RP-MPLC (gradient elution with MeOH/H_2_O 6:4 to MeOH/H_2_O 7:3) amino acid *rac*-**11c** was afforded as white amorphous solid (82.4 mg, 46%): mp: 62°C; ^1^H NMR (400 MHz, MeOD, NaOD): *δ*=1.33 (qd, *J*=12.8/4.2 Hz, 1H, NCH_2_CHC*H_ax_*H_eq_), 1.55 (qt, *J*=12.8/3.9 Hz, 1H, NCH_2_C*H_ax_*H_eq_CH_2_CH), 1.63–1.76 (m, 1H, NCH_2_CH_ax_*H_eq_*CH_2_CH), 1.86–2.01 (m, 2H, NC*H_ax_*H_eq_CH_2_CH_2_CH and NCH_2_CHCH_ax_*H_eq_*), 2.05 (t, *J*=11.3 Hz, 1H, NC*H_ax_*H_eq_CHCH_2_), 2.32–2.47 (m, 3H, NCH_2_C*H_ax_*CH_2_ and NCH_2_C*H_2_*CHCC), 2.50–2.61 (m, 2H, NC*H_2_*CH_2_CHCC), 2.90 (d_br_, *J*=11.3 Hz, 1H, NCH_ax_*H_eq_*CH_2_CH_2_CH), 3.08–3.19 (m, 1H, NCH_ax_*H_eq_*CHCH_2_), 5.77 (t, *J*=6.6 Hz, 1H, NCH_2_CH_2_C*H*CC), 7.01–7.14 (m, 4H, FCC*H*), 7.22–7.37 (m, 4H, FCCHC*H*); ^13^C NMR (101 MHz, MeOD, NaOD): *δ*=26.0 (1C, NCH_2_*C*H_2_CH_2_CH), 27.2 (1C, NCH_2_*C*H_2_CHCC), 29.5 (1C, NCH_2_CH_2_*C*H_2_CH), 46.3 (1C, NCH_2_CH_2_CH_2_*C*H), 54.9 (1C, N*C*H_2_CH_2_CH_2_CH), 58.2 (1C, N*C*H_2_CHCH_2_), 59.3 (1C, N*C*H_2_CH_2_CHCC), 93.5 (1C, NCH_2_CH_2_*C*HCC), 109.7 (1C, NCH_2_CH_2_CHC*C*), 116.3 (d*_CF_*, *^2^J_CF_*=21.8 Hz, 2C, FC*C*HCHC), 116.3 (d*_CF_*, *^2^J_CF_*=21.8 Hz, 2C, FC*C*HCHC), 131.1 (d*_CF_*, *^3^J_CF_*=8.0 Hz, 4C, FCCH*C*HC), 134.4 (d*_CF_*, *^4^J_CF_*=3.4 Hz, 2C, FCCHCH*C*), 163.5 (d*_CF_*, *^1^J_CF_*=245.3 Hz, 2C, F*C*), 182.8 (1C, *C*O), 206.5 (1C, CH_2_CH*C*C); ^19^F NMR (376 MHz, MeOD, NaOD): *δ*=- 116.5, -116.6; IR (KBr): ṽ=3422, 3044, 2942, 2861, 2804, 2494, 2027, 1944, 1896, 1711, 1600, 1505, 1468, 1450, 1390, 1298, 1281, 1222, 1156, 1096, 1013, 911, 838, 799, 767, 724, 606, 582, 550, 519, 490 cm^-1^; HRMS-ESI *m/z* [*M*+H]^+^ calcd for C_23_H_23_F_2_NO_2_: 384.1775, found: 384.1766.

### *rac*-{1-[5,5-Bis(4-chlorophenyl)penta-3,4-dien-1-yl]piperidine-3-carboxylic acid} (*rac*-11d)

GP6 was followed using nipecotic acid ester *rac*-**16d** (0.510 mmol, 227 mg), EtOH (2.55 mL) and 2M NaOH (0.77 mL) for 45 min. The amino acid *rac*-**11d** was obtained as colorless amorphous solid (53.2 mg, 25%): mp: 79 °C; ^1^H NMR (500 MHz, MeOD, NaOD): *δ*=1.01 (qd, *J*=12.7/3.9 Hz, 1H, NCH_2_CHC*H_ax_*H_eq_), 1.15–1.30 (m, 1H, NCH_2_C*H_ax_*H_eq_CH_2_CH), 1.30–1.42 (m, 1H, NCH_2_CH_ax_*H_eq_*CH_2_CH), 1.55–1.69 (m, 2H, NCH_2_CHCH_ax_*H_eq_* and NC*H_ax_*H_eq_CH_2_CH_2_CH), 1.72 (t, *J*=11.3 Hz, 1H, NC*H_ax_*H_eq_CHCH_2_), 1.99–2.16 (m, 3H, NCH_2_C*H_2_*CHCC and NCH_2_C*H_ax_*CH_2_), 2.23 (t, *J*=7.7 Hz, 2H, NC*H_2_*CH_2_CHCC), 2.57 (d_br_, *J*=11.3 Hz, 1H, NCH_ax_*H_eq_*CH_2_CH_2_CH), 2.82 (d_br_, *J*=11.2 Hz, 1H, NCH_ax_*H_eq_*CHCH_2_), 5.49 (t, *J*=6.6 Hz, 1H, NCH_2_CH_2_C*H*CC), 6.95 (d, *J*=8.1 Hz, 4H, ArH), 7.02 (d, *J*=8.1 Hz, 4H, ArH); ^13^C NMR (126 MHz, MeOD, NaOD): *δ*=26.0 (1C, NCH_2_*C*H_2_CH_2_CH), 27.0 (1C, NCH_2_*C*H_2_CHCC), 29.5 (1C, NCH_2_CH_2_*C*H_2_CH), 46.3 (1C, NCH_2_CH_2_CH_2_*C*H), 54.9 (1C, N*C*H_2_CH_2_CH_2_CH), 58.2 (1C, N*C*H_2_CHCH_2_), 59.3 (1C, N*C*H_2_CH_2_CHCC), 94.0 (1C, NCH_2_CH_2_*C*HCC), 109.6 (1C, NCH_2_CH_2_CHC*C*), 129.7 (2C, ArC), 129.7 (2C, ArC), 130.8 (4C, ArC), 134.2 (2C, ArC_q_), 136.7 (2C, ArC_q_), 182.8 (1C, *C*O), 206.8 (1C, CH_2_CH*C*C); IR (KBr): ṽ=3435, 2936, 2855, 2803, 1941, 1711, 1590, 1488, 1450, 1396, 1299, 1218, 1154, 1090, 1013, 909, 832, 745, 722, 701, 639, 613, 595, 532, 511, 461 cm^-1^; HRMS-ESI *m/z* [*M*+H]^+^ calcd for C_23_H_23_Cl_2_NO_2_: 416.1184, found: 416.1178; average purity by qHNMR (internal calibrant: maleic acid Lot#BCBM8127V): 98%.

### *rac*-{(*R_a_*)-1-[5-(1,1’-biphenyl-4-yl)-5-phenylpenta-3,4-dien-1-yl]-(*3R*)-piperidine-3-carboxylic acid} [*rac*-(*3R*,*R_a_*)-11f] and *rac*-{(*S_a_*)-1-[5-(1,1’-biphenyl-4-yl)-5- phenylpenta-3,4-dien-1-yl]-(*3R*)-piperidine-3-carboxylic acid} [*rac*-(*3R*,*S_a_*)-11f]

GP6 was followed using nipecotic acid esters *rac*-(*3R*,*R_a_*)-**16f** and *rac*-(*3R*,*S_a_*)-**16f** (0.202 mmol, 91.2 mg), EtOH (1.0 mL) and 2M NaOH (0.30 mL) for 2.5 h. The amino acids *rac*-(*3R*,*R_a_*)-**11f** and *rac*-(*3R*,*S_a_*)-**11f** were obtained as ∼ 1:1 mixture of racemic diastereomers as yellow amorphous solid (25.6 mg, 30%): mp: 86 °C; ^1^H NMR (500 MHz, MeOD, NaOD): *δ*=1.34 (qd, *J*=12.8/4.2 Hz, 1H, NCH_2_CHC*H_ax_*H_eq_), 1.57 (qt, *J*=13.0/4.0 Hz, 1H, NCH_2_C*H_ax_*H_eq_CH_2_CH), 1.65–1.73 (m, 1H, NCH_2_CH_ax_*H_eq_*CH_2_CH), 1.91–2.02 (m, 2H, NCH_2_CHCH_ax_*H_eq_* and NC*H_ax_*H_eq_CH_2_CH_2_CH), 2.07 (t, *J*=11.4 Hz, 1H, NC*H_ax_*H_eq_CHCH_2_), 2.33–2.51 (m, 3H, NCH_2_C*H_2_*CHCC and NCH_2_C*H_ax_*CH_2_), 2.54–2.64 (m, 2H, NC*H_2_*CH_2_CHCC), 2.92 (d_br_, *J*=11.3 Hz, 1H, NCH_ax_*H_eq_*CH_2_CH_2_CH), 3.16 (dd_br_, *J*=10.5/3.5 Hz, 1H, NCH_ax_*H_eq_*CHCH_2_), 5.79 (t, *J*=6.6 Hz, 1H, NCH_2_CH_2_C*H*CC), 7.24–7.47 (m, 10H, ArH), 7.57–7.66 (m, 4H, ArH); ^13^C NMR (126 MHz, MeOD, NaOD): *δ*=26.0 (1C, NCH_2_*C*H_2_CH_2_CH), 27.2 (1C, NCH_2_*C*H_2_CHCC), 29.5 (1C, NCH_2_CH_2_*C*H_2_CH), 46.3 (1C, NCH_2_CH_2_CH_2_*C*H), 54.9 (1C, N*C*H_2_CH_2_CH_2_CH), 58.3 (1C, N*C*H_2_CHCH_2_), 59.5 (1C, N*C*H_2_CH_2_CHCC), 93.1 (1C, NCH_2_CH_2_*C*HCC), 111.2 (1C, NCH_2_CH_2_CHC*C*), 127.9 (2C, ArC), 128.0 (2C, ArC), 128.3 (1C, ArC), 128.4 (1C, ArC), 129.5 (2C, ArC), 129.5 (2C, ArC), 129.8 (2C, ArC), 129.9 (2C, ArC), 137.3 (1C, ArC_q_), 138.3 (1C, ArC_q_), 141.3 (1C, ArC_q_), 142.0 (1C, ArC_q_), 182.8 (1C, *C*O), 206.9 (1C, NCH_2_CH_2_CH*C*C); IR (KBr): ṽ=3422, 3054, 3027, 2931, 2855, 2802, 1943, 1711, 1655, 1598, 1485, 1446, 1390, 1312, 1279, 1183, 1154, 1101, 1074, 1029, 1007, 907, 842, 768, 729, 697, 629 cm^-1^; HRMS-ESI *m/z* [*M*+H]^+^ calcd for C_29_H_29_NO_2_: 424.2277, found: 424.2269.

### *rac*-[Ethyl (*R_a_*)-1-{3-[6-methoxy-3,4-dihydronaphthalen-1(*2H*)-ylidene]allyl}-(*3R*)- piperidine-3-carboxylate] [*rac*-(*3R*,*R_a_*)-14g] and *rac*-[ethyl (*S_a_*)-1-{3-[6-methoxy-3,4-dihydronaphthalen-1(*2H*)-ylidene]allyl}-(*3R*)-piperidine-3-carboxylate] [*rac*- (*3R*,*S_a_*)-14g]

GP4 was followed applying alkyne *rac*-**12**^26^^26^ (98 mg, 0.50 mmol), CuI (19 mg, 0.10 mmol), Li*t*OBu (140 mg, 1.75 mmol), and *N’*-[6-methoxy-3,4-dihydronaphthalen-1(*2H*)-ylidene]-4-methylbenzenesulfonohydrazide (**18**, 379 mg, 1.10 mmol) in 1,4-dioxane (5.0 mL). The solution was stirred at 90 °C for 1 h. After purification by column chromatography (PE/EtOAc 7:3) *rac*-(*3R*,*R_a_*)-**14g** and *rac*- (*3R*,*S_a_*)-**14g** were obtained as ∼ 1:1 mixture of racemic diastereomers as yellow oil (107 mg, 60%): *R*_f_=0.35 (PE/EtOAc 6:4); ^1^H NMR (500 MHz, CD_2_Cl_2_): *δ*=1.23 (t, *J*=7.1 Hz, 1.5H, OCH_2_C*H*_3_, dia2), 1.24 (t, *J*=7.1 Hz, 1.5H, OCH_2_C*H*_3_, dia1), 1.43 (qd, *J*=11.8/4.0 Hz, 1H, NCH_2_CHC*H_ax_*H_eq_), 1.50–1.66 (m, 1H, NCH_2_C*H_ax_*H_eq_CH_2_CH), 1.68–1.77 (m, 1H, NCH_2_CH_ax_*H_eq_*CH_2_CH), 1.86 (p, *J*=6.2 Hz, 2H, NCH_2_CHCCCH_2_C*H_2_*CH_2_), 1.89–1.95 (m, 1H, NCH_2_CHCH_ax_*H_eq_*), 2.10 (td, *J*=11.0/2.4 Hz, 1H, NC*H_ax_*H_eq_CH_2_CH_2_CH), 2.23 (t, *J*=10.6 Hz, 1H, NC*H_ax_*H_eq_CHCH_2_), 2.46–2.59 (m, 3H, NCH_2_C*H_ax_*CH_2_ and NCH_2_CHCCC*H_2_*CH_2_CH_2_), 2.77 (t, *J*=6.2 Hz, 2H, NCH_2_CHCCCH_2_CH_2_C*H_2_*), 2.78–2.87 (m, 1H, NCH_ax_*H_eq_*CH_2_CH_2_CH), 2.98–3.08 (m, 1H, NCH_ax_*H_eq_*CHCH_2_), 3.09 (d, *J*=6.9 Hz, 1H, NC*H_2_*CHCC, dia2), 3.10 (d, *J*=6.9 Hz, 1H, NC*H_2_*CHCC, dia1), 3.76 (s, 3H, OC*H_3_*), 4.10 (q, *J*=7.1 Hz, 1H, OC*H*_2_CH_3_, dia2), 4.10 (q, *J*=7.1 Hz, 1H, OC*H*_2_CH_3_, dia1), 5.45 (tt, *J*=6.9/3.0 Hz, 1H, NCH_2_C*H*CC), 6.62 (d, *J*=2.8 Hz, 1H, CCC*H*COCH_3_), 6.68 (dt, *J*=8.6/2.8 Hz, 1H, CH_3_OCC*H*CH), 7.33 (d, *J*=8.6 Hz, 1H, CH_3_OCCHC*H*); ^13^C NMR (126 MHz, CD_2_Cl_2_): *δ*=14.6 (0.5C, OCH_2_*C*H_3_, dia2), 14.6 (0.5C, OCH_2_*C*H_3_, dia1), 23.6 (1C, NCH_2_CHCCCH_2_*C*H_2_CH_2_), 25.2 (0.5C, NCH_2_CHCH_2_*C*H_2_CH_2_, dia1), 25.3 (0.5C, NCH_2_CHCH_2_*C*H_2_CH_2_, dia2), 27.4 (1C, NCH_2_CH*C*H_2_CH_2_CH_2_), 29.7 (1C, NCH_2_CHCC*C*H_2_CH_2_CH_2_), 30.9 (1C, NCH_2_CHCCCH_2_CH_2_*C*H_2_), 42.6 (0.5C, NCH_2_*C*HCH_2_CH_2_CH_2_, dia1 or dia2), 42.6 (0.5C, NCH_2_*C*HCH_2_CH_2_CH_2_, dia1 or dia2), 53.7 (0.5C, NCH_2_CHCH_2_CH_2_*C*H_2_, dia1 or dia2), 53.8 (0.5C, NCH_2_CHCH_2_CH_2_*C*H_2_, dia1 or dia2), 55.6 (0.5C, N*C*H_2_CHCH_2_CH_2_CH_2_, dia1 or dia2), 55.7 (0.5C, N*C*H_2_CHCH_2_CH_2_CH_2_, dia1 or dia2), 55.7 (1C, O*C*H_3_), 58.9 (0.5C, N*C*H_2_CHCC, dia1), 58.9 (0.5C, N*C*H_2_CHCC, dia2), 60.7 (1C, O*C*H_2_CH_3_), 91.2 (0.5C, NCH_2_*C*HCC, dia1), 91.2 (0.5C, NCH_2_*C*HCC, dia2), 101.9 (0.5C, NCH_2_CHC*C*, dia2), 101.9 (0.5C, NCH_2_CHC*C*, dia1), 113.2 (1C, CH_3_OC*C*HCH), 113.8 (1C, CC*C*HCOCH_3_), 124.5 (0.5C, ArC_q_, dia1 or dia2), 124.5 (0.5C, ArC_q_, dia1 or dia2), 128.6 (1C, CH_3_OCCH*C*H), 138.5 (1C, ArC_q_), 158.9 (1C, *C*OCH_3_), 174.6 (0.5C, *C*O, dia1 or dia2), 174.6 (0.5C, *C*O, dia1 or dia2), 202.6 (0.5C, NCH_2_CH*C*C, dia1 or dia2), 202.6 (0.5C, NCH_2_CH*C*C, dia1 or dia2); IR (film): ṽ=2937, 2862, 2834, 2796, 1946, 1730, 1607, 1571, 1498, 1466, 1441, 1366, 1319, 1261, 1245, 1180, 1152, 1132, 1111, 1094, 1068, 1033, 884, 836, 809 cm^-1^; HRMS-ESI *m/z* [*M*+H]^+^ calcd for C_22_H_29_NO_3_: 356.2226, found: 356.2221.

### *rac*-[(*R_a_*)-1-{3-[6-methoxy-3,4-dihydronaphthalen-1(*2H*)-ylidene]allyl}-(*3R*)- piperidine-3-carboxylic acid] [*rac*-(*3R*,*R_a_*)-8g] and *rac*-[(*S_a_*)-1-{3-[6-methoxy-3,4- dihydronaphthalen-1(*2H*)-ylidene]allyl}-(*3R*)-piperidine-3-carboxylic acid] [*rac*- (*3R*,*S_a_*)-8g]

GP7 was followed using nipecotic acid esters *rac*-(*3R*,*R_a_*)-**14g** and *rac*- (*3R*,*S_a_*)-**14g** (107 mg, 0.300 mmol) and Ba(OH)_2_x8H_2_O (189 mg, 0.600 mmol) in 2.1 mL EtOH/H_2_O (2:1) for 24 h. After purification by RP-MPLC (MeOH/H_2_O 6:4) the amino acids *rac*-(*3R*,*R_a_*)-**8g** and *rac*-(*3R*,*S_a_*)**-8g** were obtained as ∼ 1:1 mixture of racemic diastereomers as pale yellow amorphous solid (59.6 mg, 61%): mp.: 115.2 °C; ^1^H NMR (400 MHz, MeOD): *δ*=1.37 (qd, *J*=12.7/4.0 Hz, 1H, NCH_2_CHC*H_ax_*H_eq_), 1.54–1.69 (m, 1H, NCH_2_C*H_ax_*H_eq_CH_2_CH), 1.70–1.80 (m, 1H, NCH_2_CH_ax_*H_eq_*CH_2_CH), 1.85 (p, *J*=6.1 Hz, 2H, NCH_2_CHCCCH_2_C*H_2_*CH_2_), 1.95–2.07 (m, 2H, NCH_2_CHCH_ax_*H_eq_* and NC*H_ax_*H_eq_CH_2_CH_2_CH), 2.13 (t, *J*=11.4 Hz, 0.5H, NC*H_ax_*H_eq_CHCH_2_, dia1 or dia2), 2.16 (t, *J*=11.4 Hz, 0.5H, NC*H_ax_*H_eq_CHCH_2_, dia1 or dia2), 2.35–2.48 (m, 1H, NCH_2_C*H_ax_*CH_2_), 2.48–2.61 (m, 2H, NCH_2_CHCCC*H_2_*CH_2_CH_2_), 2.76 (t, *J*=6.1 Hz, 2H, NCH_2_CHCCCH_2_CH_2_C*H_2_*), 2.92–3.05 (m, 1H, NCH_ax_*H_eq_*CH_2_CH_2_CH), 3.05–3.18 (m, 2H, NC*H_2_*CHCC), 3.18–3.28 (m, 1H, NCH_ax_*H_eq_*CHCH_2_), 3.75 (s, 3H, OC*H_3_*), 5.40–5.56 (m, 1H, NCH_2_C*H*CC), 6.63 (d, *J*=2.6 Hz, 0.5H, CCC*H*COCH_3_, dia1 or dia2), 6.63 (d, *J*=2.4 Hz, 0.5H, CCC*H*COCH_3_, dia1 or dia2), 6.69 (dd, *J*=8.6/2.6 Hz, 0.5H, CH_3_OCC*H*CH, dia1 or dia2), 6.71 (dd, *J*=8.6/2.6 Hz, 0.5H, CH_3_OCC*H*CH, dia1 or dia2), 7.29 (d, *J*=8.6 Hz, 0.5H, CH_3_OCCHC*H*, dia1 or dia2), 7.32 (d, *J*=8.6 Hz, 0.5H, CH_3_OCCHC*H*, dia1 or dia2); ^13^C NMR (101 MHz, MeOD): *δ*=24.2 (1C, NCH_2_CHCCCH_2_*C*H_2_CH_2_), 26.0 (0.5C, NCH_2_CHCH_2_*C*H_2_CH_2_, dia1 or dia2), 26.0 (0.5C, NCH_2_CHCH_2_*C*H_2_CH_2_, dia1 or dia2), 29.4 (1C, NCH_2_CH*C*H_2_CH_2_CH_2_), 30.2 (1C, NCH_2_CHCC*C*H_2_CH_2_CH_2_), 31.4 (1C, NCH_2_CHCCCH_2_CH_2_*C*H_2_), 46.5 (0.5C, NCH_2_*C*HCH_2_CH_2_CH_2_, dia1 or dia2), 46.6 (0.5C, NCH_2_*C*HCH_2_CH_2_CH_2_, dia1 or dia2), 54.3 (0.5C, NCH_2_CHCH_2_CH_2_*C*H_2_, dia1 or dia2), 54.5 (0.5C, NCH_2_CHCH_2_CH_2_*C*H_2_, dia1 or dia2), 55.6 (1C, O*C*H_3_), 57.8 (0.5C, N*C*H_2_CHCH_2_CH_2_CH_2_, dia1 or dia2), 57.9 (0.5C, N*C*H_2_CHCH_2_CH_2_CH_2_, dia1 or dia2), 59.6 (0.5C, N*C*H_2_CHCC, dia1 or dia2), 59.7 (0.5C, N*C*H_2_CHCC, dia1 or dia2), 90.6 (0.5C, NCH_2_*C*HCC, dia1 or dia2), 90.6 (0.5C, NCH_2_*C*HCC, dia1 or dia2), 103.0 (1C, NCH_2_CHC*C*), 113.8 (0.5C, CH_3_OC*C*HCH, dia1 or dia2), 113.8 (0.5C, CH_3_OC*C*HCH, dia1 or dia2), 114.4 (0.5C, CC*C*HCOCH_3_, dia1 or dia2), 114.4 (0.5C, CC*C*HCOCH_3_, dia1 or dia2), 124.7 (0.5C, ArC_q_, dia1 or dia2), 124.7 (0.5C, ArC_q_, dia1 or dia2), 129.2 (0.5C, CH_3_OCCH*C*H, dia1 or dia2), 129.3 (0.5C, CH_3_OCCH*C*H, dia1 or dia2), 139.0 (0.5C, ArC_q_, dia1 or dia2), 139.1 (0.5C, ArC_q_, dia1 or dia2), 160.0 (1C, *C*OCH_3_), 182.7 (1C, *C*O), 203.8 (0.5C, NCH_2_CH*C*C, dia1 or dia2), 203.8 (0.5C, NCH_2_CH*C*C, dia1 or dia2); IR (KBr): ṽ=3433, 2935, 2860, 2835, 2525, 1946, 1715, 1606, 1571, 1498, 1465, 1452, 1394, 1320, 1271, 1246, 1195, 1154, 1125, 1067, 1035, 953, 884, 837, 820, 766, 718, 668, 565 cm^-1^; HRMS-ESI *m/z* [*M*+H]^+^ calcd for C_20_H_25_NO_3_: 328.1913, found: 328.1907.

## Biological evaluations

### *In vitro* part

#### MS Binding Assays

MS Binding Assays were performed with mGAT1 membrane preparations obtained from a stable HEK293 cell line and NO711 as unlabeled marker in competitive binding experiments as described previously.^32^

#### [^3^H]GABA uptake assays

The [^3^H]GABA uptake assays were performed in a 96-well plate format with intact HEK293 cells stably expressing mGAT1, mGAT2, mGAT3, mGAT4 as described earlier.^33^

### *In vivo* part

#### Materials and methods

##### Animals and housing conditions

For *in vivo* tests that assessed the pharmacological activity of the compound (*S*)-**8d** (DDPM-3960) adult male Albino Swiss (CD-1) mice weighing 18 - 22 g were used. The animals were housed in groups of 10 mice per cage at room temperature of 22±2 °C, under light/dark (12:12) cycle. The animals had free access to food and tap water before the experiments. The ambient temperature of the experimental room and humidity (50 ± 10%) were kept consistent throughout all the tests. For behavioral experiments the animals were selected randomly. Each experimental group consisted of 6-12 animals/dose, and each mouse was used only once. The experiments were performed between 9 AM and 2 PM. Immediately after the *in vivo* assay the animals were euthanized by cervical dislocation. All procedures were approved by the Local Ethics Committee of the Jagiellonian University in Krakow (337/2019, release date: 30.10.2019) and the treatment of animals was in full accordance with ethical standards laid down in respective Polish and EU regulations (Directive No. 86/609/EEC).

#### Chemicals used in pharmacological tests

For *in vivo* tests the compound DDPM-3960 [(*S*)-**8d**] was prepared in 1% Tween 80 solution (Polskie Odczynniki Chemiczne, Poland) and was administered by the intraperitoneal route 60 min before behavioral tests. Control mice received 1% Tween 80. PTZ, pilocarpine hydrochloride, scopolamine butylbromide and capsaicin were provided by Sigma Aldrich (Poland). For the tests they were prepared in 0.9% saline (Polfa Kutno, Poland). Formalin (5% solution in distilled water, Polskie Odczynniki Chemiczne, Poland), PTZ and pilocarpine were administered 60 min after the test compound or vehicle.

To assess anticonvulsant properties of the test compound five assays were used: PTZ, maximal electroshock seizure (MES), electroconvulsive threshold, 6-Hz and pilocarpine tests. In these assays the dose of 60 mg/kg was chosen as a starting dose. If it turned out to be effective, lower doses were also tested.

The dose of 30 mg/kg was the highest dose tested in assays assessing potential anxiolytic-like, antidepressant-like and antinociceptive properties of DDPM-3960.

#### PTZ seizure test

The test was performed according to a method previously described.^45^ Clonic convulsions were induced by the subcutaneous administration of PTZ at a dose of 100 mg/kg. After PTZ injection, each mouse was immediately placed in a transparent Plexiglas cage (30 cm x 20 cm x 15 cm) and was observed during the next 30 min for the occurrence of clonic seizures. Clonic seizures were defined as clonus of the whole body lasting more than 3 s, with an accompanying loss of righting reflex. Latency time to first clonus, the number of seizure episodes and mortality rate were noted and compared between vehicle-treated and DDPM-3960-treated groups.

### Maximal electroshock seizure test

MES test was performed according to a method previously described.^45^ In this test vehicle-treated mice and drug-treated mice received a stimulus of 25 mA delivered by an electroshock generator (Hugo Sachs rodent shocker, Germany) to induce maximal seizures (tonic extension) of hind limbs. Electroconvulsions were produced with the use of auricular electrodes and the stimulus duration was 0.2 s. Tonic extension of the hind limbs was regarded as the endpoint for this procedure.

#### Electroconvulsive threshold test

The electroconvulsive threshold test was performed using the method recently described.^45^ In this assay, the estimated parameter was the CS_50_ value, defined as median current strength, i.e. current intensity that caused tonic hind limb extension in 50% of the mice challenged. To evaluate this parameter, at least four groups of animals were used. The mice were subjected to electroshocks of various intensities to yield 10- 30%, 30-50%, 50-70%, and 70-90% of animals with convulsions. CS_50_ value was determined using the log-probit method.^64^

This test was performed according to a method described by Barton and colleagues.^65^ It is an alternative electroshock paradigm that involves low-frequency (6 Hz), long-duration (3 s) electrical stimulation. Corneal stimulation (0.2 ms-duration monopolar rectangular pulses at 6 Hz for 3 s) was delivered by a constant-current device. During electrical stimulation mice were manually restrained and released into the observation cage immediately after current application. At the time of drug administration a drop of 0.5% tetracaine (Altacaine sterile solution, Altaire Pharmaceuticals Inc., USA) was applied into the eyes of all animals. Prior to the placement of corneal electrodes, a drop of 0.9% saline was applied on the eyes. In this model, seizures manifest in ‘stunned’ posture associated with rearing, forelimb automatic movements and clonus, twitching of the vibrissae and Straub-tail. At the end of the seizure episode the animals resume their normal exploratory behavior. In this test protection against a seizure episode is considered as the end point and animals are considered to be protected if they resume their normal exploratory behavior within 10 s after electrical stimulation.

#### Pilocarpine-induced seizures

In this test the mice were pretreated with the investigated compound or vehicle and 60 min later they received pilocarpine (400 mg/kg, ip). To avoid cholinergic side effects: peripheral toxicity and diarrhea, masticatory and stereotyped movements, animals treated with pilocarpine also received scopolamine butylbromide (1 mg/kg, ip) which was injected 45 min before pilocarpine. After the administration of the convulsant, the mice were observed during the next 60 min for behavioral changes. Latency to the onset of *status epilepticus* was considered as the endpoint in this test.^66^

#### Rotarod test

The test was performed according to the method recently described.^45^ The mice were trained daily for 3 consecutive days on the rotarod apparatus (Rotarod apparatus, May Commat RR0711, Turkey; rod diameter: 2 cm) rotating at a constant speed of 18 and rotations per minute (rpm). During each training session, the animals were placed on a rotating rod for 3 min with an unlimited number of trials. The proper experimentation was conducted 24 h after the final training trial. Briefly, 60 min before the rotarod test the mice were ip pretreated with the test compound (10, 30 and 60 mg/kg) and then, they were tested on the rotarod apparatus revolving at 6, 18 and 24 rpm. Motor impairments, defined as the inability to remain on the rotating rod for 1 min were measured and mean time spent on the rod was counted in each experimental group.

#### Grip strength test

The test enables to assess the effect of a drug on muscular strength. The grip-strength apparatus (TSE Systems, Germany) consists of a wire grid (8 × 8 cm) connected to an isometric force transducer (dynamometer). The mice were lifted by the tails so that their forepaws could grasp the grid. The mice were then gently pulled backward by the tail until the grid was released. The maximal force exerted by the mouse before losing grip was recorded. The mean of 3 measurements for each animal were averaged and subsequently the mean maximal force in each experimental group was determined.^67^

#### Effect on locomotor activity

The locomotor activity test was performed using activity cages (40 cm x 40 cm x 30 cm, supplied with I.R. beam emitters), (Activity Cage 7441, Ugo Basile, Italy) connected to a counter for the recording of light-beam interrupts. Sixty minutes before the experiment, the mice were pretreated with the test compound or vehicle, then being individually placed in the activity cages in a sound-attenuated room. The animals’ movements (i.e., the number of light-beam crossings) were counted during the next 30 min of the test.^45^

#### Four-plate test

The four-plate apparatus (Bioseb, France) consists of a cage (25 cm x 18 cm x 16 cm) that is floored with four rectangular metal plates (11 cm x 8 cm). The plates are separated from one another by a gap of 4 mm, and they are connected to an electroshock generator. In this test, after the habituation period (15 s), each mouse was subjected to an electric shock (intensity: 0.8 mA, duration: 0.5 s) when crossing from one plate to another (two limbs on one plate and two on another). The number of punished crossings was counted during 60 s. Agents with anxiolytic-like properties increase the number of punished passages in this assay.^68^

#### Elevated plus maze

The elevated plus maze for mice consists of two opposing open (30 cm x 5 cm) and two enclosed arms (30 cm x 5 cm x 25 cm) connected by a central platform forming the shape of a plus sign. The dimensions of the central field which connects the open and closed arms are 5 x 5 cm.

In this test, each mouse was individually placed at the central field of the apparatus with the head turned towards one of the closed arms. Animals’ behavior during 5 min was observed and recorded. In this test, the following parameters were measured in DDPM-3960-treated and control animals: number of entries in open arms, % entries in open arms, time spent in open arms, % time spent in open arms. To exclude the impact of excrements or smell left by a previous mouse on behavior of the next one, the device was cautiously cleaned after each testing session.^69^

#### Forced swim test

The experiment was carried out according to the method originally described by Porsolt et al. (1977).^70^ The mice were dropped individually into glass cylinders (height: 25 cm, diameter: 10 cm) filled with water to a height of 10 cm (maintained at 23-25 °C). In this assay, after an initial 2-min period of vigorous activity, each mouse assumes an immobile posture. The duration of immobility in experimental groups was recorded during the final 4 min of the total 6-min testing period. Mice were judged to be immobile when they remained floating passively in the water, making only small movements to keep their heads above the water surface.

#### Acute, thermally-induced pain (hot plate test)

The hot plate apparatus (Hot/cold plate, Bioseb, France) consists of electrically heated surface and is equipped with a temperature-controller that keeps the temperature constant at 55-56° C. In the hot plate test, the latency to pain reaction (hind paw licking or jumping) was measured as the indicative of nociception.^71^ In order to avoid paw tissue damage, a cut-off time of 60 s was established, and animals that did not respond within 60 s were removed from the hot plate apparatus and assigned a score of 60 s.

#### Neurogenic pain model (capsaicin test)

After an adaptation period (15 min), the mice received 1.6 μg of capsaicin dissolved in 20 μl of physiological saline and ethanol (5:1, v/v). Capsaicin was injected into the dorsal surface of the right hind paw of a mouse. In this assay, the animals were observed individually for 5 min following capsaicin injection. Pain-related behavior, i.e. the amount of time spent on licking, biting, flinching, or lifting the injected paw was measured using a chronometer.^72^

#### Tonic pain model (formalin test)

In rodents, the injection of diluted formalin solution evokes a biphasic nocifensive behavioral response (licking or biting the injected paw) of experimental animals. The first (acute) nociceptive phase of the test lasts for 5 min, whilst the second (late) one occurs between 15 and 30 min after formalin injection. In this assay 20 μl of 5% formalin solution was injected into the dorsal surface of the right hind paw of each mouse.^45^ Then, the mice were put separately into glass beakers and were observed for the next 30 min. The total time spent on licking or biting the formalin-injected paw was measured during the first 5 min of the test, and then between 15 and 30 min of the test in DDPM-3960-treated and vehicle-treated mice.

#### Data analysis

Data analysis of the results obtained in behavioral tests was provided by GraphPad Prism Software (ver. 8, CA, USA). The results were statistically evaluated using one-way analysis of variance (ANOVA), or repeated-measures ANOVA followed by Dunnett’s *post hoc* comparison and Student’s t-test. P < 0.05 was considered significant.

## Supporting information

Suppl_Inf

