## Supplementary material for "Synthesis and Biological Evaluation of Nipecotic Acid Derivatives with Terminally Double-Substituted Allenic Spacers as mGAT4 Inhibitors": Suppl_Inf

### Table of contents

| | | page |
|---|---|---|
| 1.1 | Synthesis of *rac*-**9** | S2 |
| 1.2 | Synthesis of ethyl esters of nipecotic acid derivatives of compound class *rac*-**10** | S3 |
| 1.3 | Experimental section | S4 |
| 2. | Analytical data of decomposition product **17** | S8 |
| 3. | Purity determination of DDPM-3960 [(*S*)-**9d**] | S9 |
| 4. | References | S10 |

### 1.1 Synthesis of *rac*-9

The synthesis of parent compound *rac*-**9** (Scheme 1), bearing a five-carbon atom *N*-substituent with a terminal allene unit has been accomplished by allenylation of the respective terminal alkyne (ATA reaction) of *rac*-**15** following a recently published procedure for the preparation of *rac*-**6**.[1] Accordingly, in the first step the propargylic amine *rac*-**21** (Scheme 1) was synthesized by reaction of terminal alkyne *rac*-**15** with paraformaldehyde (**19**) and allyl(*tert*-butyl)amine (**20**) in the presence of CuBr in a yield of 79% (Scheme 1). Subsequent treatment of *rac*-**21** with $ZnI_2$ led to the corresponding terminal allene *rac*-**22** in a yield of 69% (Scheme 1). Finally, after hydrolysis of the carboxylic acid ester function of the nipecotic acid ester derivative *rac*-**22** under basic conditions the desired nipecotic acid derived terminal allene *rac*-**9** was obtained. Parent compound *rac*-**9** appeared to be prone to side reactions under the conditions employed for the hydrolysis and the subsequent workup. Hence, this compound was considered as too labile for biological studies.

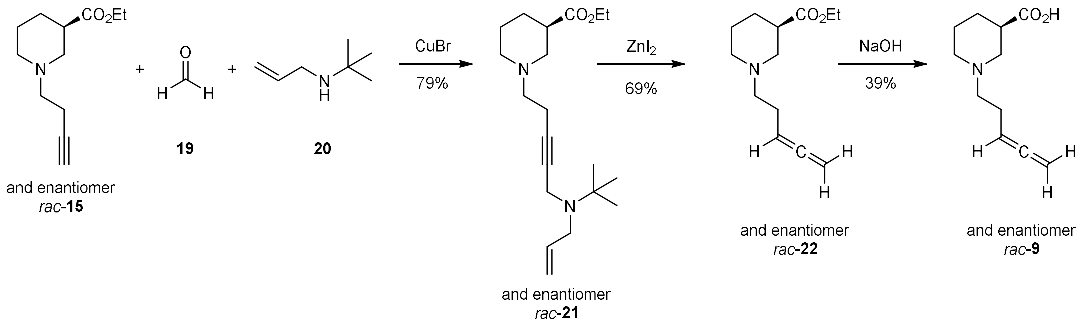


***Scheme 1.*** *Synthesis route to nipecotic acid derived parent compound rac-**9** with an allenic five-carbon atom comprising N-substituent.*

### 1.2 Synthesis of ethyl esters of nipecotic acid derivatives of compound class *rac*-10

For the syntheses of derivatives of parent compound *rac*-**9** exhibiting a five-carbon atom allenic spacer with one terminal aryl unit, i.e. of *rac*-1**0a**-**b** a method published by Wang et al. has been used. By this method based on a $Cu^I$ carbene migratory insertion mechanism terminal alkynes are directly transformed into the respective allene derivatives upon reaction with appropriate *N*-tosylhydrazones.[2,3] When this $Cu^I$-catalyzed cross-coupling reaction was performed with alkyne *rac*-**15** and the *N*-tosylhydrazone **23a** and **23b** delineated from benzaldehyde and from [1,1'-biphenyl]-2-carbaldehyde, respectively, the desired nipecotic acid derivatives *rac*-(*3R*,*R$_a$*)-**24a**/*rac*-(*3R*,*S$_a$*)-**24a** and *rac*-(*3R*,*R$_a$*)-**24b**/*rac*-(*3R*,*S$_a$*)-**24b** with a five-carbon atom allenic spacer were obtained in high yields (Scheme 2). In both cases, products represented ~ 1:1 mixtures of racemic diastereomers. Unfortunately, upon attempts to transform *rac*-(*3R*,*R$_a$*)-**24a**/*rac*-(*3R*,*S$_a$*)-**24a** and *rac*-(*3R*,*R$_a$*)-**24b**/*rac*-(*3R*,*S$_a$*)-**24b** into the free amino acids *rac*-(*3R*,*R$_a$*)-**10a**/*rac*-(*3R*,*S$_a$*)-**10a** and *rac*-(*3R*,*R$_a$*)-**10b**/*rac*-(*3R*,*S$_a$*)-**10b**, respectively, under basic or acidic conditions, no such product could be isolated. This behavior is in line with the low stability observed for parent compound *rac*-**9** (Scheme 1).

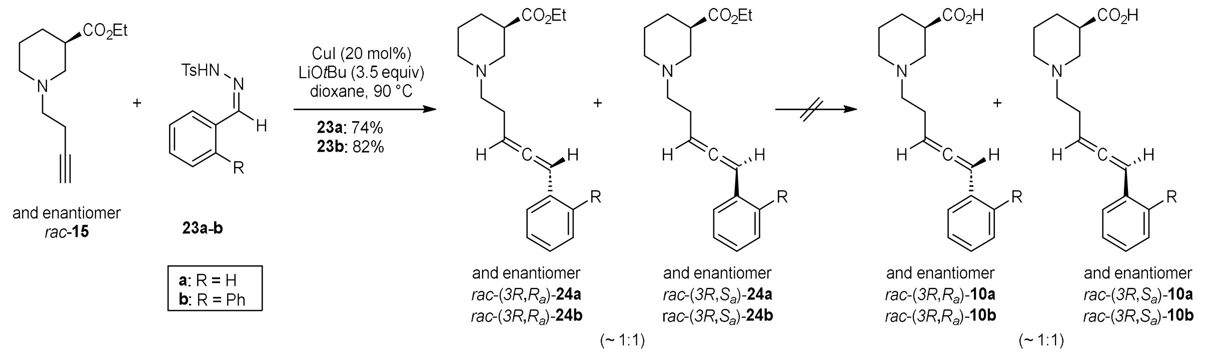


***Scheme 2.** Synthesis of nipecotic acid derivatives rac-(3R,R$_a$)-**24a**/rac-(3R,S$_a$)-**24a** and rac-(3R,R$_a$)-**24b**/rac-(3R,S$_a$)-**24b** with terminally mono-substituted five-carbon atom allenic spacer as ~ 1:1 mixture of racemic diastereomers by reaction of rac-**15** with N-tosylhydrazones **23a-b**.*

#### 1.3 Experimental section

General remarks:

General procedures (GP) refer to those given in the experimental part of the article. For compounds already presented in the article with a compound number, the same compound number is also used for these compounds in this section. With the highest compound number in the main article being **18**, all compounds newly introduced in this section are consecutively designated with compound numbers starting from **19**.

***rac*-[Ethyl 1-{5-[allyl(*tert*-butyl)amino]pent-3-yn-1-yl}piperidine-3-carboxylate] (*rac*-21):** This modified procedure is based on a procedure described by Yu et al.[4] To a Schlenk flask was added CuBr (65 mg, 0.45 mmol, 0.15 equiv) and molecular sieve (4Å). Toluene anhyd. (15 mL, 5.00 mL/mmol) was then added, followed by paraformaldehyde (171 mg, 5.40 mmol, 1.80 equiv), *N*-allyl-*N*-*tert*-butylamine (0.63 mL, 4.2 mmol, 1.4 equiv) and alkyne *rac*-**15** (628 mg, 3.00 mmol, 1.00 equiv). The reaction mixture was stirred at rt overnight. The completion of the reaction was monitored by TLC. The reaction mixture was then filtrated, washed with EtOAc and concentrated under vacuum. The crude product was purified by column chromatography (PE/EtOAc 1:1) to afford the desired propargylic amine *rac*-**21** as pale yellow oil (791 mg, 79%): $R_f$=0.26 (PE/EtOAc 1:1); $^1$H NMR (400 MHz, $CDCl_3$): $\delta$=1.16 (s, 9H, *t*-Bu), 1.25 (t, *J*=7.1 Hz, 3H, $OCH_2CH_3$), 1.42 (qd, *J*=11.8/3.8 Hz, 1H, $NCH_2CHCH_{ax}H_{eq}$), 1.49–1.64 (m, 1H, $NCH_2CH_{ax}H_{eq}CH_2CH$), 1.72 (dp, *J*=13.3/3.8 Hz, 1H, $NCH_2CH_{ax}H_{eq}CH_2CH$), 1.88–1.99 (m, 1H, $NCH_2CHCH_{ax}H_{eq}$), 2.05 (td, *J*=11.0/3.0 Hz, 1H, $NCH_{ax}H_{eq}CH_2CH_2CH$), 2.20 (t, *J*=10.7 Hz, 1H, $NCH_{ax}H_{eq}CHCH_2$), 2.31–2.41 (m, 2H, $NCH_2CCCH_2CH_2N$), 2.47–2.62 (m, 3H, $NCH_2CCCH_2CH_2N$ and $NCH_2CH_{ax}CH_2$), 2.78 (dt, *J*=10.5/4.0 Hz, 1H, $NCH_{ax}H_{eq}CH_2CH_2CH$), 2.94–3.04 (m, 1H, $NCH_{ax}H_{eq}CHCH_2$), 3.27 (dt, *J*=6.5/1.3 Hz, 2H, $NCH_2CH{=}CH_2$), 3.44 (t, *J*=2.2 Hz, 2H, $NCH_2CCCH_2CH_2N$), 4.13 (q, *J*=7.1 Hz, 2H, $OCH_2CH_3$), 5.09 (ddt, *J*=10.0/2.2/1.2 Hz, 1H, $NCH_2CH{=}CH_{trans}H_{cis}$), 5.23 (ddt, *J*=17.1/2.2/1.4 Hz, 1H, $NCH_2CH{=}CH_{trans}H_{cis}$), 5.82 (ddt, *J*=17.1/10.0/6.5 Hz, 1H, $NCH_2CH{=}CH_2$); $^{13}$C NMR (101 MHz, $CDCl_3$): $\delta$=14.2 (1C, $OCH_2CH_3$), 17.2 (1C, $NCH_2CCCH_2CH_2N$), 24.6 (1C, $NCH_2CHCH_2CH_2CH_2$), 26.9 (1C, $NCH_2CHCH_2CH_2CH_2$), 27.7 (3C, $NC(CH_3)_3$), 36.6 (1C, $NCH_2CCCH_2CH_2N$), 41.9 (1C, $NCH_2CHCH_2CH_2CH_2$), 50.0 (1C, $NCH_2CH{=}CH_2$), 53.4 (1C, $NCH_2CHCH_2CH_2CH_2$), 54.9 (1C, $NC(CH_3)_3$), 55.1 (1C, $NCH_2CHCH_2CH_2CH_2$), 57.7 (1C, $NCH_2CCCH_2CH_2N$), 60.3 (1C, $OCH_2CH_3$), 79.0 (1C, $NCH_2CCCH_2CH_2N$), 82.2 (1C, $NCH_2CCCH_2CH_2N$), 116.7 (1C, $NCH_2CH{=}CH_2$), 137.5 (1C, $NCH_2CH{=}CH_2$), 174.1 (1C, *C*O); IR (film): ṽ=3416, 2971, 2814, 2359, 2337, 1732, 1641, 1467, 1445, 1390, 1364, 1309, 1270, 1203, 1179, 1153, 1133, 1101, 1032, 994, 916 $cm^{-1}$; HRMS-ESI *m/z* $[M+H]^+$ calcd for $C_{20}H_{34}N_2O_2$: 335.2699, found: 335.2698.

***rac*-[Ethyl 1-(penta-3,4-dien-1-yl)piperidine-3-carboxylate] (*rac*-22):** To a Schlenk tube $ZnI_2$ (107 mg, 0.336 mmol, 0.800 equiv) was added. The $ZnI_2$ was then heated with a heat gun under vacuum until the pale yellow solid turned to darker yellow. Chlorobenzene anhyd. (8.0 mL/mmol) was added, followed by propargylic amine *rac*-**21** (140 mg, 0.420 mmol, 1.00 equiv). The reaction mixture was stirred at 100 °C for 30 min and the reaction was monitored by TLC. After cooling to rt, the reaction mixture was directly purified by column chromatography (PE/EtOAc 1:1) to afford the desired allene *rac*-**22** as pale yellow oil (64.4 mg, 69%): $R_f$=0.35 (PE/EtOAc 1:1); $^1$H NMR (500

MHz, $CDCl_3$): δ=1.25 (t, *J*=7.1 Hz, 3H, $OCH_2CH_3$), 1.44 (qd, *J*=11.4/4.2 Hz, 1H, $NCH_2CHCH_{ax}H_{eq}$), 1.53–1.66 (m, 1H, $NCH_2CH_{ax}H_{eq}CH_2CH$), 1.73 (dp, *J*=13.4/3.9 Hz, 1H, $NCH_2CH_{ax}H_{eq}CH_2CH$), 1.90–1.99 (m, 1H, $NCH_2CHCH_{ax}H_{eq}$), 2.02 (td, *J*=11.1/3.0 Hz, 1H, $NCH_{ax}H_{eq}CH_2CH_2CH$), 2.14–2.25 (m, 3H, $NCH_2CH_2CHCCH_2$ and $NCH_{ax}H_{eq}CHCH_2$), 2.43–2.50 (m, 2H, $NCH_2CH_2CHCCH_2$), 2.52–2.62 (m, 1H, $NCH_2CH_{ax}CH_2$), 2.76–2.82 (m, 1H, $NCH_{ax}H_{eq}CH_2CH_2CH$), 3.01 ($d_{br}$, *J*=10.6 Hz, 1H, $NCH_{ax}H_{eq}CHCH_2$), 4.13 (q, *J*=7.1 Hz, 2H, $OCH_2CH_3$), 4.67 (dt, *J*=6.7/3.2 Hz, 2H, $NCH_2CH_2CHCCH_2$), 5.11 (p, *J*=6.7 Hz, 1H, $NCH_2CH_2CHCCH_2$); $^{13}C$ NMR (126 MHz, $CDCl_3$): δ=14.2 (1C, $OCH_2CH_3$), 24.6 (1C, $NCH_2CHCH_2CH_2$), 25.7 (1C, $NCH_2CH_2CHCCH_2$), 27.0 (1C, $NCH_2CHCH_2CH_2CH_2$), 41.9 (1C, $NCH_2CHCH_2CH_2CH_2$), 53.7 (1C, $NCH_2CHCH_2CH_2CH_2$), 55.4 (1C, $NCH_2CHCH_2CH_2CH_2$), 58.1 (1C, $NCH_2CH_2CHCCH_2$), 60.3 (1C, $OCH_2CH_3$), 75.0 (1C, $NCH_2CH_2CHCCH_2$), 87.9 (1C, $NCH_2CH_2CHCCH_2$), 174.2 (1C, CO), 208.6 (1C, $CHCCH_2$); IR (film): ṽ=3425, 2942, 2856, 2805, 1956, 1732, 1641, 1468, 1444, 1371, 1311, 1273, 1211, 1180, 1153, 1101, 1031, 844 $cm^{-1}$; HRMS-ESI *m/z* $[M+H]^+$ calcd for $C_{13}H_{21}NO_2$: 224.1651, found: 224.1645.

***rac*-[1-(Penta-3,4-dien-1-yl)piperidine-3-carboxylic acid] (*rac*-9):** GP6 was followed using nipecotic acid ester *rac*-**22** (0.30 mmol, 67 mg), EtOH (1.0 mL) and 2M NaOH (2 equiv, 0.30 mL) for 2.75 h. The crude compound was purified by RP-MPLC (MeOH/$H_2O$ 3:7) to obtain the free amino acid *rac*-**9** as colorless viscous oil (23 mg, 39%): $^1H$ NMR (500 MHz, MeOD, NaOD): δ=1.35 (qd, *J*=12.8/4.1 Hz, 1H, $NCH_2CHCH_{ax}H_{eq}$), 1.59 (qt, *J*=12.8/3.9 Hz, 1H, $NCH_2CH_{ax}H_{eq}CH_2CH$), 1.67–1.75 (m, 1H, $NCH_2CH_{ax}H_{eq}CH_2CH$), 1.94 (td, *J*=11.8/2.8 Hz, 1H, $NCH_{ax}H_{eq}CH_2CH_2CH$), 1.96–2.01 (m, 1H, $NCH_2CHCH_{ax}H_{eq}$), 2.04 (t, *J*=11.4 Hz, 1H, $NCH_{ax}H_{eq}CHCH_2$), 2.15–2.28 (m, 2H, $CH_2CCHCH_2CH_2N$), 2.38 (tt, *J*=11.8/3.8 Hz, 1H, $NCH_2CH_{ax}CH_2$), 2.42–2.49 (m, 2H, $CH_2CCHCH_2CH_2N$), 2.89 ($d_{br}$, *J*=10.4 Hz, 1H, $NCH_{ax}H_{eq}CH_2CH_2CH$), 3.12 (ddt, *J*=11.3/3.5/1.6 Hz, 1H, $NCH_{ax}H_{eq}CHCH_2$), 4.68 (dt, *J*=6.6/3.2 Hz, 2H, $NCH_2CH_2CHCCH_2$), 5.12 (p, *J*=6.8 Hz, 1H, $NCH_2CH_2CHCCH_2$); $^{13}C$ NMR (126 MHz, MeOD, NaOD): δ=25.9 (1C, $NCH_2CH_2CH_2CH$), 26.3 (1C, $NCH_2CH_2CHCCH_2$), 29.4 (1C, $NCH_2CH_2CH_2CH$), 46.3 (1C, $NCH_2CH_2CH_2CH$), 54.9 (1C, $NCH_2CH_2CH_2CH$), 58.1 (1C, $NCH_2CHCH_2$), 59.5 (1C, $NCH_2CH_2CHCCH_2$), 75.3 (1C, $NCH_2CH_2CHCCH_2$), 88.4 (1C, $NCH_2CH_2CHCCH_2$), 182.8 (1C, CO), 210.1 (1C, $CH_2CHCCH_2$); IR (film): ṽ=3385, 2956, 1956, 1585, 1450, 1391, 1150, 1076, 857 $cm^{-1}$; HRMS-ESI *m/z* $[M+H]^+$ calcd for $C_{11}H_{17}NO_2$: 196.1338, found: 196.1331.

***rac*-[Ethyl ($R_a$)-1-(5-phenylpenta-3,4-dien-1-yl)-(3*R*)-piperidine-3-carboxylate] [*rac*-(3*R*,$R_a$)-24a] and *rac*-[ethyl ($S_a$)-1-(5-phenylpenta-3,4-dien-1-yl)-(3*R*)-piperidine-3-carboxylate] [*rac*-(3*R*,$S_a$)-24a]:** GP4 was followed applying alkyne *rac*-**15** (84 mg, 0.40 mmol), CuI (15 mg, 0.080 mmol), Li*t*OBu (112 mg, 1.40 mmol), and *N'*-benzylidene-4-methylbenzenesulfonohydrazide (**23a**, 241 mg, 0.880 mmol) in 1,4-dioxane (5 mL). The solution was stirred at 90 °C for 40 min. Purification by column chromatography (DCM/MeOH 98:2) afforded the desired allenes *rac*-(3*R*,$R_a$)-**24a** and *rac*-(3*R*,$S_a$)-**24a** as ~ 1:1 mixture of racemic diastereomers as yellow viscous oil (88.7 mg, 74%): $R_f$=0.20 (PE/EtOAc 8:2); $^1H$ NMR (500 MHz, $CDCl_3$): δ=1.19–1.30 (m, 3H, $OCH_2CH_3$), 1.37–1.50 (m, 1H, $NCH_2CHCH_{ax}H_{eq}$), 1.49–1.64 (m, 1H, $NCH_2CH_{ax}H_{eq}CH_2CH$), 1.66–1.77 (m, 1H, $NCH_2CH_{ax}H_{eq}CH_2CH$), 1.91–1.95 (m, 1H, $NCH_2CHCH_{ax}H_{eq}$), 1.98–2.05 (m, 1H, $NCH_{ax}H_{eq}CH_2CH_2CH$), 2.15–2.20 (m, 1H,

NC*H*$_{ax}$H$_{eq}$CHCH$_2$), 2.28–2.37 (m, 2H, NCH$_2$C*H*$_2$CHCCH), 2.46–2.63 (m, 3H, NC*H*$_2$CH$_2$CHCCH and NCH$_2$C*H*$_{ax}$CH$_2$), 2.77–2.80 (m, 1H, NCH$_{ax}$*H*$_{eq}$CH$_2$CH$_2$CH), 2.96–3.06 (m, 1H, NCH$_{ax}$*H*$_{eq}$CHCH$_2$), 4.07–4.16 (m, 2H, OC*H*$_2$CH$_3$), 5.58 (q, *J*=6.6 Hz, 1H, NCH$_2$CH$_2$C*H*CCH), 6.13 (dt, *J*=6.2/3.0 Hz, 1H, NCH$_2$CH$_2$CHCC*H*), 7.14–7.22 (m, 1H, ArH), 7.23–7.36 (m, 4H, ArH); $^{13}$C NMR (126 MHz, CDCl$_3$): *δ*=14.2 (1C, OCH$_2$*C*H$_3$), 24.6 (1C, NCH$_2$CHCH$_2$*C*H$_2$CH$_2$), 26.3 (0.5C, NCH$_2$*C*H$_2$CHCCH, dia1 or dia2), 26.3 (0.5C, NCH$_2$*C*H$_2$CHCCH, dia1 or dia2), 27.0 (0.5C, NCH$_2$CH*C*H$_2$CH$_2$CH$_2$, dia1 or dia2), 27.0 (0.5C, NCH$_2$CH*C*H$_2$CH$_2$CH$_2$, dia1 or dia2), 41.9 (0.5C, NCH$_2$*C*HCH$_2$CH$_2$CH$_2$, dia1 or dia2), 42.0 (0.5C, NCH$_2$*C*HCH$_2$CH$_2$CH$_2$, dia1 or dia2), 53.7 (0.5C, NCH$_2$CHCH$_2$CH$_2$*C*H$_2$, dia1 or dia2), 53.8 (0.5C, NCH$_2$CHCH$_2$CH$_2$*C*H$_2$, dia1 or dia2), 55.4 (0.5C, N*C*H$_2$CHCH$_2$CH$_2$CH$_2$, dia1 or dia2), 55.5 (0.5C, N*C*H$_2$CHCH$_2$CH$_2$CH$_2$, dia1 or dia2), 58.0 (1C, N*C*H$_2$CH$_2$CHCCH), 60.3 (0.5C, O*C*H$_2$CH$_3$, dia1 or dia2), 60.3 (0.5C, O*C*H$_2$CH$_3$, dia1 or dia2), 92.9 (1C, NCH$_2$CH$_2$*C*HCCH), 94.9 (0.5C, NCH$_2$CH$_2$CHC*C*H, dia1 or dia2), 94.9 (0.5C, NCH$_2$CH$_2$CHC*C*H, dia1 or dia2), 126.6 (2C, ArC), 126.7 (1C, ArC), 128.5 (2C, ArC), 134.8 (0.5C, ArC$_q$, dia1 or dia2), 134.8 (0.5C, ArC$_q$, dia1 or dia2), 174.2 (0.5C, *C*O, dia1 or dia2), 174.2 (0.5C, *C*O, dia1 or dia2), 205.3 (0.5C, NCH$_2$CH$_2$CH*C*CH, dia1 or dia2), 205.3 (0.5C, NCH$_2$CH$_2$CH*C*CH, dia1 or dia2); IR (film): ṽ=3061, 3030, 2936, 2855, 2807, 1948, 1730, 1598, 1495, 1459, 1370, 1300, 1273, 1210, 1179, 1151, 1134, 1100, 1072, 1029, 962, 911, 874, 776, 722, 692, 650, 629 cm$^{-1}$; HRMS-ESI *m/z* [*M*+H]$^+$ calcd for $C_{19}H_{25}NO_2$: 300.1964, found: 300.1956.

***rac*-{Ethyl (*R*$_a$)-1-[5-([1,1'-biphenyl]-2-yl)penta-3,4-dien-1-yl]-(*3R*)-piperidine-3-carboxylate} [*rac*-(*3R*,*R*$_a$)-24b] and *rac*-{ethyl (*S*$_a$)-1-[5-([1,1'-biphenyl]-2-yl)penta-3,4-dien-1-yl]-(*3R*)-piperidine-3-carboxylate} [*rac*-(*3R*,*S*$_a$)-24b]:** GP4 was followed applying alkyne *rac*-**15** (84 mg, 0.40 mmol), CuI (15 mg, 0.08 mmol), Li*t*OBu (112 mg, 1.40 mmol), and *N'*-([1,1'-biphenyl]-2-ylmethylene)-4-methylbenzenesulfonohydrazide (**23b**, 308 mg, 0.880 mmol) in 1,4-dioxane (5 mL). The solution was stirred at 90 °C for 40 min. Purification by column chromatography (PE/EtOAc 8:2) afforded the desired allenes *rac*-(*3R*,*R*$_a$)-**24b** and *rac*-(*3R*,*S*$_a$)-**24b** as ~ 1:1 mixture of racemic diastereomers as yellow viscous oil (122 mg, 82%): *R*$_f$=0.36 (PE/EtOAc 7:3); $^1$H NMR (500 MHz, CD$_2$Cl$_2$): *δ*=1.21 (t, *J*=7.1 Hz, 1.5H, OCH$_2$C*H*$_3$, dia1 or dia2), 1.22 (t, *J*=7.1 Hz, 1.5H, OCH$_2$C*H*$_3$, dia1 or dia2), 1.34–1.47 (m, 1H, NCH$_2$CHC*H*$_{ax}$H$_{eq}$), 1.47–1.59 (m, 1H, NCH$_2$C*H*$_{ax}$H$_{eq}$CH$_2$CH), 1.60–1.78 (m, 1H, NCH$_2$CH$_{ax}$*H*$_{eq}$CH$_2$CH), 1.82–1.93 (m, 1H, NCH$_2$CHCH$_{ax}$*H*$_{eq}$), 1.99 (td, *J*=11.3/3.0 Hz, 0.5H, NC*H*$_{ax}$H$_{eq}$CH$_2$CH$_2$CH, dia1 or dia2), 2.02 (td, *J*=11.3/3.0 Hz, 0.5H, NC*H*$_{ax}$H$_{eq}$CH$_2$CH$_2$CH, dia1 or dia2), 2.16 (t, *J*=10.3 Hz, 1H, NC*H*$_{ax}$H$_{eq}$CHCH$_2$), 2.26 (q, *J*=6.9 Hz, 1H, NCH$_2$C*H*$_2$CHCCH, dia1 or dia2), 2.27 (q, *J*=6.9 Hz, 1H, NCH$_2$C*H*$_2$CHCCH, dia1 or dia2), 2.40–2.57 (m, 3H, NC*H*$_2$CH$_2$CHCCH and NCH$_2$C*H*$_{ax}$CH$_2$), 2.62–2.82 (m, 1H, NCH$_{ax}$*H*$_{eq}$CH$_2$CH$_2$CH), 2.93 (d$_{br}$, *J*=10.6 Hz, 0.5H, NCH$_{ax}$*H*$_{eq}$CHCH$_2$, dia1 or dia2), 2.96 (d$_{br}$, *J*=10.6 Hz, 0.5H, NCH$_{ax}$*H*$_{eq}$CHCH$_2$, dia1 or dia2), 4.07 (q, *J*=7.1 Hz, 1H, OC*H*$_2$CH$_3$, dia1 or dia2), 4.08 (q, *J*=7.1 Hz, 1H, OC*H*$_2$CH$_3$, dia1 or dia2), 5.53 (q, *J*=6.7 Hz, 1H, NCH$_2$CH$_2$C*H*CCH), 6.15 (dt, *J*=6.2/3.0 Hz, 1H, NCH$_2$CH$_2$CHCC*H*), 7.19–7.26 (m, 2H, ArH), 7.26–7.32 (m, 1H, ArH), 7.32–7.38 (m, 3H, ArH), 7.38–7.45 (m, 2H, ArH), 7.57 (dd, *J*=8.0/0.8 Hz, 1H, ArH); $^{13}$C NMR (126 MHz, CD$_2$Cl$_2$): *δ*=14.4 (1C, OCH$_2$*C*H$_3$), 25.1 (1C, NCH$_2$CHCH$_2$*C*H$_2$CH$_2$), 26.8 (0.5C, NCH$_2$*C*H$_2$CHCCH, dia1 or dia2), 26.8 (0.5C, NCH$_2$*C*H$_2$CHCCH, dia1 or dia2), 27.4 (0.5C, NCH$_2$CH*C*H$_2$CH$_2$CH$_2$, dia1 or dia2), 27.4 (0.5C, NCH$_2$CH*C*H$_2$CH$_2$CH$_2$, dia1 or dia2), 42.3 (0.5C, NCH$_2$*C*HCH$_2$CH$_2$CH$_2$, dia1 or dia2), 42.4 (0.5C, $NCH_2\mathit{C}HCH_2CH_2CH_2$, dia1 or dia2), 54.1 (1C, $NCH_2CHCH_2CH_2\mathit{C}H_2$), 55.9 (0.5C, $N\mathit{C}H_2CHCH_2CH_2CH_2$, dia1 or dia2), 56.0 (0.5C, $N\mathit{C}H_2CHCH_2CH_2CH_2$, dia1 or dia2), 58.4 (0.5C, $N\mathit{C}H_2CH_2CHCCH$, dia1 or dia2), 58.4 (0.5C, $N\mathit{C}H_2CH_2CHCCH$, dia1 or dia2), 60.5 (0.5C, $O\mathit{C}H_2CH_3$, dia1 or dia2), 60.5 (0.5C, $O\mathit{C}H_2CH_3$, dia1 or dia2), 93.0 (0.5C, $NCH_2CH_2\mathit{C}HCCH$ or $NCH_2CH_2CHC\mathit{C}H$, dia1 or dia2), 93.0 (0.5C, $NCH_2CH_2\mathit{C}HCCH$ or $NCH_2CH_2CHC\mathit{C}H$, dia1 or dia2), 93.1 (0.5C, $NCH_2CH_2CHC\mathit{C}H$ or $NCH_2CH_2\mathit{C}HCCH$, dia1 or dia2), 93.1 (0.5C, $NCH_2CH_2CHC\mathit{C}H$ or $NCH_2CH_2\mathit{C}HCCH$, dia1 or dia2), 126.9 (1C, ArC), 127.4 (1C, ArC), 127.7 (0.5C, ArC, dia1 or dia2), 127.7 (0.5C, ArC, dia1 or dia2), 127.8 (0.5C, ArC, dia1 or dia2), 127.8 (0.5C, ArC, dia1 or dia2), 128.5 (2C, ArC), 130.1 (2C, ArC), 130.5 (0.5C, ArC, dia1 or dia2), 130.5 (0.5C, ArC, dia1 or dia2), 132.6 (0.5C, $ArC_q$, dia1 or dia2), 132.6 (0.5C, $ArC_q$, dia1 or dia2), 140.5 (0.5C, $ArC_q$, dia1 or dia2), 140.5 (0.5C, $ArC_q$, dia1 or dia2), 141.3 (0.5C, $ArC_q$, dia1 or dia2), 141.3 (0.5C, $ArC_q$, dia1 or dia2), 174.4 (0.5C, *C*O, dia1 or dia2), 174.4 (0.5C, *C*O, dia1 or dia2), 206.1 (1C, $NCH_2CH_2CH\mathit{C}CH$); IR (film): $\tilde{\nu}$=3058, 3026, 2940, 2854, 2804, 1947, 1730, 1596, 1499, 1480, 1467, 1436, 1370, 1304, 1210, 1179, 1152, 1102, 1032, 1009, 881, 770, 746, 702 $cm^{-1}$; HRMS-ESI *m/z* $[M+H]^+$ calcd for $C_{25}H_{29}NO_2$: 376.2277, found: 376.2271.

### 2. Analytical data of decomposition product 17

$^{1}$H NMR spectrum of decomposition product mixture **17**

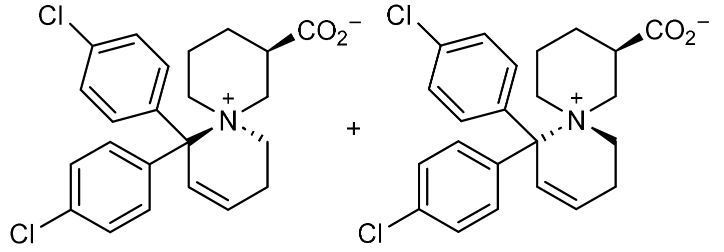

and enantiomer **17** and enantiomer

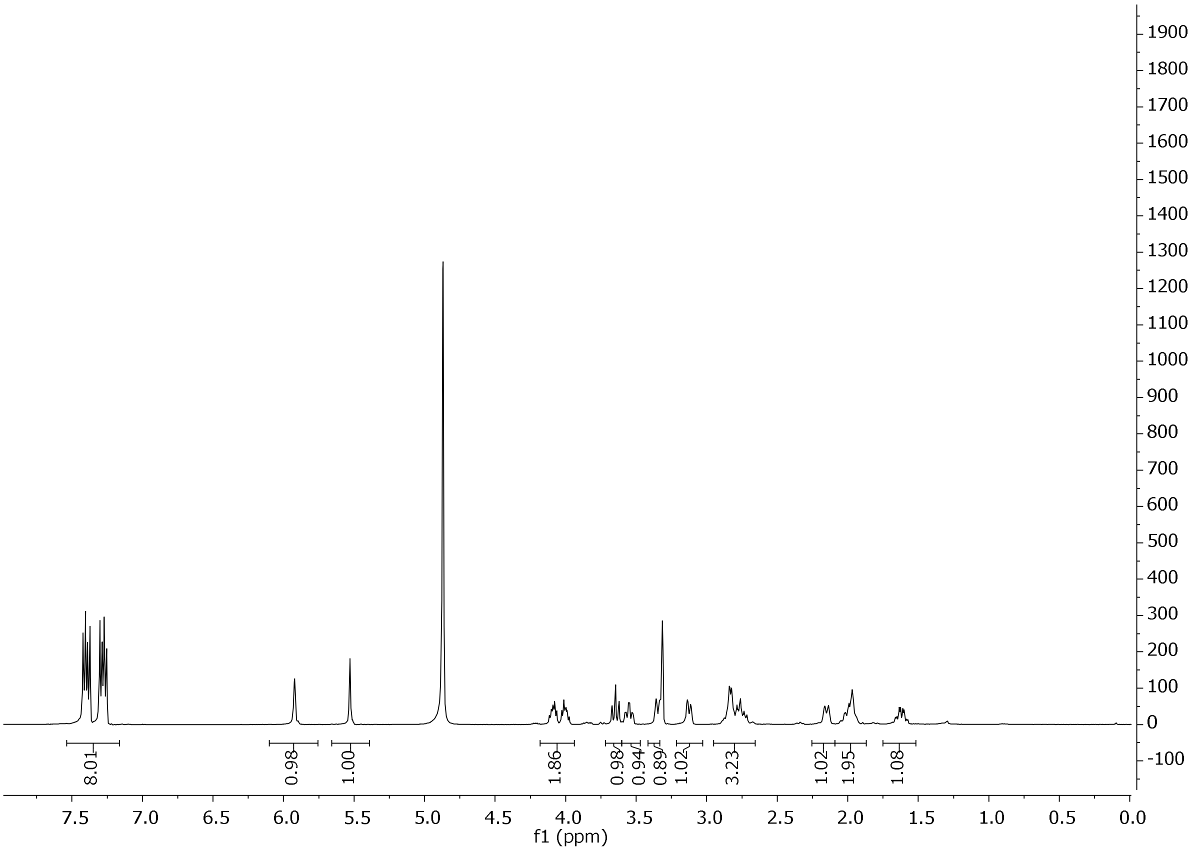


***SI-Figure 1.** $^{1}$H-NMR of compound mixture **17** in MeOD.*

Numerical listing of analytical data of decomposition product mixture **17**

$^{1}$H NMR (500 MHz, MeOD): $\delta$=1.62 (qd, $J$=12.9/4.8 Hz, 1H, C$H_2$), 1.87–2.08 (m, 2H, C$H_2$), 2.15 (d, $J$=12.5 Hz, 1H, C$H_2$), 2.64–2.92 (m, 3H, C$H_2$ and C$H$), 3.13 (d, $J$=11.7 Hz, 1H, C$H_2$), 3.33–3.39 (m, 1H, C$H_2$), 3.55 (td, $J$=12.3/4.3 Hz, 1H, C$H_2$), 3.65 (t, $J$=12.4 Hz, 1H, C$H_2$), 3.95–4.14 (m, 2H, C$H_2$), 5.53 (s, 1H, C$H$), 5.88–5.97 (m, 1H, C$H$), 7.21–7.35 (m, 4H, ArH), 7.35–7.48 (m, 4H, ArH); $^{13}$C NMR (126 MHz, MeOD): $\delta$=20.8 (1C, $C$H$_2$), 24.9 (1C, $C$H$_2$), 26.5 (1C, $C$H$_2$), 40.7 (1C, $C$H), 45.2 (1C, $C$H), 59.1 (1C, $C$H$_2$), 59.5 (1C, $C$H$_2$), 61.2 (1C, $C$H$_2$), 128.9 (2C, Ar$C$), 128.9 (2C, Ar$C$), 129.3 (1C, $C$H), 129.9 (2C, Ar$C$), 130.0 (2C, Ar$C$), 133.3 (1C, $C$), 133.4 (1C, $C$), 138.5 (1C, $C$), 138.8 (1C, $C$), 150.8 (1C, $C$), 176.2 (1C, $C$). The assignment was done by means of 2D NMR spectra (HMQC, COSY, DEPT). HRMS-ESI *m/z* $[M+H]^+$ calcd for $C_{23}H_{23}Cl_2NO_2$: 416.1184, found: 416.1172.

**4. Purity determination of DDPM-3960 [(*S*)-9d]**

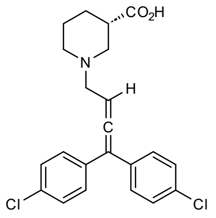


DDPM-3960
(*S*)-**9d**

Chemical Purity

Chemical purity was determined by means of analytical HPLC on an Agilent 1100 instrument (G1329A ALS autosampler, G1316A column compartment, G1314A VWD detector, G1312A binary pump, G1379A degasser), equipped with a Lichrospher 100 RP-18 (5 µm) in a LiChroCART 250-4 column, with elution at 0.5 mL/min with ammonium formate buffer (10 mM, pH 6.8) to MeOH 20:80.

Chemical purity amounted to ~ 100%.

HPLC trace

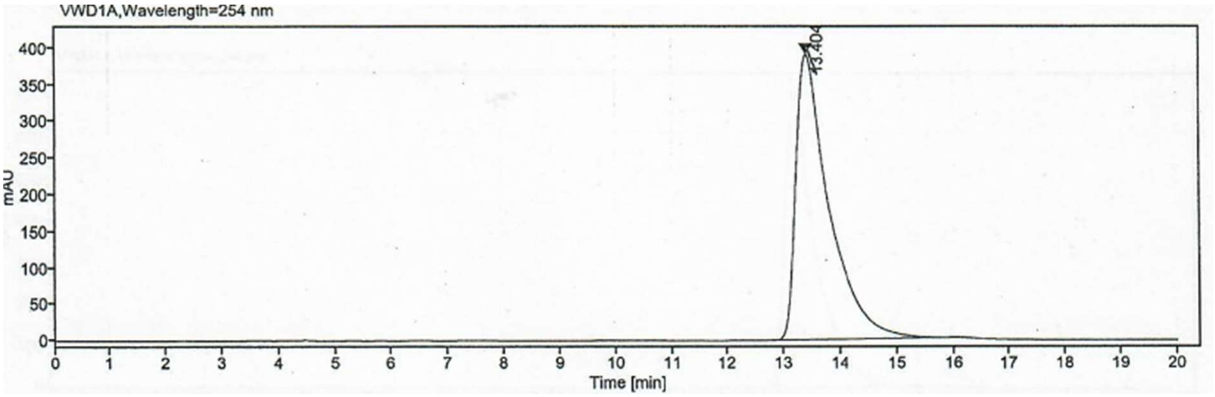


**Signal:** VWD1A,Wavelength=254 nm

| RT [min] | Width [min] | Area | Height | Area% | Name |
|---|---|---|---|---|---|
| 13.404 | 2.95 | 15910.20 | 389.67 | 100.00 | peak@13,404min |
| | Sum | 15910.20 | | | |

Enantiopurity

Enantiopurity was determined by means of chiral HPLC with a Chiralpak ZWIX(-) column and a solvent mixture of 40% MeOH, 60% ACN, 50 mM FA, 25 mM DEA as eluent at a flow rate of 0.5 mL/min.

The enantiopurity of (*S*)-**9d** amounted to 92.3% ee.

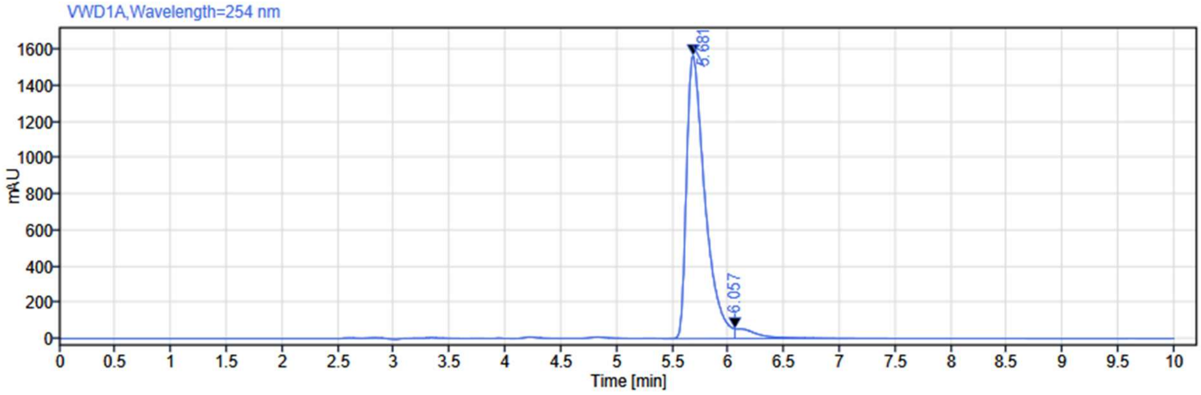


**Signal:** VWD1A,Wavelength=254 nm

| RT [min] | Width [min] | Area | Height | Area% |
|---|---|---|---|---|
| 5.681 | 0.17 | 17975.01 | 1567.12 | 96.14 |
| 6.057 | 0.23 | 722.46 | 52.58 | 3.86 |
